# A New Paralog Removal Pipeline Resolves Conflict between RAD-seq and Enrichment

**DOI:** 10.1101/2020.10.26.355248

**Authors:** Wenbin Zhou, John Soghigian, Qiu-yun (Jenny) Xiang

## Abstract

Target enrichment and RAD-seq are well-established high throughput sequencing technologies that have been increasingly used for phylogenomic studies, and the choice between methods is a practical issue for plant systematists studying the evolutionary histories of biodiversity of relatively recent origins. However, few studies have compared the congruence and conflict between results from the two methods within the same group of organisms, especially in plants, where extensive genome duplication events may complicate phylogenomic analyses. Unfortunately, currently widely used pipelines for target enrichment data analysis do not have a vigorous procedure for remove paralogs in Hyb-Seq data. In this study, we employed RAD-seq and Hyb-Seq of Angiosperm 353 genes in phylogenomic and biogeographic studies of *Hamamelis* (the witch-hazels) and *Castanea* (chestnuts), two classic examples exhibiting the well-known eastern Asian-eastern North American disjunct distribution. We compared these two methods side by side and developed a new pipeline (PPD) with a more vigorous removal of putative paralogs from Hyb-Seq data. The new pipeline considers both sequence similarity and heterozygous sites at each locus in identification of paralogous. We used our pipeline to construct robust datasets for comparison between methods and downstream analyses on the two genera. Our results demonstrated that the PPD identified many more putative paralogs than the popular method HybPiper. Comparisons of tree topologies and divergence times showed significant differences between data from HybPiper and data from our new PPD pipeline, likely due to the error signals from the paralogous genes undetected by HybPiper, but trimmed by PPD. We found that phylogenies and divergence times estimated from our RAD-seq and Hyb-Seq-PPD were largely congruent. We highlight the importance of removal paralogs in enrichment data, and discuss the merits of RAD-seq and Hyb-Seq. Finally, phylogenetic analyses of RAD-seq and Hyb-Seq resulted in well-resolved species relationships, and revealed ancient introgression in both genera. Biogeographic analyses including fossil data revealed a complicated history of each genus involving multiple intercontinental dispersals and local extinctions in areas outside of the taxa’s modern ranges in both the Paleogene and Neogene. Our study demonstrates the value of additional steps for filtering paralogous gene content from Angiosperm 353 data, such as our new PPD pipeline described in this study. [RAD-seq, Hyb-Seq, paralogs, *Castanea*, *Hamamelis*, eastern Asia-eastern North America disjunction, biogeography, ancient introgression]

## INTRODUCTION

High throughput sequencing (HTS) technologies, such as those associated with amplicon sequencing, restriction site digestion, target enrichment, and transcriptome sequencing, have empowered systematists and evolutionary biologists to infer phylogeny with genome-wide molecular markers for a better understanding of species relationships and to answer evolutionary questions with new perspectives that were not possible in the past (e.g. Landis et al. 2018; Lu et al. 2018; Dong et al. 2019; One Thousand Plant Transcriptomes Initiative 2019; and see reviews in Lemmon and Lemmon 2013; McCormack et al. 2013; Zimmer and Wen 2015; Chanderbali et al. 2017; Ma et al. 2017; Funk 2018; Lewin et al. 2018; Dodsworth et al. 2019). Among these HTS technologies, restriction site associated DNA sequencing (RAD-seq) and target enrichment (Hyb-Seq in plants or sequence capture and ultraconserved elements, UCEs, in animals) are highly promising and are increasingly used for phylogenomic study of lineages across different evolutionary phylogenetic timescales (e.g. Eaton and Ree 2013; Fernández-Mazuecos et al. 2018; Zhou et al. 2018; Du et al. 2019; MacGuigan and Near 2019 for RAD-seq data; Faircloth et al. 2013; McCormack et al. 2013; Leache et al. 2015; Léveillé-Bourret et al. 2018; Gaynor et al. 2020 for Target enrichment data).

RAD-seq and target enrichment methods each have their own strengths and weaknesses, which have been previously discussed (reviewed in Lemmon and Lemmon 2013; McCormack et al. 2013; Zimmer and Wen 2015; Harvey et al. 2016; Dodsworth et al. 2019). The main differences involve the portion of the genome captured, library preparations procedures, pipelines for data analyses, and cost. RAD-seq approaches are relatively easy and have a relatively lower cost for the laboratory work and provide deep coverage of numerous loci associated with the cutting sites of the restriction enzyme(s) employed. Furthermore, no prior knowledge of gene sequences is required (Peterson et al. 2012; Leache et al. 2015; Andrews et al. 2016; Harvey et al. 2016; Manthey et al. 2016; Dodsworth et al. 2019). In contrast, target enrichment methods cost more per sample and have relatively more intensive lab work, and the number of loci is generally limited to a targeted set of highly conserved genomic regions (and their flanking areas), often protein coding genes. Prior knowledge of target sequences is needed to design the probes of targeted genes, either from the organism of interest, or a related species. However, the target enrichment methods generally have greater repeatability between experiments and between labs if the same probes are used (Harvey et al. 2016), and typically generate relatively longer sequences at each locus. Therefore, data from target enrichment are a lasting and amplifiable resource for comparative studies at multiple taxonomic scales. Additionally, as target enrichment is often based on a known set of conserved regions or genes, researchers may also know better the function and genomic distribution of the genes targeted. Conversely, RAD-seq data have relatively low repeatability between experiments, labs, and taxa as a result of variation in DNA samples and experimental procedures and reagents (DaCosta and Sorenson 2014; Flanagan and Jones 2018; Malinsky et al. 2018). In addition, target enrichment data are suitable to phylogenomic studies of both deep and shallow phylogenetic divergence because the data contain both conserved coding sequences and their flanking variable sequences, whereas RAD-seq data is traditionally thought to be more appropriate for studies of shallow phylogenetic divergence (e.g., studies within a genus or species, and sometimes in relatively young families) due to allele dropout effects (Arnold et al. 2013; Gautier et al. 2013; Eaton 2014, and reviewed in Leaché and Oaks 2017). However, recent studies have shown that the allele-drop out (missing data) in RAD-seq data is usually systematic (or hierarchical) and phylogenetically informative (DaCosta and Sorenson 2016). Recent studies using RAD-seq have resolved phylogenies of lineages diverging during the Eocene and even the late Cretaceous (Eaton et al. 2016; Hou et al. 2016; Leaché and Oaks 2017; Lecaudey et al. 2018; Zhou et al. 2018; MacGuigan and Near 2019; Du et al. 2020; Mu et al. 2020).

For a phylogenetic study of a lineage at the phylogenetic timescales applicable to both target enrichment and RAD-seq approaches, e.g., building species level phylogenies, researchers may choose one of these approaches with the belief that either option would produce robust data for correct inference of phylogeny and divergence times. However, we have little knowledge about the level of congruence between the two data types and the impact of data analysis procedures. Few studies have made efforts to compare results derived from target enrichment and RAD-seq. At present, only a few studies on animals have applied both methods to allow a direct comparison of the phylogenetic utilities between the target enrichment data and RAD-seq data and to evaluate their congruences (Leache et al. 2015; Harvey et al. 2016; Manthey et al. 2016). Harvey et al. (2016) and Manthey et al. (2016) found that the topologies from RAD-seq and sequence capture were similar for lineages of birds that originated in the late Miocene or more recently, but with longer branch lengths for sequence capture. However, Leache et al. (2015), in an analysis of lizard lineage dating back to the Eocene, found differences in concatenated gene topologies and the species relationships between the target enrichment data and RAD-seq data. These findings are limited to only a few case studies, in which how the two datasets behave in subsequent phylogeny-based and branch length-dependent comparative analyses, such as divergence time dating and range evolution, was not assessed. To our knowledge, this is the first study comparing the phylogenetic outcomes between data from Hyb-Seq, the target enrichment approach developed for plant phylogenomics by Weitemier et al. (2014), and data from RAD-seq.

The development of the Angiosperm 353 kit (Johnson et al. 2019), which captures 353 low copy nuclear genes across angiosperms, has enabled phylogenomic studies across angiosperm lineages from family to genes (e.g., Gaynor et al. 2020 for Diapensaceae; Larridon et al. 2020 for Cyperaceae; Murphy et al. 2020 for *Nepenthes* in Nepenthaceae; Shee et al. 2020 for *Scheffera* in Araliaceae). An explosion of phylogenomic studies using the Angiosperm 353 probes is expected in the plant systematics community in the coming years. This endeavor will result in combinable datasets for building the “tree of life” of angiosperms through global-scale analysis (Dodsworth et al. 2019; Johnson et al. 2019). However, the universal probe kit has a disadvantage compared to taxon-specific kits in that the 353 genes targeted may or may not all be single copy across all species on which the kit is used, and probe binding affinity may cause probes to target unintended paralogous sequences (McCartney et al. 2016). In other words, the potential high divergence of some of the targeted 353 genes among the diverse angiosperm genomes poses a concern on possible prevalence of paralogs in the Hyb-Seq data. It is unknown if the current pipelines developed for analyses of target enrichment data have the ability to reliably exclude paralogous gene copies in the Hyb-Seq data derived from the Angiosperm 353 probe kit.

Orthologs are genes related by descent from a common ancestor (due to a speciation event) and their evolutionary history tracks the phylogeny of species, while paralogs are products of gene duplication events. Comparisons of paralogous copies of genes among species compromise phylogenetic inferences because the gene trees do not track speciation events, and hence, do not depict the true species relationships (Altenhoff et al. 2019; Fig. 1a). Therefore, excluding paralogous comparisons in phylogenetic studies of species relationships is pivotal, although paralogous gene sequences have value in other areas of comparative genomics (Madlung 2013; Limborg et al. 2016; McKinney et al. 2017). However, in Hyb-Seq data, orthologs and paralogs are often difficult to distinguish due to their high similarity in sequence identity (Altenhoff et al. 2019). Current pipelines for target enrichment data, including HybPiper (Johnson et al. 2016), PHYLUCE (Faircloth 2016), and SECAPR (Andermann et al. 2018a), all merely consider the sequence similarity in detecting paralogs (Figs. 1b and 1c; details in Supplementary Information, available on Dryad).

**Fig. 1.**
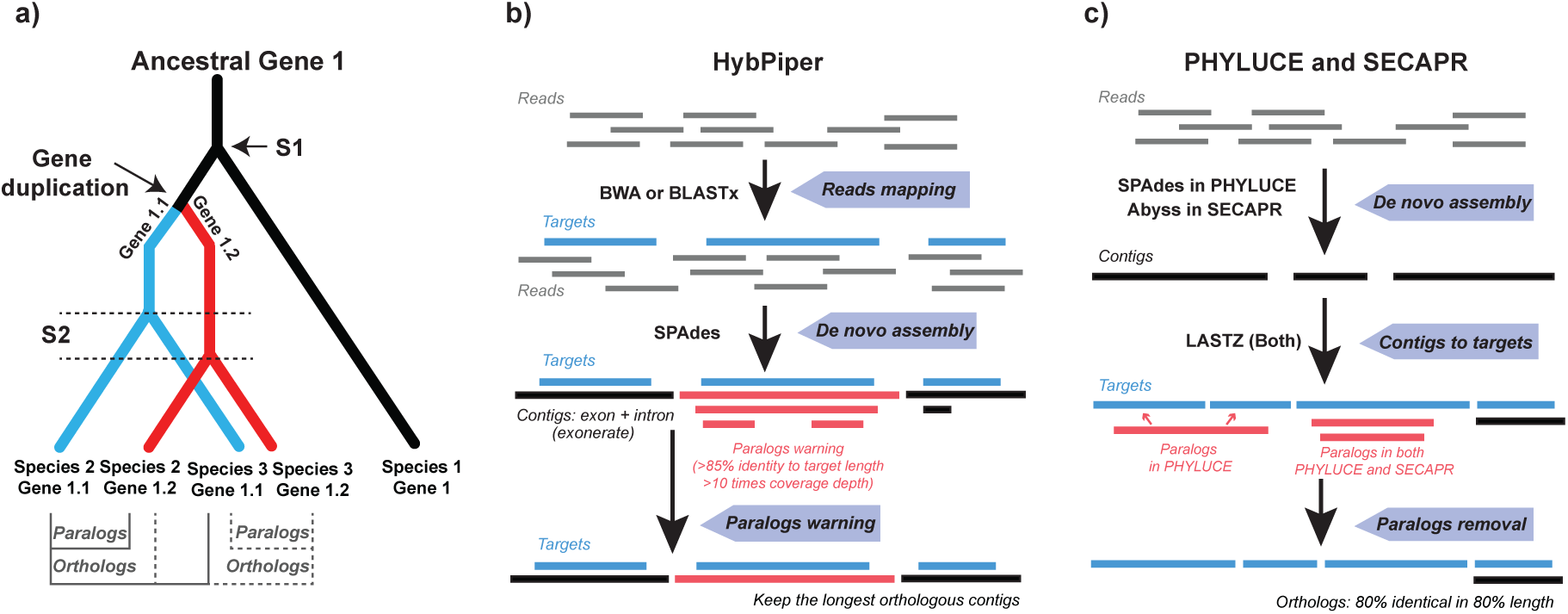
Concepts of ortholog-paralog and the key steps of workflow of current bioinformatics programs for target enrichment data analysis. a) Illustration of orthologs and paralogs. Orthologs are generated by speciation events, while paralogs are generated by gene duplication events. Speciation events are indicated by S1 and S2. b) The workflow of HybPiper (Johnson et al. 2016), including reads mapping using BLASTx or BWA (Li and Durbin 2010), de novo assembly of contigs using SPAdes (Bankevich et al. 2012) and paralogs (in red) detection. c) The workflow of PHYLUCE (Faircloth 2016) and SECAPR (Andermann et al. 2018a), including de novo assembly of contigs from reads using SPAdes (in PHYLUCE) and Abyss (Simpson et al. 2009) (in SECAPR), respectively, contigs mapping using LASTZ (Harris 2007), and detection of paralogs (in red), followed by removal of detected paralogs. All three methods detect putative paralogs by observation of more than one unique contigs in an individual mapped to the same reference gene with >85% (HybPiper) or >80% (PHYLUCE and SECAPR) sequence identity or the same contig mapped to more than one reference genes.

A sequence similarity-based approach for calling paralogous genes, such as those implemented in the three popular pipelines mentioned above, may be sufficient for phylogenetic studies using custom designed probes based on orthologous sequences encompassing the study group. However, for studies leveraging probes built from evolutionarily distant taxa from the focal group of investigation, especially in groups where gene and genome duplication are thought to be common such as plants, sequence similarity between contig and target genes alone may not be sufficient for removing all paralogs. Additional analyses of the sequence data may be needed to remove the potential paralogous sequences before performing phylogenetic analyses. In this study we propose a supplementary criterion to the sequence similarity for detecting and removing the problematic paralogous gene data from Hyb-Seq by examining heterozygous sites within and among individuals in the aligned sequences (Fig. 2). Low heterozygosity in a species-level dataset is expected under the assumption that polymorphisms among species are more likely to be fixed differences between paralogs over deep divergences (Eaton 2014). Even at the shallow level of phylogenetic divergence (e.g. population genetics), high heterozygosity within a locus may also be attributed to paralogs (Hohenlohe et al. 2011; Harvey et al. 2016; McKinney et al. 2017). Additionally, a high number of heterozygous sites of a locus within an individual can be used as an indicator for gene duplication events or previously undetected polyploidy (Medina et al. 2019). Most enrichment pipelines do not by default consider the sequence variants of contigs/loci (details in Supplementary Information, available on Dryad). In contrast, the methods developed for analyses of RAD-seq data are usually stricter and use more parameters to filter putative paralogs (Harvey et al. 2016). For example, PyRAD (Eaton 2014) not only uses the parameter of clustering based on sequence similarity within each locus, but also uses two other parameters to filter sequences with high heterozygous sites and with high percentage of shared heterozygous sites (Eaton 2014; McKinney et al. 2017). Therefore, we modified HybPiper (Fig. 2) to include analyses of the heterozygosity parameters for further filtering of putative paralogs from enrichment data.

**Fig. 2.**
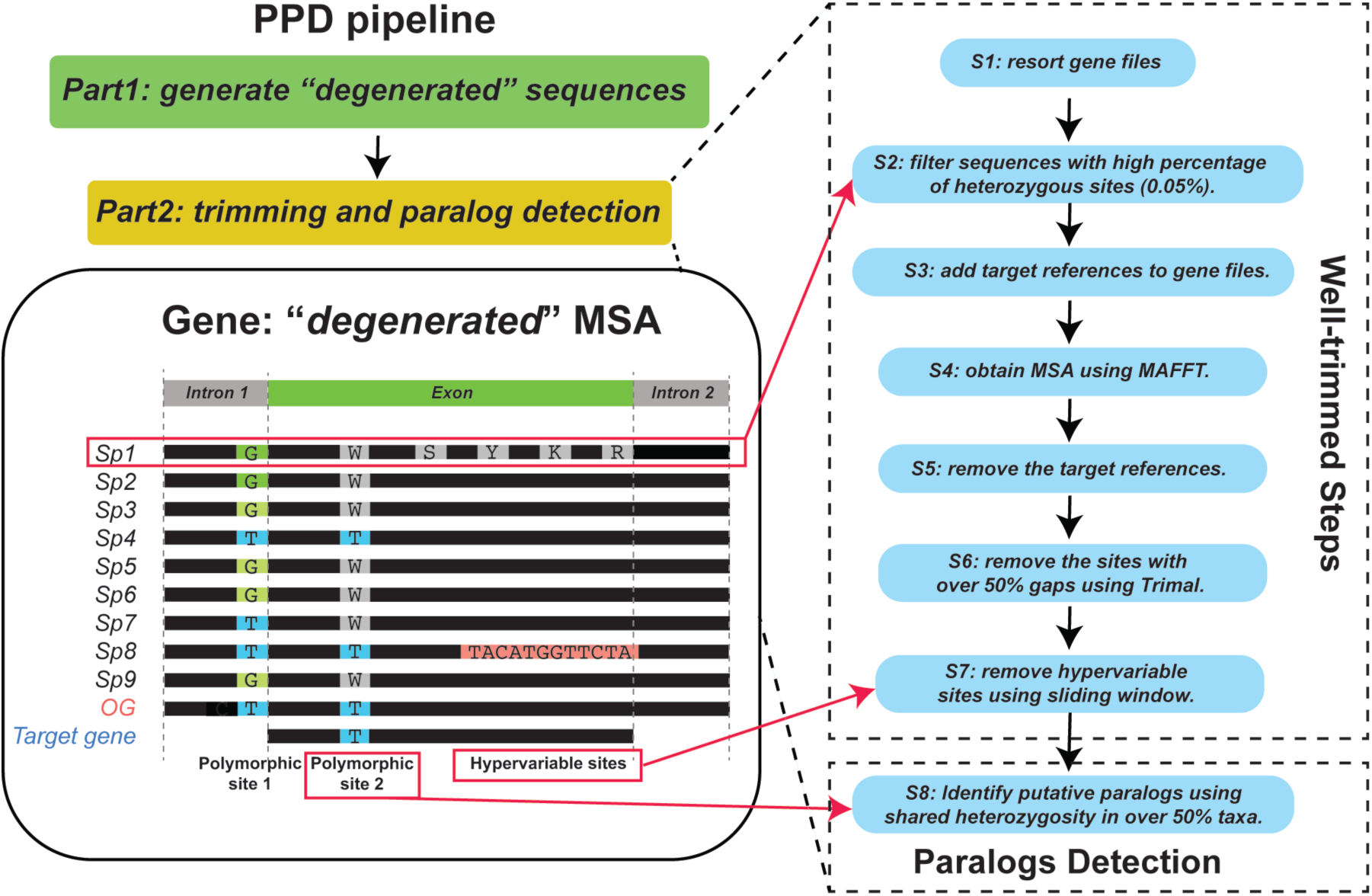
The putative paralogs detection (PPD) pipeline analytical workflow. The flowchart shows the basic PPD functions, including two major steps: first, generating a “*degenerated*” sequences; second, generating well-trimmed matrix and detecting paralogous genes. The “well-trimmed” process (s1 to s7) include filtering out sequences with high percentage of heterozygous sites (>5%), constructing Multiple Sequence Alignment (MSA), trimming MSA by removing sites missing in >50% samples (gaps) and hypervariable sites by a sliding window trimming method. The paralog detection step (s8) can detect additional paralogs indicated by presence of heterozygous site(s) present in >50% samples in the MSA. The “*degenerated*” sequence means the sequence were coded by IUPAC ambiguity codes. See details in Supplementary Information, available on Dryad. MSA: multiple sequence alignment; Sp: species; OG: Outgroup.

In order to compare Hyb-Seq and RAD-seq, as well as to evaluate how putative paralog detection impacted the results of a Hyb-Seq analysis, we conducted extensive analyses of Hyb-Seq data from the Angiosperm 353 kit and RAD-seq data from two genera: *Castanea* (Chestnuts of Fagaceae) and *Hamamelis* (Witch-hazel of Hamamelidaceae). From both genera, we inferred species relationships, divergence times, and biogeographic histories and compared the results to gain insights into the congruence between Angiosperm 353 and RAD-seq data. These two genera are common elements of the deciduous forests of the northern hemisphere and discontinuously distributed in eastern Asia (EA) and eastern North America (ENA); the study of which allow us to gain insights into the origin and evolution of this well-known floristic disjunction, one of the first order biodiversity patterns in the Northern Hemisphere that has been extensively studied (Wolfe 1980; Boufford and Spongberg 1983; Tiffney 1985a; Xiang et al. 1998b, 2000; Wen 1999; Donoghue and Smith 2004; Wen et al. 2010).

The chestnut genus *Castanea* Miller (Fagaceae) includes seven tree species, restricted to EA and ENA, with one species endemic to Europe. The species were divided into three sections (Dode 1908): section *Eucastanon* Dode, including the five species with three nuts per cupule: *C. mollissima* Blume and *C. seguinii* Dode from China and *C. crenata* Siebold & Zucc. from Japan, *C. dentata* (Marshall) Brokh. from North American, and *C. sativa* Mill. from Europe. Sections *Balanocastanon* Dode and *Hypocastanon* Dode each is monotypic including a single species and both make fruits containing one nut per cupule. Section *Balanocastanon* contains *C. pumila* (L.) Mill. from North America and Section *Hypocastanon* contains *C. henryi* (Skan) Rehder & Wilson from China. Within *C. pumila*, two varieties were recognized by Johnson (1988) and Nixon (1997), *C. pumila* in the southeastern United States and *C. pumila* var. *ozarkensis* (Ashe) A.E. Murray limited to the Ozark mountains. Phylogenetic studies of *Castanea* were previously conducted using data from six chloroplast regions in Lang et al. (2006, 2007). The studies found that sect. *Eucastanon* is paraphyletic and sect. *Balanocastanon* and sect. *Hypocastanon* did not form a sister group. Authors of these studies also hypothesized an eastern Asia origin of *Castanea* in the Eocene, followed by range expansion to Europe and North America in the mid Eocene, as an example of the “Out of Asia” pattern (Donoghue and Smith 2004).

The witchhazel genus *Hamamelis* L. (Hamamelidaceae) is also a small genus consisting of four or six species of shrubs and small trees isolated in EA and ENA. The EA species include *H. mollis* Oliv. from eastern and southern China (Chang 1979; Zhang and Lu 1995) and *H. japonica* Siebold & Zucc. from Japan (Sargent 1890; Ohwi 1978). The ENA species include *H. virginiana* L., that is widely distributed from Canada to the Gulf coast (Bradford and Marsh 1977), *H. vernalis* Sarg., a species endemic to the Ozark Mountains in Arkansas, Missouri, and eastern Oklahoma (Bradford and Marsh 1977), *H. ovalis* S.W. Leonard that is restricted to a small area of Mississippi (Leonard 2006), and *H. mexicana* Standl. endemic to northeastern Mexico (Standley 1937), which is also known as *Hamamelis virginiana* var. *mexicana* (Standl.) C.Lane. A few phylogenetic studies of *Hamamelis* have been conducted using data from ITS, ETS, waxy gene, and several plastid genes (Wen and Shi 1999; Li et al. 2000; Xie et al. 2010). However, the species relationships within *Hamamelis* have remained uncertain due to low nodal support values and short internal branches, especially regarding the relationships within the ENA clade.

In this study, the first goal is to evaluate the relative merits of RAD-seq and target enrichment methods for addressing within-genus-level phylogenetic researches, through gene tree and species tree comparisons, as well as divergence time comparisons. Our second goal is to generate a new pipeline for detecting more putative paralogs for downstream enrichment data analyses rather than just using the paralog detection method from HybPiper. We compared multiple sequence alignment (MSA) matrices generated using our pipeline with the MSA matrices generated from HybPiper using different character coding and trimming methods (See Methods). We further compared the phylogenies and divergence times resulted from analyses of the different data matrices to determine the effects of heterozygotes and paralogs on phylogenetic and divergence time analyses. Finally, we addressed longstanding questions regarding the evolutionary history of *Castanea* and *Hamamelis* and the EA-ENA floristic disjunction. Combining the molecular and fossil records, we conducted rigorous phylogenomic and comprehensive biogeographic analyses to uncover their geographic origins, migration routes, and past hybridization and to gain new insights into the formation of the EA-ENA disjunct phytogeographic pattern.

## MATERIALS AND METHODS

### Data Generation

#### DNA samples preparation

A total of 58 individuals were sampled and sequenced. The samples consisted of 30 plants of *Castanea* and 28 plants of *Hamamelis*, including two and three outgroup individuals, respectively (Table 1). Outgroup species were chosen based on their phylogenetic positions in Fagaceae and Hamamelidaceae, respectively inferred by Lang et al. (2006) and Xie et al. (2010). *Fothergilla* and *Parrotiopsis* were used as the outgroups of *Hamamelis* while *Quercus* was used as the outgroup of *Castanea*. The sampling included all species of the two ingroup genera recognized in modern floras. Leaf samples were collected from the fields or plants grown in arboreta or botanical gardens (Table 1). Fresh leaves were stored in silica gel to dry. The dry leaves were stored at -20 °C until they were used for the DNA extraction. All the vouchers were deposited in the herbarium of North Carolina State University (acronym NCSC).

**Table 1.**
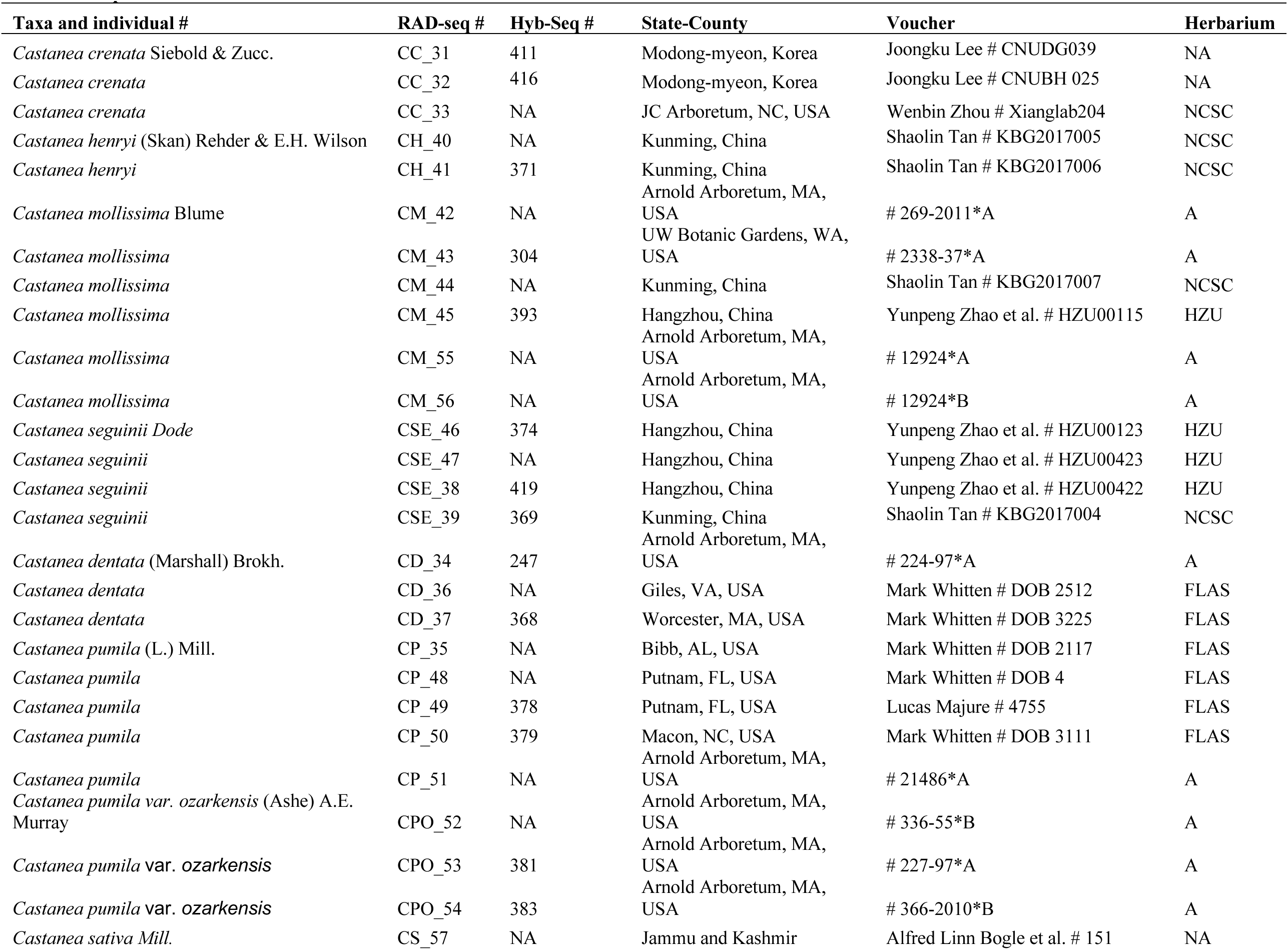
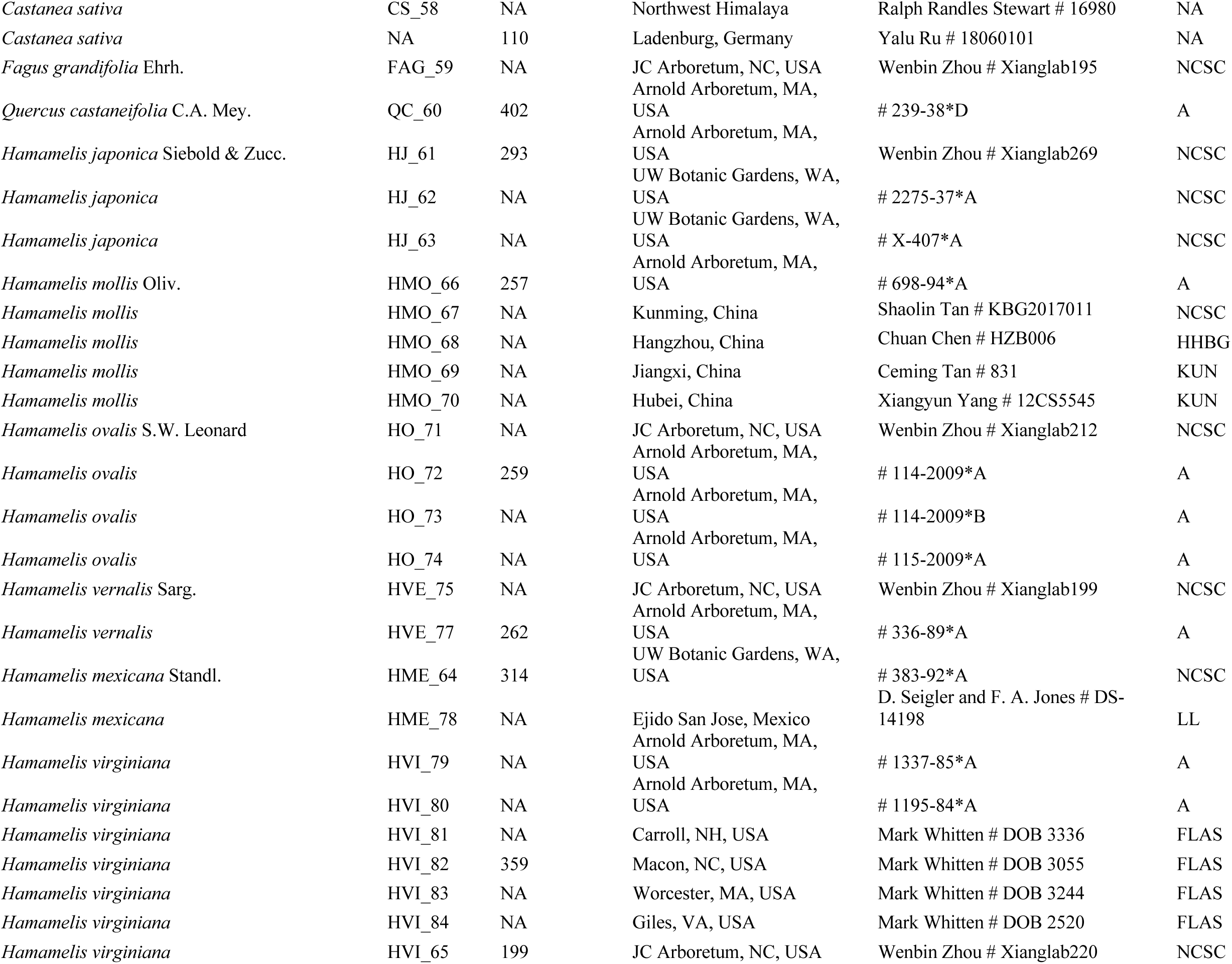
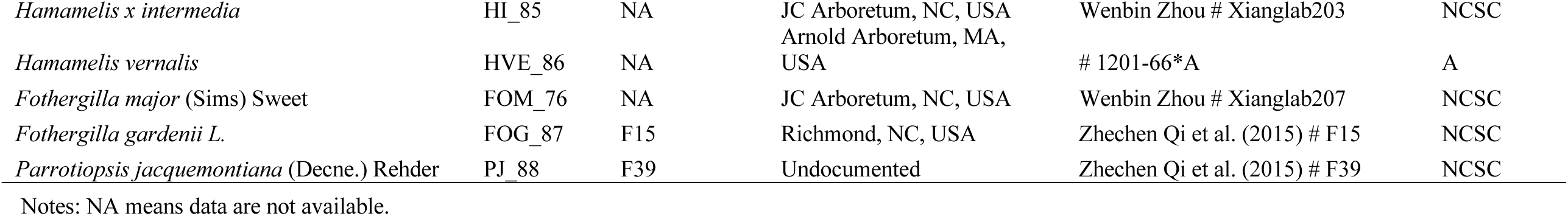
Sample information.

Total genomic DNAs were extracted from leaf samples using the CTAB protocol (Doyle 1991) with modification described in Cullings (1992) and Xiang et al. (1998a). For leaf samples of *Castanea* that are rich in secondary compounds, they were washed five times with 0.8 mL of a washing buffer containing 10% polyethylene glycol, 0.35 M sorbitol, 50 mM Tris-HCl, 0.1% bovine serum albumin and 0.1% β-mercaptoethanol (Sakaguchi et al. 2018; Zhou et al. 2020) prior to DNA extraction with the modified CTAB method. The quantity and quality of DNA samples were first checked by 1% agarose gel electrophoresis followed by measurement on a Nanodrop spectrophotometer (ThermoFisher) and with a PicoGreen fluorescent dye assay (Life Technologies, ThermoFisher).

#### Library preparation for RAD-seq and sequencing

Double digest RAD-seq (ddRAD) (Peterson et al. 2012) libraries were prepared following Pais et al. (2018, 2020) and Du et al. (2020) for all samples. Two restriction enzymes, *PstI-HF* and *MspI* were used to digest 350 ng of the total DNA from each sample (Table 1). We used the *PstI* adaptors and forward primer described in Pais et al. (2017) and Zhou et al. (2018), but modified the protocol with the *MspI* adaptor and reverse primer from Du et al. (2020). Libraries were pooled and size selected to 400-600 bp using a Pippin prep (Sage Science, Beverly, MA, USA) and verified by analysis on Bioanalyzer 2100 (Agilent Technologies, Santa Clara, CA, USA). Size-selected libraries were enriched by PCR for 12 cycles, followed by purification with AMPure Beads before delivered to the Genomic Science Lab (GSL) at North Carolina State University for sequencing. Single-end sequencing (150 bp) from the end of the *PstI* cutting site was performed on an Illumina HiSeq 2500 platform.

#### Library preparation of Angiosperm 353 gene enrichment and sequencing

Due to the relative high cost of sequence capture, we reduced the samples in this experiment to include 16 *Castanea*, eight *Hamamelis*, and four outgroups (Table 1), which also covers all species of the two genera. A total of 1000 ng DNA of each sample concentrated to ∼35 μL was delivered to Rapid Genomics Lab (Gainesville, Florida, USA) for Hyb-Seq library reconstruction and sequencing. The DNA samples were pooled for hybridization to biotinylated probes using the Angiosperm-353 v. 1 target capture kit (Johnson et al. 2019) available from Arbor Biosciences (Arbor Biosciences, Ann Arbor, Michigan, USA). Sequencing of DNAs pulled from the hybridization experiment was performed with Illumina MiSeq (Illumina, San-Diego, California. USA) to produce 2 x 150 bp paired-end reads, as described in Gaynor et al. (2020).

### Locus Data Assembly and MSA Generation

#### RAD-seq data

Raw sequence reads were analyzed in ipyrad v.0.7.28 (Eaton and Overcast 2020), summarized briefly here. The reads were first de-multiplexed and then trimmed to remove specific barcodes and adapters. Then, low quality reads (with Q score < 20) were trimmed and filtered. To remove the influence of DNA sequences from the plastid genome and microorganisms on phylogenetic inference, chloroplast genomes of *Castanea mollissima* (NC_014674, Jansen et al. 2011) and *Hamamelis mollis* (NC_037881, Dong et al. 2018) and the full bacterial and fungal database (https://ftp.ncbi.nlm.nih.gov/refseq/release/) were downloaded from NCBI as the references. We chose the “denovo-reference” function in ipyrad, which uses VSEARCH (Rognes et al. 2016) to cluster reads of the same locus and uses BWA (Li and Durbin 2009) and BEDTools (Quinlan and Hall 2010) to remove the reads/loci mapping to the plastid and microorganisms references. We selected a default threshold of 85% of similarity in sequences for reads clustering in ipyrad based on results in Du et al. (2020) for orthologs identification among plant species. We also used 2 as the maximum alleles per sample in the consensus sequence construction for each locus in a sample because both *Castanea* and *Hamamelis* are diploid organisms and a maximum of 50% samples sharing heterozygosity at a site (i.e., 0.5 shared heterozygosities in ipyrad) of a locus as the cut-off of orthology. The resulting read clusters were aligned across samples using MUSCLE (Edgar 2004). The loci were finally trimmed (5 bases on 3’ end) and filtered if they contained over 30 SNPs or 8 indels. To compare across different levels of missing data, we output matrices containing full-length loci in different minimum thresholds of samples using the parallel branching assembly function. We selected thresholds from at least 20% of samples present at a locus, to 80%, increasing in increments of 10%. We named these matrices based on this threshold, e.g. M20, which contained all loci present in at least 20% of samples. All matrices have been uploaded to Dryad.

#### Hyb-Seq data

All samples for Hyb-Seq were demultiplexed using Illumina’s BCLtofastq by Arbor Biosciences. Raw sequencing reads were then cleaned and trimmed by Trimmomatic v.0.38 (Bolger et al. 2014) using parameters MAXINFO:100:0.5 and TRAILING:20. Subsequently, the HybPiper pipeline v. 3 (Johnson et al. 2016) was used to recover both coding sequences (CDS) and their flanking intron/non-coding regions. The process includes three major steps: using the nuclear sequences of Angiosperm 353 genes (Johnson et al. 2019) as the references to capture all the reads from sequenced accessions via the BWA option (Li and Durbin 2009), applying the SPAdes (Bankevich et al. 2012) to assemble reads into long contigs, and implementing the intronerate.py module to recover “intron” and “supercontig” (CDS + intron fragments) sequences. Finally, we generated multiple Hyb-Seq data matrices to assess the phylogenetic effects of character coding, data trimming, and paralogous genes in order to find the best dataset for comparison with the RAD-seq data and for assessing their impacts on divergence time. Specifically, we identified loci containing putative paralogs using our new pipeline, named putative paralog detection (PPD) (available on Github: https://github.com/Bean061/putative_paralog) and built the following matrices for comparisons: exon vs. supercontig matrices of orthologous loci, paralogous loci, or all genes (including the paralogous genes), respectively. The exon matrices contained the sequences of the coding regions only while the supercontig matrices contained sequences of both coding and their flanking regions of the three respective gene groups. The pipeline includes two major steps that occur after generating alignment matrices in HybPiper or other enrichment pipelines: first, “*degenerated*” sequences are generated from mapping reads back to contigs, which applies the IUPAC ambiguity codes to capture the heterozygous sites in generating the exon and supercontig matrices of the three sets of aforementioned genes; and second, the matrix is trimmed of highly heterozygous sites, misaligned regions, and particularly gappy columns, as one trimming step which is shown in Figure 2 (see details in Supplementary Information available on Dryad). Thresholds for trimming are configurable by the user. To directly compare the impact of character coding, we compared the “*degenerated*” orthologous matrix generated through both steps of PPD with the orthologous data matrix built from HybPiper without degenerate character coding in the matrix (referred to as “*consensus*” matrices) and also trimmed these matrices with the second step of PPD (referred to as the “well-trimmed *consensus*” matrices). We also compared data from the Angiosperm 353 genes with the plastid genes captured by off-target in the Hyb-Seq experiment, and the combined genes from RAD-seq and Hyb-Seq (excluding plastid genes). In addition, we compared data from exon and supercontig, paralogs and orthologous, and applied three different trimming approaches (no trimming, automated trimming in Trimal, and our “well-trimmed” method in PPD) in producing these matrices (see details in Supplementary Information, available on Dryad). For all the Hyb-Seq data matrices below, if no trimming description is directly indicated, the data matrices were “well-trimmed” through PPD (PPD Part 2: s1-s7 in Fig. 2). All Hyb-Seq comparisons are listed in Table 2 (see details in Supplementary Information, available on Dryad).

**Table 2.**
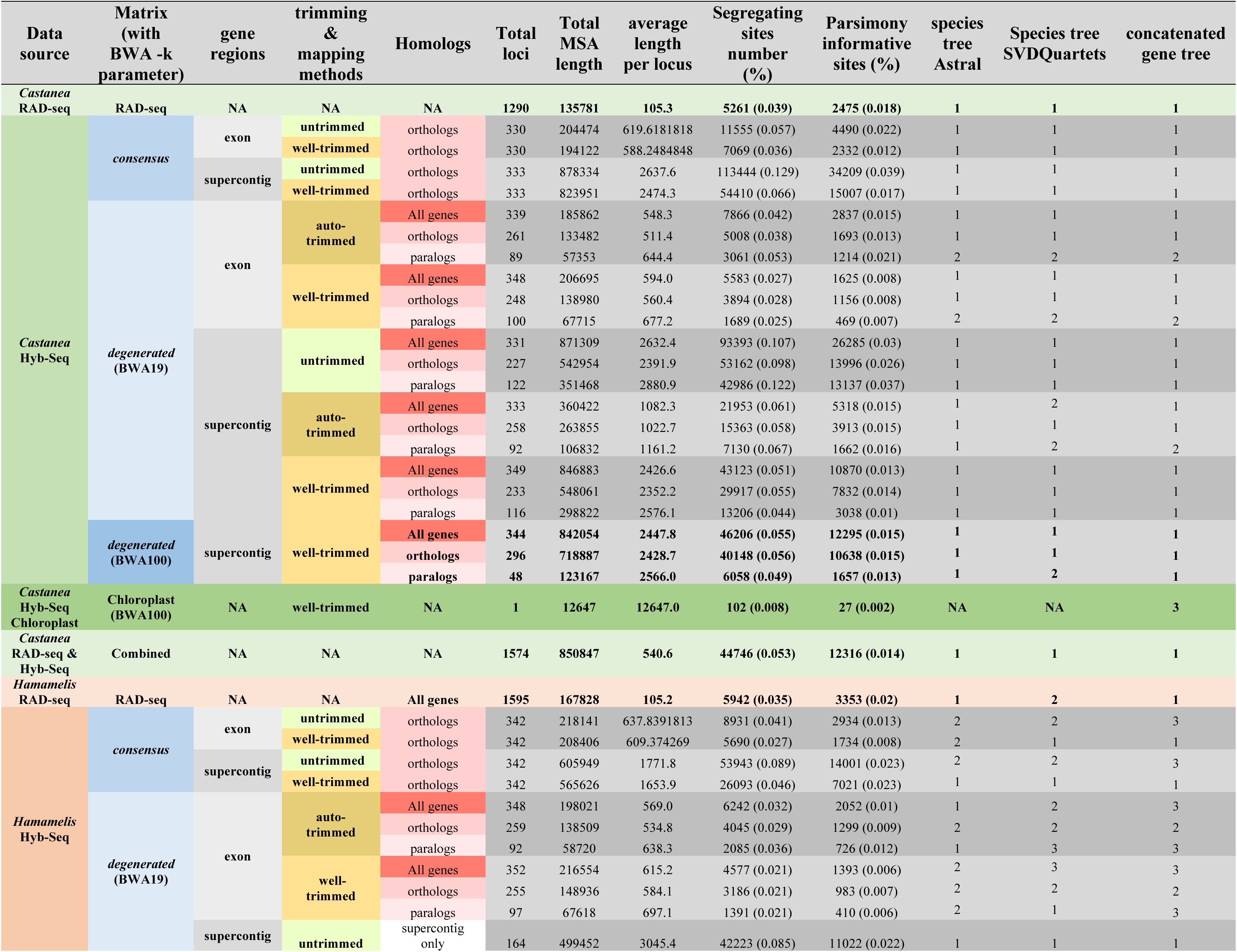
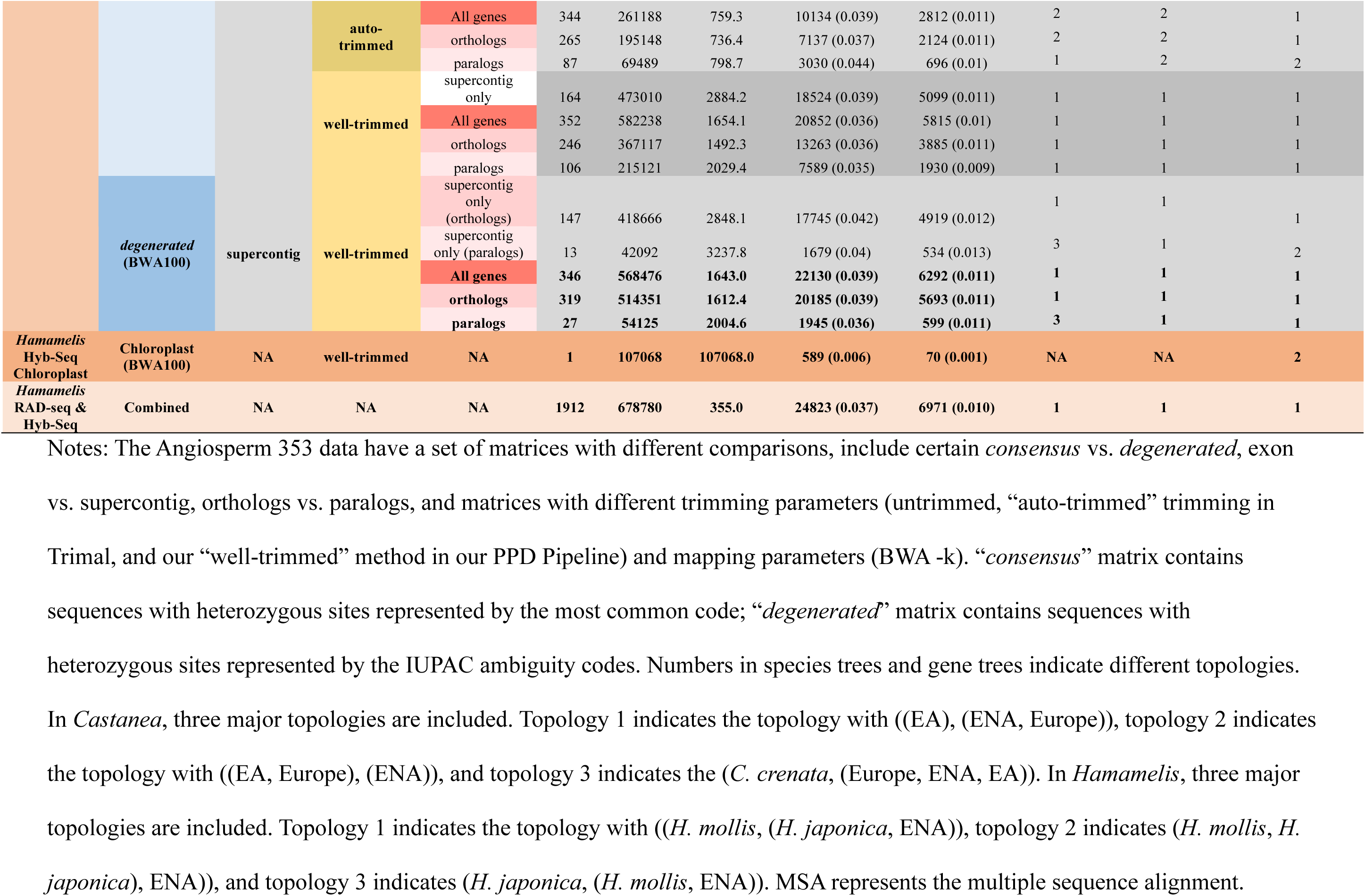
Summary information of RAD-seq data, Angiosperm 353 data, chloroplast plastid data, and combined RAD-Hyb-Seq data, including the loci number, total MSA length, average length per locus, segregating sites number and percentage, parsimony segregating sites number and percentage, and concatenated gene tree and species tree topologies.

### Phylogenetic Analyses

#### Phylogeny of RAD-seq data

Preliminary phylogenetic analyses of all matrices (M20-M80) from RAD-seq were initially conducted using a partitioned maximum likelihood (ML) analysis implemented in the IQ-Tree program (Nguyen et al. 2015). All analyses used the TESTNEW option to obtain the best molecular model per partition. The results showed that the trees derived from different datasets were congruent in their topologies while the M50-based tree had the highest support values at most nodes. Due to this reason and the higher number of loci compared to M60-M80 and lower level of missing data compared to M20-M40 (see Results), we used the M50 dataset for final phylogenetic and downstream analyses. The M50 dataset of each genus was further filtered by removing samples containing 30% or more uncertain bases (Ns) and hybrid species *Hamamelis × intermedia* that contains many degenerate bases in the concatenated sequence. This filtering resulted in the removal of 2 *C. sativa* samples (CS_57 and CS_58) from *Castanea* and 3 samples (HI_85, HME_78, and HVE_75) from *Hamamelis*. A total of 25 samples of *Castanea* (plus one outgroup) and 22 samples of *Hamamelis* (plus three outgroups) representing all species of the two genera were remained for further analyses.

We conducted phylogenetic analyses of the filtered data using both the ML and Bayesian Inference (BI) methods. The ML analyses were partitioned by locus and were conducted using both IQ-Tree (Nguyen et al. 2015) and RAxML v8.2.4 (Stamatakis 2014), while the BI analysis was conducted using MrBayes v.3.2.2 (Ronquist and Huelsenbeck 2003). Analyses with IQ-Tree used the modelfinder (flag -m TESTNEW) and 1000 replicates of ultrafast bootstrap. The TESTNEW option was applied to find the best molecular model for each locus. In the ML analysis with RAxML, jModelTest v.2.1.4 (Darriba et al. 2012) was used to determine the best substitution model based on AICs (GTR+Γ and HKY+Γ models were chosen for *Castanea* and *Hamamelis*, respectively), with 1000 bootstrap replicates. For BI analysis, the HKY+Γ model was chosen for both *Castanea* and *Hamamelis* according to the BIC in jModelTest v.2.1.4. Two independent runs of Markov Chain Monte Carlo (MCMC) analyses were executed for each genus; each run was improved by the Metropolis Coupling method, which includes one cold chain and three heated chains that started randomly in the parameter space. Ten million generations with tree sampling in every 1000 generations and a burn-in of the first 25% of sampled trees were set for each run. Convergence was assessed by evaluation of standard deviation of split frequencies between tree samples (which approached zero, <0.001), the effective sample size (ESS >200 in all cases), and by evaluating the plot of the likelihood scores from the resulting MrBayes trees in Tracer (Rambaut et al. 2018). The IQ-Tree, RAxML, and BI analyses were all conducted on the CIPRES Science Gateway Portal (Miller et al. 2010).

We constructed species trees using ASTRAL-III (Zhang et al. 2018) and SVDquartets (Chifman and Kubatko 2014). For the analyses with ASTRAL-III, we used IQ-Tree as above to generate gene trees for both genera as inputs and ran ASTRAL-III with the default parameters. For the analyses with SVDquartets, PAUP* v4.0a166 (Swofford 2003) was used to generate a total of 100,000 quartets with 100 bootstrap replicates and then the quartet assembly method QFM was used to produce a summary tree (Reaz et al. 2014), following Zhou et al. (2020). All gene trees and species trees were visualized and edited in FigTree v.1.4.2 (Rambaut 2012) and edited with ggtree [R] (Yu et al. 2018) and Adobe Illustrator 2020 (Adobe Systems, San Francisco, CA, USA).

#### Phylogeny of Hyb-Seq data

Phylogenetic analyses of the Hyb-Seq data were also performed for the various data matrices listed in Table 2 with IQ-Tree (Nguyen et al. 2015) to generate concatenated gene trees, and with both ASTRAL-III (Zhang et al. 2018) and SVDquartets (Chifman and Kubatko 2014) to generate the species trees, as described above for the RAD-seq data. Phylogenetic analyses of the plastid DNA data were performed with both IQ-Tree and RAxML v8.2.4 (Stamatakis 2014).

#### Phylogeny of the combined data

The combined RAD-seq and Hyb-Seq data (primary Hyb-Seq-PPD, see Results) matrix (referred to as RAD-Hyb-Seq) was built from M50 and “*degenerated*” supercontig matrix of orthologous genes containing accessions with both RAD-seq and Hyb-Seq data available. We checked overlapping sequences between the two datasets by mapping RAD-seq data to the target reference genes using ipyrad and manually removed the redundant sequences (see Results below). Phylogenetic analyses of the combined data were also performed using IQ-Tree (Nguyen et al. 2015), ASTRAL-III (Zhang et al. 2018), and SVDquartets (Chifman and Kubatko 2014) to infer the “total evidence” gene phylogeny and species tree of both *Castanea* and *Hamamelis*.

### Dating the Phylogenies

BEAST2 2.6.2 (Bouckaert et al. 2014) was employed to estimate the divergence times of lineages within each genus. BEAST2.6.2 can consider information at heterozygous sites in divergence time estimation. The divergence time analyses were conducted for both the RAD-seq data and the Hyb-Seq data. We reduced the RAD-seq data in sampling to include only the samples with Hyb-Seq data in the analyses so that the two types of data included identical samples (Table 1). The only exception was that RAD-seq data were missing for *Castanea sativa* due to the failure of herbarium samples in the RAD-seq experiment and lack of silica dried leaf samples at the time the experiment was conducted. Among the different Hyb-Seq data matrices, we performed divergence time dating analyses for several matrices for each genus, including the “*degenerated*” and “*consensus*” supercontig matrices for orthologous genes, the “*degenerated*” exon matrix for orthologous genes, the “*degenerated*” supercontig matrix for paralogous genes, the combined RAD-Hyb-Seq data, and matrices using different trimming processes to allow comparisons of supercontig and exon and the effect of coding methods, paralogous genes, and trimming methods. The stem ages of *Castanea* and *Hamamelis* were constrained based on fossil evidence, as 66 to 72 Ma (lognormal) and 50 to 56 Ma (lognormal), respectively (for details, see Supplementary Information, available on Dryad).

For the RAD-seq data, divergence-time analyses were run under the HKY+Γ model for both *Castanea* and *Hamamelis* according to the result of jModelTest, an uncorrelated lognormal relaxed clock (Drummond et al., 2006), and the birth–death process model (Stadler, 2010). We set the mean GrowthRate (net diversification rate) to have a uniform distribution with a range of 0– 100, with an initial value of 0.0, and the relative Death Rate (extinction rate/speciation rate) to have a 0–1 range, with an initial value of 0.5. These values were chosen based on the estimated average net diversification rate and extinction rate in plants (De Vos et al., 2014). All the other prior parameters were determined with empty runs without the sequence data. The empty runs were conducted to check the appropriates of prior settings; the posterior distribution of the divergence times of the constrained nodes produced from the empty runs are expected to cover or be similar to the constrained values with ESS over 200. The priors in the empty runs were then used in the real divergence time analyses with data. We ran the analyses with data for 200 million generations, with sampling of trees every 10,000 generations. Quality of the runs and parameter convergence were assessed using Tracer v.1.6.0. The maximum credibility tree of mean heights was then constructed using TreeAnnotator after discarding 20% trees as burn-in. We compared different Hyb-Seq matrices using the same fossil calibrations and same priors except substitution models which were decided by BIC values from jModelTest. For analyses with the Hyb-Seq data, the divergence time analyses were conducted in the same way as for the RAD-seq data, except using the GTR+Γ molecular model for all orthologous gene matrices and HKY+Γ molecular model for paralogous gene matrices. For the RAD-Hyb-Seq data, the GTR+Γ model was chosen in the divergence time analysis. These models were chosen based on jModelTest.

### Biogeographic Analyses

#### Total-evidence dating

For our biogeographic analyses, we added fossils of *Castanea* and *Hamamelis* to the phylogeny to provide a more accurate evolutionary framework for the study. *Castanea* and *Hamamelis* have a number of fossils from the Paleogene and Neogene periods in the Northern Hemisphere (listed in Table 3 and see details in Supplementary Information, available on Dryad). We estimated divergence times of both genera with a total-evidence dating approach (Ronquist et al. 2012; Gavryushkina et al. 2017) in BEAST2 (Bouckaert et al. 2014) with a matrix that included both extant and fossil taxa and data from molecules for extant taxa and morphological characters for fossils and extant taxa. We were able to score seven morphological traits available in fossils of *Castanea* and six morphological traits available in fossils of *Hamamelis* (Supplementary Table S1, available on Dryad). The molecular matrix used in the analysis was the “*degenerated*” supercontig matrix of orthologous genes from Hyb-Seq, as this data set contains the most informative characters of orthologous genes and supported phylogenetic relationships consistent with the completely resolved species tree and the RAD-seq trees (see Results below). The molecular characters were recorded as missing in the fossil taxa. Following previous studies (Gavryushkina et al. 2017; Zhou et al. 2020), the fossilized birth death model (FBD) and relaxed clock log normal were selected for these analyses. The GTR+Γ was chosen for the molecular substitution model and the Lewis MK model (Lewis 2001) was chosen for the morphological data. A preliminary analysis with FBD showed unresolved placement of most fossil taxa with unexpected divergence times, similarly reported in a recent simulation study on tip-dating method (Luo et al. 2020), probably due to homoplasy of the few morphological characters available. Therefore, we modified the analysis to constrain the fossils with their most closely related clades or species based on paleobotany literature, with dates for fossils set as the value in between the age range of each fossil (Table 3). We also enforced the species tree backbone derived from the molecular data in the analysis to reduce the run time and ensure the recovery of relationships of extant species depicted in the species tree. We ran the analysis for 200 million generations with sampling of trees every 10,000 generations. Quality of the runs and parameter convergence were assessed using Tracer v.1.7.1. The maximum credibility tree of median heights was constructed using TreeAnnotator (Bouckaert et al. 2014) after discarding 20% trees as burn-in. As a comparison, we also constructed a dated total evidence tree using a matrix containing the combined data from RAD-seq and Hyb-Seq and the morphological characters using the same processes described above. The total evidence dating analysis was also repeated with the combined RAD-Hyb-Seq data.

**Table 3.**
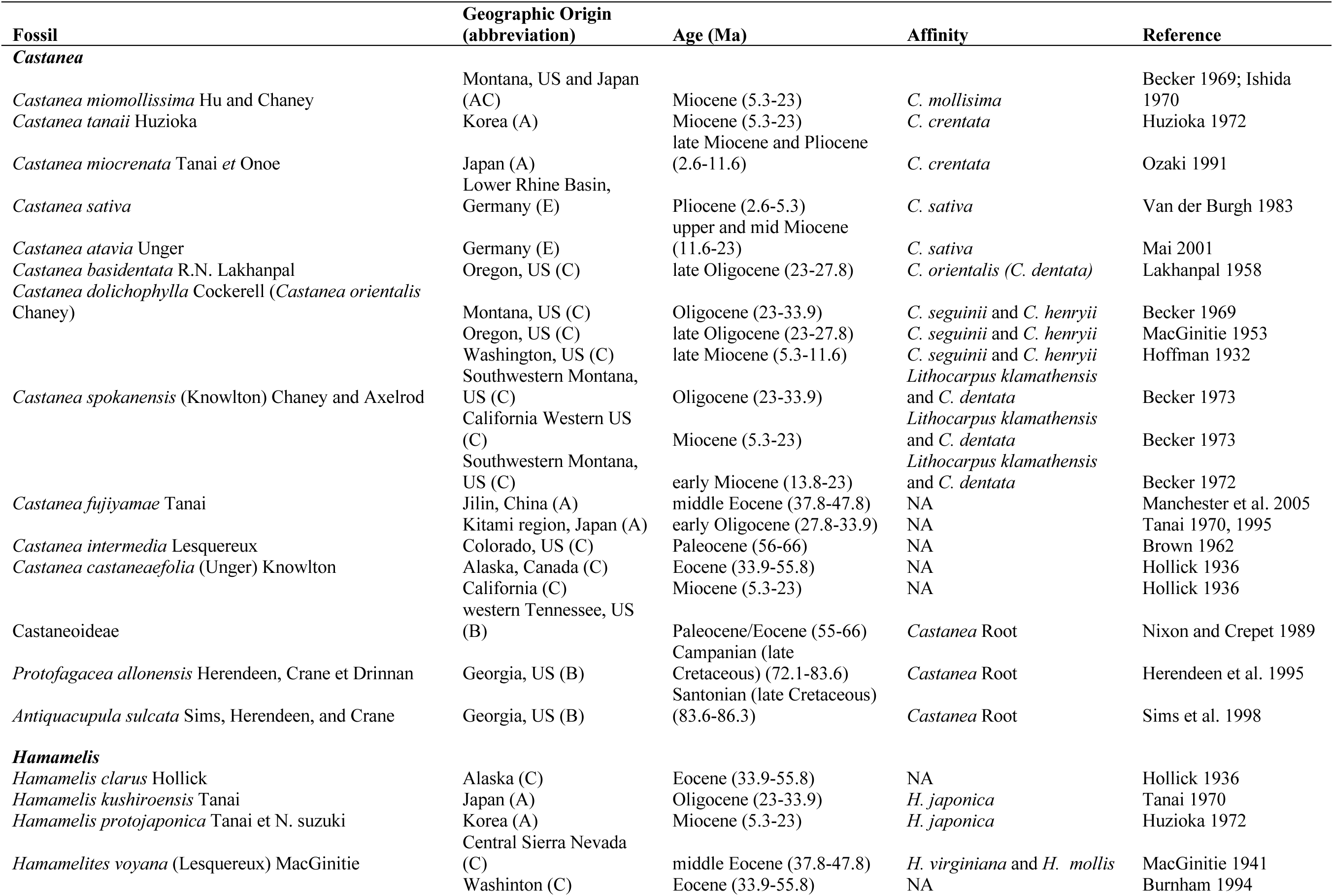
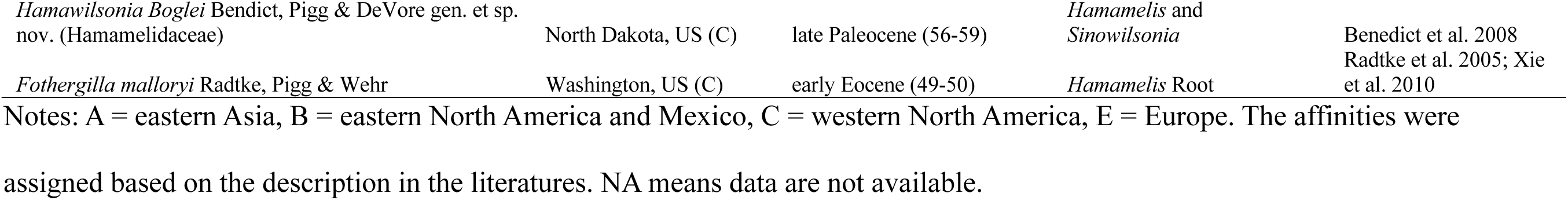
Fossils reported for *Castenea* and *Hamamelis*.

#### Reconstructing the biogeographic history with DEC

We conducted the biogeographic analyses using the Dispersal-Extinction-Cladogenesis (DEC) method (Ree and Smith 2008) based on the dated total evidence trees (from Hyb-Seq data only and the combined RAD-Hyb-Seq data) including all fossil species for *Castanea* and *Hamamelis*. The selection of this method was based on a test in BioGeoBEARS (Matzke 2013) in RASP (Yu et al. 2015) that compared among the Dispersal-Vicariance Analysis (DIVA; Ronquist and Cannatella 1997), Dispersal-Extinction-Cladogenesis (DEC), and BayArea methods (Landis et al. 2013). We did not consider the parameter j (“jump”) that represents the founder-event, allowing daughter species to “jump” outside the geographical range of parental species (Matzke 2013) in the test, due to concerns raised in Ree and Sanmartín (2018). Among the three methods compared, the DEC method had the highest AICc value. We defined five geographic regions for both *Castanea* and *Hamamelis*, in the DEC analyses to cover the distributions of modern and extinct taxa: EA (“A”); ENA and Mexico (“B”); western North America (WNA) (“C”); and Europe (“E”) (Table 3). We combined Mexico with eastern North America according to the more or less continuous distribution between *H. mexicana* and ENA clade. We implemented a matrix of relative dispersal rates in time slices between areas (Supplementary Table S2, available on Dryad) based on largely the physical connections between areas, according to (Graham 1999a, 1999b, 2018; Tiffney and Manchester 2001; Mann 2007). The rates were arbitrarily set using the following criteria: as 0.001 for the time slices when the two areas were physically disconnected or isolated by barriers (such as water or dry areas); as 1.0 when the two areas were physically and ecologically connected, and as values between 0.001 – 1.0 (e.g., 0.2, 0.5, or 0.75) when the two areas were partially connected by islands that were separated at different levels (Supplementary Table S2, available on Dryad). For ancestral ranges with disjunctions, e.g., AB, CE in both *Castanea* and *Hamamelis*, we also set the probability value to 0.001 for low probability because such an ancestral distribution would require frequent intercontinental long distance dispersal to maintain the disjunct populations as a single gene pool of the ancestor across disjunct areas which we believe is unlikely for *Castanea* and *Hamamelis* given the expected short-distance dispersal of their seeds (heavy in *Castanea* and likely dispersed by mammals; gravity-dispersed in *Hamamelis* ejected by capsules). For both analyses, we implemented a constraint of maximum areas of four to allow all four areas of the Northern Hemisphere to be included in an ancestral range.

### Test for ILS and Introgression

Both incomplete lineage sorting (ILS) and introgression can result in phylogenetic incongruences among genes. We applied the D-statistic (ABBA-BABA) method in Dsuite (Malinsky 2019) and phylogenetic network method to test which of these causes may explain the conflicts among gene trees in *Castanea* and *Hamamelis*. The D-statistic test provides a measure of introgression between taxa given a guide tree topology (((P1, P2), P3), O), where O represents a closely related outgroup taxon and P1 and P2 are taxa tested for signatures of introgression with P3. The null hypothesis is as follows: if ILS is the only source of gene tree discordance, then the frequency of derived alleles exclusively present in P1 and P3 is expected to be equal to the frequency of derived alleles exclusively present in P2 and P3 (D = 0). However, if the introgression happened between P3 and P2 or P3 and P1, it will result in an excess of shared derived alleles between the taxa that experienced introgression (D > 0 or D < 0). The Dsuite is a tool for calculating D-statistics based on VCF files, assessing the significance via the jackknife test, and generating two-tailed P-values based on the Z-scores. To explain conflicts between the concatenated-gene tree and species tree, we performed two D-statistic tests for *Castanea* and *Hamamelis* to assess ILS vs. introgression among specific lineages involved in the phylogenetic conflicts. Specifically, for *Castanea*, we performed the test with the Hyb-Seq data (primary Hyb-Seq-PPD, see Results) (as it contains complete species samples while the RAD-seq data had *C. sativa* missing) and combined RAD-Hyb-Seq data. We tested for introgression events among ENA *Castanea*, EA *Castanea*, and European *Castanea* (*C. sativa*) due to the weak support for the sister relationship between *C. sativa* and the ENA clade in the species tree, but strong support in the concatenated gene tree (see Results) and among *C. crenata*, the clade consisting of the remaining eastern Asian species, and the clade containing the ENA-*C. sativa* due to their conflicting relationships between the chloroplast gene tree and the nuclear gene tree (see Results). For all tests, the outgroup species used in phylogenetic studies, *Quercus castaneifolia*, was used as the outgroup. For *Hamamelis*, due to the low support in both the concatenated gene tree and the species tree at the two lower nodes of the phylogeny, involving relationships among the *H. mollis*, *H. japonica* (both EA), and ENA *Hamamelis* clade, see Results), we conducted the test with three data sets, the RAD-seq, Hyb-Seq data, and combined RAD-Hyb-Seq data for the presence of introgression. Similarly, the same outgroup taxa used in phylogenetic studies were used as the outgroup for the D-statistic test. We also performed the test among *H. mexicana*, *H. vernalis*, and *H. ovalis* within the ENA clade, due to low support of their relationships, using *H. virginiana* as the outgroup. Input VCF files were generated with ipyrad for the RAD-seq data and with msa2vcf in JVarkit (Lindenbaum 2015) for the Hyb-Seq data.

We further conducted analyses of phylogenetic networks to confirm the potential introgression events among species or lineages within each genus suggested by the D-statistic tests. For both *Castanea* and *Hamamelis*, we used the quartet-based maximum-pseudolikelihood approach implemented in PhyloNet v.3.6.4 (Than et al. 2008; Yu and Nakhleh 2015) for the purpose. The set of rooted gene trees from the Hyb-Seq data (primary Hyb-Seq-PPD, see Results) were used as the input for PhyloNet. We did not use RAD-seq loci as their average length was short relative to Hyb-Seq data, resulting in unreliable gene-tree topologies. The number of runs of the search (-x) was set as 100 and the maximum number of reticulations was set as 2 according to the results from the D-statistic tests. All other parameters were left as default settings, as recommended by the authors.

## RESULTS

### RAD-seq and Hyb-Seq Data

#### Locus recovery

From RAD-seq, an average of 659,529 and 698,799 reads per sample were generated in *Castanea* and *Hamamelis*, respectively. After quality filtering by the ipyrad pipeline, an average of 659,405 reads were retained in *Castanea* and 698,651 reads remained in *Hamamelis* for the subsequent cluster procedure (Supplementary Table S3, available on Dryad). The clustering process resulted in matrices of M20 to M80 containing 3597 to 415 loci, respectively, in *Castanea* and 3935 to 395 loci, respectively, in *Hamamelis* (Supplementary Table S4, available on Dryad). After filtering, the length of each locus from RAD-seq is approximately 105 bp for both *Castanea* and *Hamamelis* (Table 2).

For Hyb-Seq datasets, the number of loci varied depending on gene region, filtering strategy, and categorization of genes in the alignment (e.g. ortholog vs. paralog or exon vs. supercontig). For complete the loci recovery and sequence length information in different datasets, see Table 2. The average length per locus in exon region of well-trimmed “*degenerated*” matrix of orthologs was approximately 594 bp in *Castanea* and 615 bp in *Hamamelis*, while the average length per locus in the well-trimmed “*degenerated*” supercontig matrix was much longer, approximately 2448 bp in *Castanea* and 1643 bp in *Hamamelis*. The information on loci number, alignment length, average length per locus, number of segregating sites, and number of parsimony informative sites of all matrices derived from different trimming methods are available in Table 2. Overall, many more loci were obtained from RAD-seq compared to Hyb-Seq (344 genes) but the average length per locus is much longer in the Hyb-Seq data, resulting in longer alignment lengths in Hyb-Seq matrices (Table 2).

For the combined RAD-Hyb-Seq data, we first checked for duplication of loci present in both data sets. We found 12 loci (out of 1290 in *Castanea*) and 2 loci (out of 1595 in *Hamamelis*) in the RAD-seq data of the two genera that could be mapped to the Hyb-Seq data. After removing these loci from the RAD-seq M50, we obtained 1574 loci for *Castanea* and 1912 loci for *Hamamelis* (Table 2) for the combined RAD-Hyb-Seq data. The chloroplast DNA data matrices extracted from the Hyb-Seq data due to off-targeting contained 12,647 bp for *Castanea and* 107,068 bp for *Hamamelis* (Table 2).

#### Paralog detection in Hyb-Seq data

The gene matrices generated by HybPiper showed 11 putative paralogs (Gene 6048, 6954, 4951, 4724, 5940, 6387, 6570, 7583, 7324, 5138, 5941) out of a total of 344 genes in *Castanea*, but only two putative paralogs (Gene 5463, 5347) out of 344 genes in *Hamamelis*. In contrast, our PPD pipeline detected all paralogs detected by HybPiper, as well as many additional paralogs. For *Castanea*, the PPD detected a total of 48 paralogs, and for *Hamamelis*, PPD detected a total of 27 paralogs (Table 2).

#### Final Hyb-Seq datasets

Although we completed numerous analyses on all data matrices presented in Table 2, we have chosen to present results in the main text primarily for the well-trimmed “*degenerated* BWA100” supercontig matrix of orthologs generated from our PPD pipeline. We will refer to the *Castanea* and *Hamamelis* alignments generated through this pipeline as primary Hyb-Seq-PPD datasets. Results of additional analyses performed on alternative matrices were presented in Supplementary Information (available on Dryad).

### Phylogenetic Analyses

#### Castanea - RAD-seq

The analyses of all different matrices (M20, M30, …, M80) with IQ-Tree yielded phylogenetic trees with congruent topology (Supplementary Figs. S1-S7, available on Dryad). The M50 matrix was selected for detailed phylogenetic and divergence time analyses (due to its relatively optimal levels of sampling on loci and taxa, and robust nodal support on the tree), which contains 1290 loci, 22 samples with less than 30% missing data, and 8500 SNPs (Supplementary Table S4, available on Dryad).

The concatenated gene trees from both ML and BI methods showed a split of the EA and ENA species in two monophyletic sister clades with full support (Fig. 3a (1) and Supplementary Figs. S8a and S9a available on Dryad). Within the EA clade, *C. henryi* diverges off first, followed by *C. mollissima*, which is sister to the *C. crenata*-*C. seguinii* clade. Within the ENA clade, a sister relationship between *C. pumila* and *C. dentata* is strongly supported. The species trees from both ASTRAL-III and SVDquartets based showed the same relationships with strong support (Fig. 3c).

**Fig. 3.**
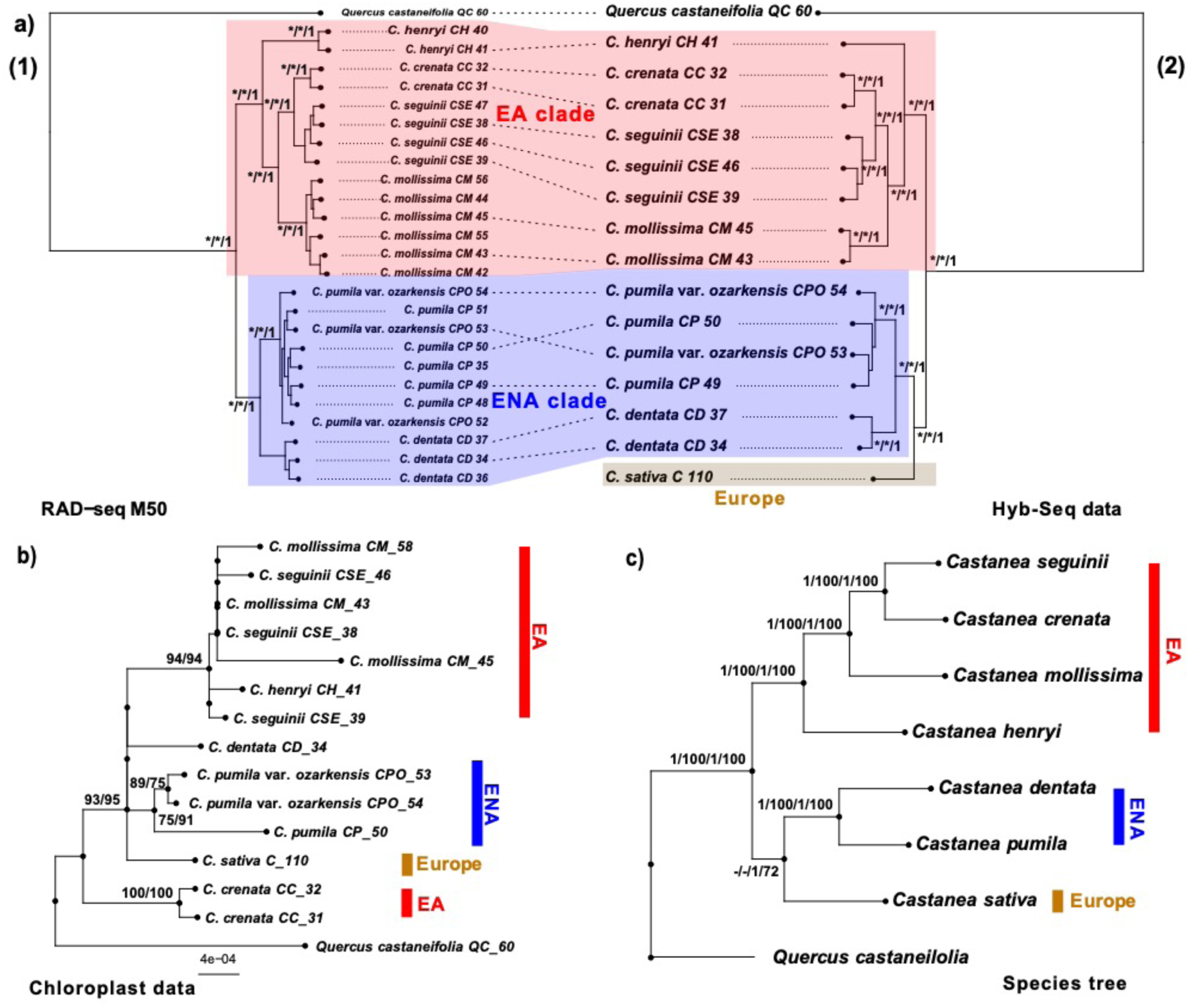
Gene trees and species tree of *Castanea* derived from different data sets. a) (1): the concatenated gene tree from RAD-seq M50 dataset; (2): the concatenated gene tree from well-trimmed “BWA100 *degenerated*” supercontig of orthologous gene data from Angiosperm 353 Hyb-Seq (primary Hyb-Seq-PPD data). Dash lines in the middle connecting the same samples used in both RAD-seq and Hyb-Seq analyses. Both concatenated gene trees in A were derived from analyses using the IQ-TREE whose topology is identical to those from RAxML and IQ-Tree. Numbers on branches are values of UF bootstrap from the IQ-TREE (only >90% numbers were shown), bootstrap values from RAxML (only >50% numbers were shown) (Supplementary Fig. S8, available on Dryad), and posterior probabilities from MrBayes (Supplementary Fig. S9, available on Dryad), respectively. Asterisks represent 100% support values. b) ML trees from 12,647 bp of plastid gene captured in the Hyb-Seq data by RAxML and IQ-Tree. Numbers on branches are combined from bootstrap in RAxML and ultrafasta bootstrap in IQ-Tree. c) The combined species tree from RAD-seq and Hyb-Seq data derived from analyses using ASTRAL-III and SVDQuartets. The numbers on the branches are support values from the RAD-seq data with ASTRAL-III, RAD-seq data with SVDQuartets, Hyb-Seq data with ASTRAL-III, and Hyb-Seq data with SVDQuartets, respectively. *Quercus castaneifolia* was used as the outgroup in all analysis. All trees were drawn by ggtree in R.

#### Castanea - Hyb-Seq

We were able to obtain Hyb-Seq data for one specimen of *C. sativa* so that all species of the genus were represented. The phylogenetic trees from ML and BI analyses of the primary Hyb-Seq-PPD dataset (see above) revealed the same relationships as shown in the RAD-seq trees, recognizing the reciprocal monophyly of the EA and ENA species, with high support (Fig. 3a (2) and Supplementary Figs. S8b and S9b available on Dryad). The European species *C. sativa* is placed as the sister to the American clade, with high support (Fig. 3a (2) and Supplementary Figs. S8b and S9b available on Dryad).

The species trees constructed from this data matrix using ASTRAL-III and SVDQuartets have identical topologies that depict the same species relationships as the concatenated gene tree, with strong support for all nodes except one. The node uniting *C. sativa* and the ENA species in the species tree from SVDQuartets is relatively weakly supported (72% - see Figure 3c).

The analysis of the combined RAD-seq and primary Hyb-Seq-PPD data (the RAD-Hyb-Seq data) matrix resulted in identical ML and species tree topologies, which were also identical to the results of both RAD-seq and the primary Hyb-Seq-PPD datasets (see above). The combined data based-species tree is also highly supported except for the node uniting *C. sativa* and the ENA clade (SVDQuartets 68%) (Supplementary Fig. S10, available on Dryad).

#### Castanea - chloroplast phylogeny

Analyses of the chloroplast data using RAxML and IQ-tree resulted in an incompletely resolved phylogeny with lower support (Fig. 3b). The trees placed one of the EA species *C. crenata* as the sister of a well-supported clade consisting of the remaining species (93% bootstrap value and 95% ultrafast bootstrap value), different from the nuclear data from the RAD-seq and Hyb-Seq (Figs. 3a and 3c). Within this large clade, the remaining EA species form a well-supported monophyletic group (94% bootstrap value and 94% ultrafast bootstrap value). However, the relationships among this EA subclade, *C. sativa* from Europe, and the ENA species (*C. pumila* and *C. dentata*) remain unresolved (Fig. 3b).

#### Hamamelis - RAD-seq

Similarly, we found little difference in tree topologies among the analyses of different matrices (M20, M30, …, M80) using IQ-Tree. These trees varied slightly in support values (most pronounced in the tree from M80) (Supplementary Figs. S11-S17, available on Dryad). The M50 matrix was chosen for the same reasons described above for *Castanea* for downstream analyses (Supplementary Table S4, available on Dryad). Analyses of the filtered M50 with RAxML and MrBayes resulted in well-resolved phylogenies of the genus (Fig. 4a (1) and Supplementary Figs. S18a and S19a, available on Dryad). In trees from all three methods, one of the two EA species - *H. mollis* - is shown as the sister to all other *Hamamelis*, and the other EA species - *H. japonica* - is strongly supported as the sister of the ENA clade in the RAxML, IQ-Tree and MrBayes trees (with rapid bootstrap value 100, ultrafast bootstrap value 100 and PP value 1, respectively). The monophyly of the ENA clade is strongly supported, within which *H. ovalis* is sister to *H. mexicana*. Species tree analyses resulted in similar topologies to the likelihood analyses, but with two notable exceptions. The species tree from ASTRAL showed a different sister-group relationship within the ENA clade (*H. ovalis* sister to *H. vernalis* – see Figure 4c), while the SVDQuartets analysis showed the two EA species as sister to one another, different from all other analyses of RAD-seq data performed, albeit with low support (Supplementary Fig. S20, available on Dryad).

**Fig. 4.**
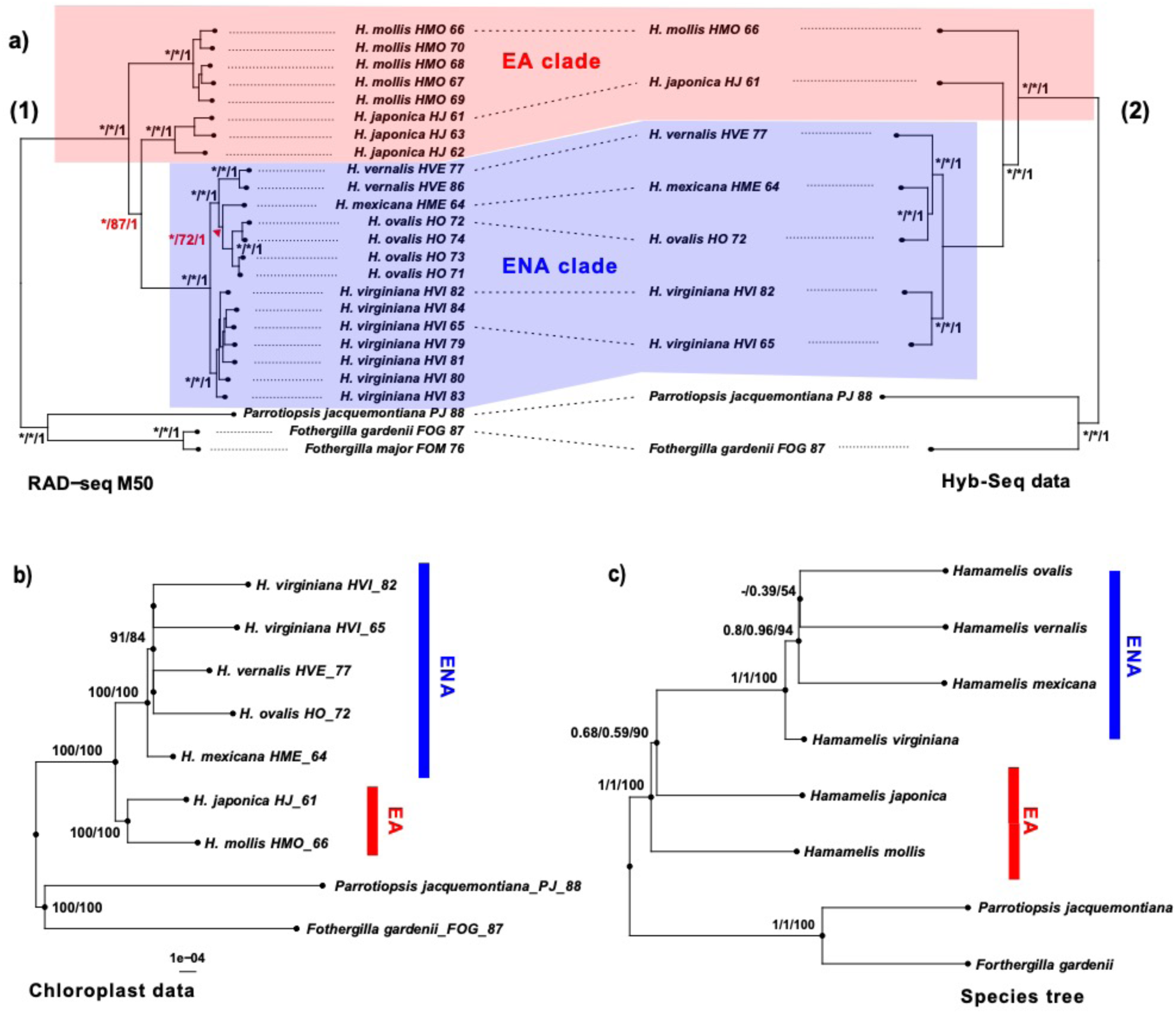
Gene trees and species tree of *Hamamelis* derived from different data sets. a) (1): the concatenated gene tree from RAD-seq M50 data; (2): the concatenated gene tree from the primary Hyb-Seq-PPD data. Dash lines in the middle connecting the same samples used in both RAD-seq and Hyb-Seq approaches. Both concatenated gene trees in A were derived from analyses using the IQ-TREE, whose topology is identical to those from RAxML and IQ-Tree. Numbers on branches are values of UF bootstrap from IQ-Tree (only >90% numbers were shown), bootstrap values from RAxML (only >50% numbers were shown) (Supplementary Fig. S18, available on Dryad), and posterior probabilities from MrBayes (Supplementary Fig. S19, available on Dryad), respectively. Asterisks represent 100% support values. b) ML tree from 107,068 bp of plastid gene from Hyb-Seq data derived from analyses using RAxML and IQ-Tree. Numbers on branches are bootstrap values from analyses using RAxML and IQ-Tree, respectively. c) The combined species tree from RAD-seq and Hyb-Seq data derived from analyses using ASTRAL and SVDQuartets. Numbers on the branches are support values from ASTRAL-III (RAD-seq), ASTRAL-III (Hyb-Seq), and SVDQuartets (Hyb-Seq), respectively. The species tree from RAD-seq data with SVDQuartets method shows a different species tree and is presented in Supplementary Fig. S20, available on Dryad. *Parrotiopsis jacquemontiana* and *Fothergilla gardenii* were used as outgroups. All trees were drawn by ggtree in R.

#### Hamamelis - Hyb-Seq

The primary Hyb-Seq-PPD data for *Hamamelis* contains only 160 genes. We increased the gene sampling to 319 genes by adding the additional genes with exon data. The new primary data consisted of 514,351 bp with 5,693 sites that are parsimony informative (Table 2). The trees resulting from analyses of this matrix using IQ-Tree, RAxML, and MrBayes are identical in topology (Fig. 4a (2) and Supplementary Figs. S18b and S19b, available on Dryad) and show species relationships identical to those resolved by the concatenated RAD-seq data (Fig. 4a (1)), with strong support. The two species trees reconstructed from ASTRAL and SVDQuartets are identical in topology (Fig. 4c), which is also identical to the concatenated Hyb-Seq gene tree and species tree inferred from the RAD-seq data with ASTRAL-III. Similarly, the node connecting the ENA clade and *H. japonica* is not well supported (0.59 in ASTRAL and 90 in SVDQuartets - see Figure 4c). The species trees inferred from the combined RAD-Hyb-Seq data were identical in topologies to trees derived from separate analyses of the RAD-seq and the primary Hyb-Seq-PPD dataset, but with higher support for the sister relationship between the ENA clade and *H. japonica* (0.99 in ASTRAL and 93 in SVDQuartets – see Supplementary Fig. S21, available on Dryad).

#### Hamamelis - chloroplast genes

Analyses of the chloroplast DNA data from RAxML and IQ-tree recovered a well-supported phylogeny similar to the species tree from RAD-seq data with SVDQuartets that recognized the monophyly of the EA species, but different from the nuclear phylogeny from other analyses as mentioned above that supported *H. japonica* and the ENA clade being sister. Relationships among the four ENA species remain unresolved in the plastid gene tree (Fig. 4b).

### Divergence Time Analyses

#### Castanea

Divergence time analyses of the RAD-seq data estimated the crown age of the genus (splitting of the EA and ENA clades) as the early Miocene (23.2 Ma, 95% HPD: 21.4-25.8 Ma). Within the genus, other divergence occurred in the mid-Miocene and late Miocene (Fig. 5a). The divergences times estimated with the primary Hyb-Seq-PPD data are slightly younger (∼0.6 to ∼5.3 Ma) on the mean values for all of the nodes than those estimated from the RAD-seq data (Fig. 5c; Supplementary Table S5, available on Dryad). The European chestnut (*C. sativa*), which was not missing in the RAD-seq data, diverged from the two ENA species in the mid-Miocene (13.6 Ma, 95% HPD: 10.9-16.7 Ma) based on the primary Hyb-Seq-PPD data (Fig. 5c). The divergence times estimated from the combined RAD-Hyb-Seq data were slightly older for the mean values than those based on the primary Hyb-Seq-PPD data (Supplementary Fig. S22a, available on Dryad) and slightly younger than the values from the RAD-seq data.

**Fig. 5.**
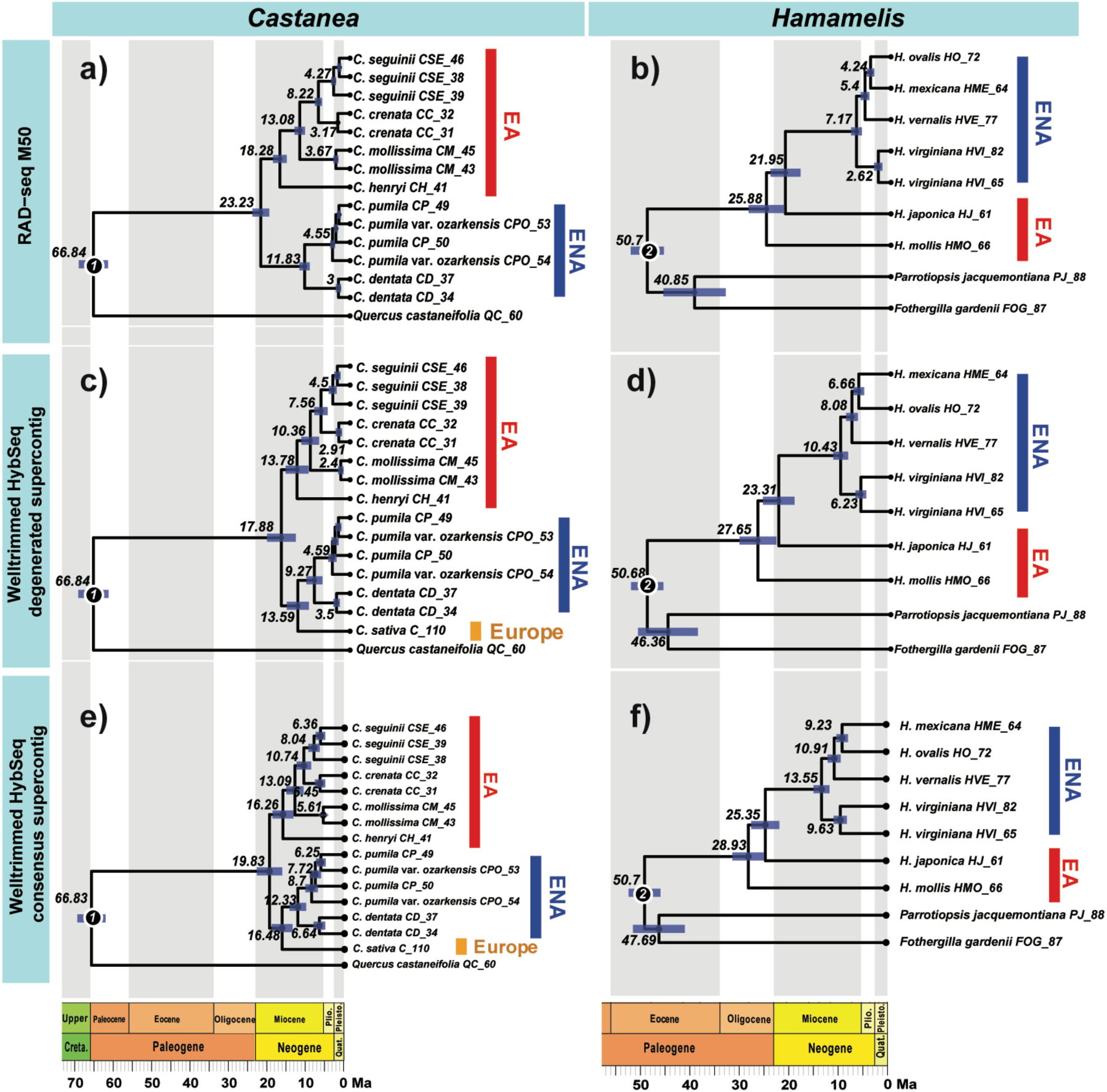
Results of divergence times estimated using RAD-seq and Hyb-Seq data. a) and b), results from RAD-seq M50 data; c) and d), results from the primary Hyb-Seq-PPD data. e) and f), results from well-trimmed “*consensus*” Hyb-Seq supercontig data. All topology was drawn by ggtree in R.

#### Hamamelis

Analyses of the RAD-seq data, Hyb-Seq-PPD data, and combined RAD-Hyb-Seq data resulted in similar estimates of divergence times (Figs. 5b and 5d; Supplementary Fig. S22b, available on Dryad). The crown node of *Hamamelis* (splitting of *H. mollis* from the remaining species) was dated back to the late Oligocene (e.g., 25.9 Ma with the 95% HPD as 22.6-30.1 Ma from the RAD-seq data; Fig. 5b). The divergence of *H. japonica* from the ENA clade was dated to the early Miocene (e.g., 22.0 Ma, with the 95% HPD as 19.1-25.3 Ma from the RAD-seq data; Fig. 5b). Divergence events within the American clade were dated to the late Miocene for *H. virginiana* and the Pliocene for the other species (Fig. 5b).

#### Comparison of Alignments, Phylogenies and Divergence Times among Different Hyb-Seq Matrices Phylogeny comparisons

Five comparisons of the phylogeny inferred from different Hyb-Seq data sets were conducted to evaluate the impacts of paralogous genes and the value of our new pipeline (Table 2). The results showed that the “*consensus*” matrix from the HybPiper pipeline and the various “*degenerated*” matrices from our PPD pipeline resulted in trees with the same topology (Table 2; Supplementary Fig. S23, available on Dryad), but the trees derived from the “*consensus*” matrices had longer branch lengths than the trees derived from the “*degenerated*” matrices (Supplementary Fig. S23, available on Dryad), probably due to the greater number of parsimony informative sites in the “*consensus*” matrices (Table 2). The results also showed that the phylogenies inferred from the putative paralogs differed from the orthologous gene phylogenies in terms of topology and branch lengths (Table 2; Supplementary Fig. S24, available on Dryad). Our results further showed congruence between the exon and supercontig trees in *Castanea* but minor incongruence between the exon and supercontig trees in *Hamamelis*. The exon tree grouped *H. mollis* and *H. japonica* as the sisters with low support (Supplementary Fig. S24, available on Dryad), a relationship not supported by analyses of the supercontig matrix (Figs. 4a (2) and 4c). Finally, our results showed that different levels of trimming of ortholog matrices did not influence topology, but did influence branch lengths (Supplementary Fig. S24, available on Dryad), whereas trimming paralogous gene alignments influenced both tree topologies and branch lengths (Supplementary Fig. S24 and details in Supplementary Information, available on Dryad).

#### Divergence time comparisons

Divergence time analyses on three data matrices (RAD-seq data, primary Hyb-Seq-PPD data (“*degenerated*”), and “*consensus*” Hyb-Seq data) resulted in some notable difference between matrices, but with mean estimates that were largely within the range of the HPDs of other estimates. The divergence times estimated from the “*consensus*” Hyb-Seq data were slightly older (∼2 - 4 Ma) at every node than those estimated from the primary Hyb-Seq-PPD data (Fig. 5; Supplementary Table S5, available on Dryad). The estimates based on the paralogs data identified by PPD were also older than those based on the orthologs and the 95% HPDs from the paralogs is also much larger in *Hamamelis* (Supplementary Fig. S25 and Fig. 5; Supplementary Table S5, available on Dryad). In contrast, the divergence times estimated from the exon and supercontig of orthologous genes are very similar at all nodes, with mean estimates overlapping HPDs of each analysis, although the exon based estimates are slightly older in *Castanea* and slightly younger in *Hamamelis*. Trimming the supertcontig matrix influenced divergence times estimates to some extent; the “auto-trimmed” matrix produced older (2∼6 Ma in *Castanea* and ∼2 Ma in *Hamamelis*) estimates than the “well-trimmed” matrix (Supplementary Fig. S25 and Fig. 5; Supplementary Table S5, available on Dryad), while the “untrimmed *consensus*” matrix (i.e., data directly from HybPiper) produced much older estimates than the two trimmed matrices (Supplementary Fig. S26 and Fig. 5; Supplementary Table S5, available on Dryad; Table 2), although these matrices supported the same species relationship.

### ILS and Introgression Tests

#### D-statistic tests

Our analyses found significant signals rejecting the null ILS hypothesis (D = 0) (P < 0.05) in both *Castanea* and *Hamamelis*. The results suggested that, in *Castanea*, introgression events occurred between the European species *C. sativa* (P2) and the EA clade (P3) (in both Hyb-Seq and the combined RAD-Hyb-Seq data: D = 0.3573, P < 0.0001) and occurred between the EA species *C. crenata* (P2) and ENA-*C. sativa* clade (P3) (Hyb-Seq: D = 0.5123, P < 0.0001; RAD-Hyb-Seq: D = 0.5135, P < 0.0001) (Table 4). In *Hamamelis*, the analyses found significant signals supporting introgression between *H. japonica* (as P3 or P2), and the American clade (as P2 or P3) in the RAD-seq data (P = 0.0015), in the primary Hyb-Seq-PPD data (P = 0.0009), and in the combined RAD-Hyb-Seq data (P < 0.0001; Table 4). However, the test did not find any significant signals supporting introgression among *H. mexicana*, *H. vernalis*, and *H. ovalis* in these three data sets (Table 4).

**Table 4.**
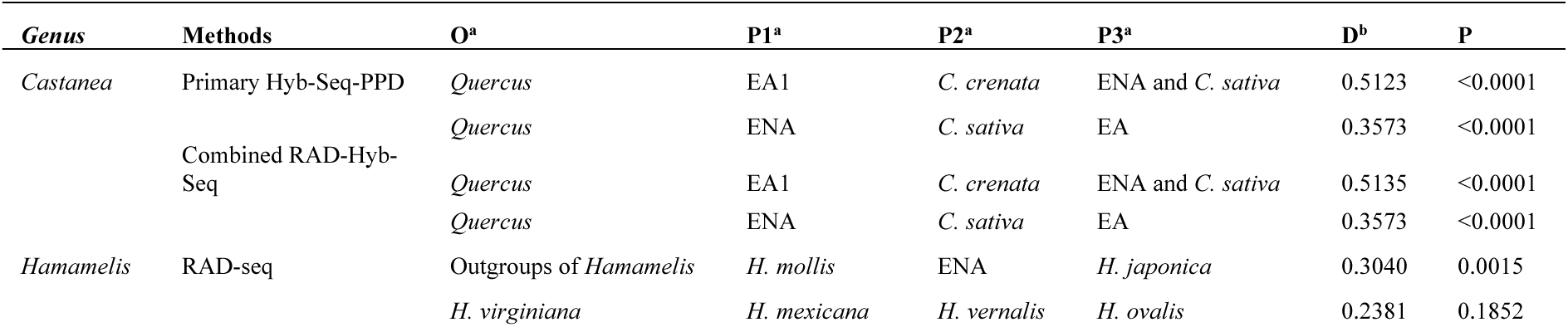
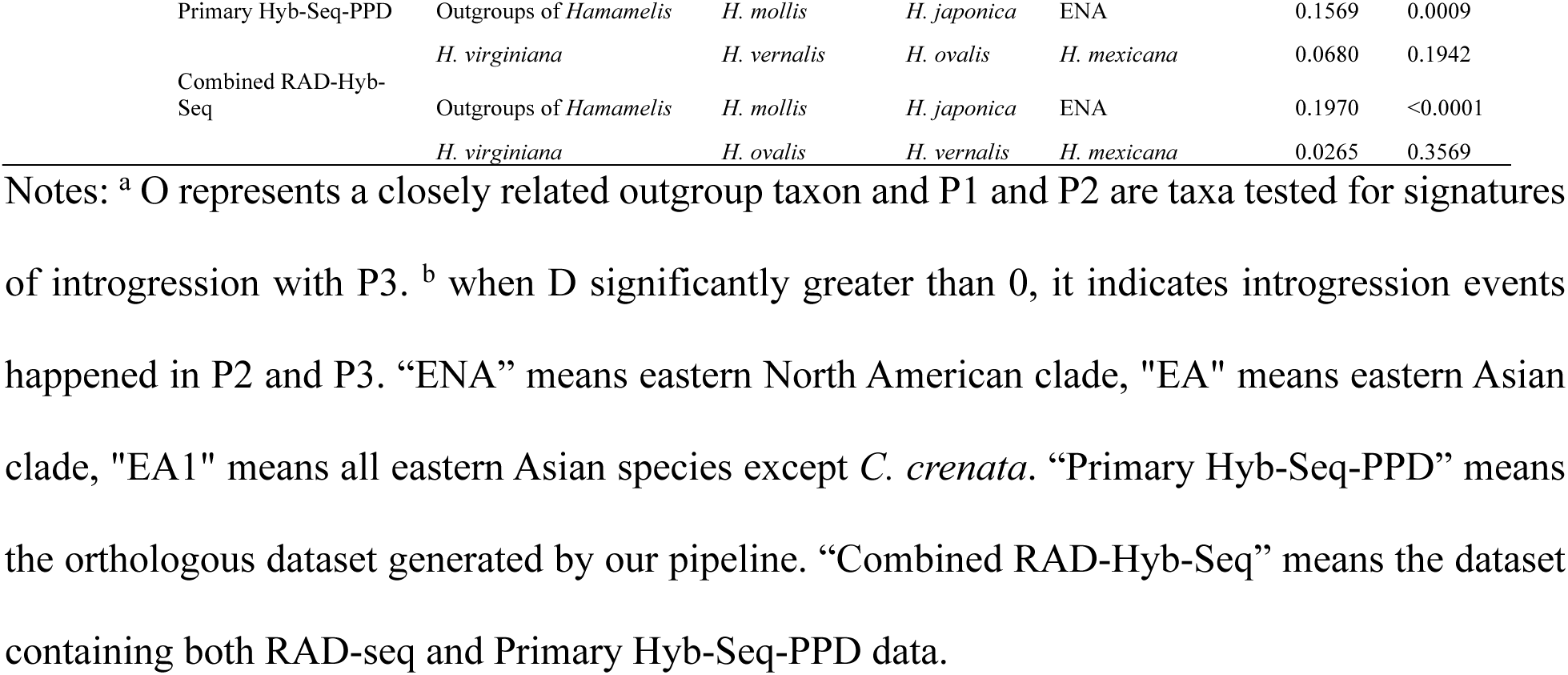
D-statistic (ABBA-BABA) result for testing introgression events in *Castanea* and *Hamamelis*.

#### Phylogenetic Network

The analyses with PhyloNet detected one reticulation event in *Castanea* between *C. sativa* and the stem of the EA clade excluding *C. henryi* (Fig. 6a). The inheritance probabilities show that *C. sativa* has 24% genomic contribution from the ancestor of EA clade excluding *C. henryi* (Fig. 6a). The analyses also detected one ancient reticulation event in *Hamamelis,* between *H. japonica* and the stem of the ENA clade. Inferred inheritance probabilities for this event indicate that the majority of the genomic contribution (65%) of the ENA clade came from its ancestor while a small proportion (35%) of the genomic contribution came from *H. japonica*. The tree topology from PhyloNet is similar to the chloroplast DNA tree showing the relationship of ((*H. mollis*, *H. japonica*), ENA clade) (Fig. 6b). The PhyloNet analyses did not detect the introgression event between *C. crenata* and ENA-*C. sativa* clade in *Castanea* suggested by the D-statistic test.

**Fig. 6.**
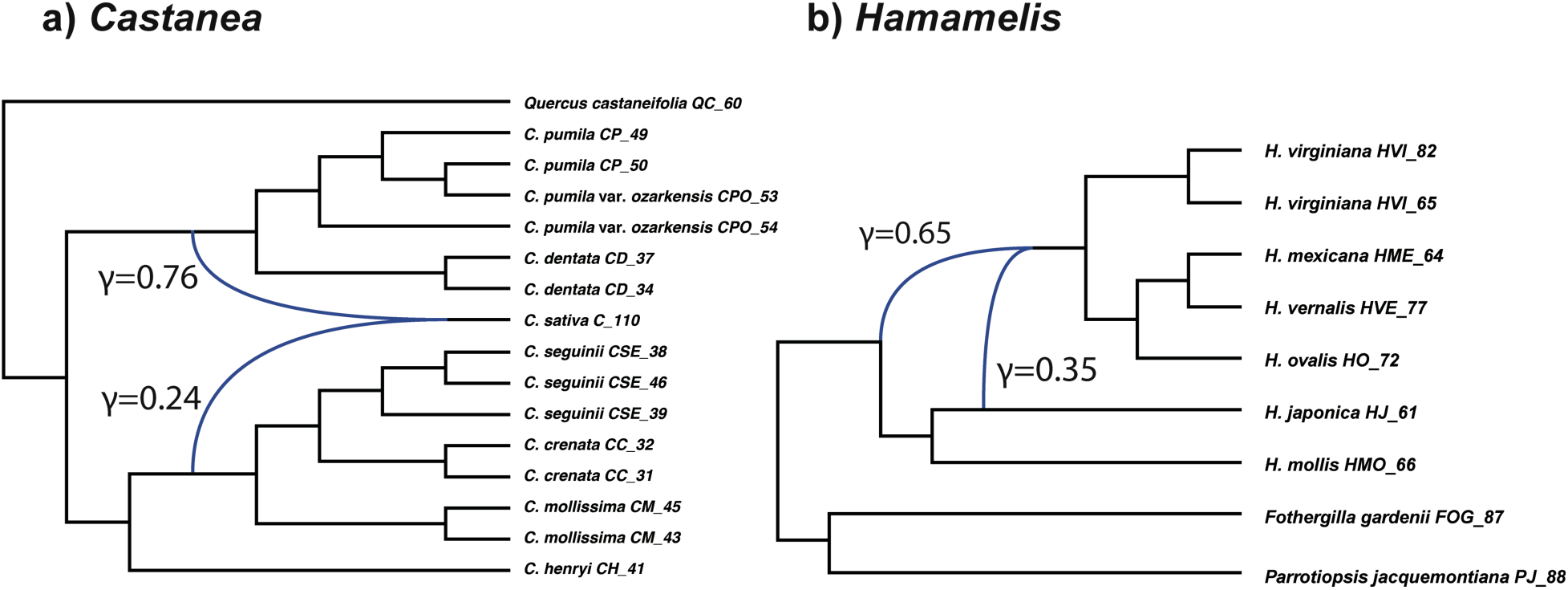
Best supported species network of the orthologous genes from primary Hyb-Seq-PPD data inferred with PhyloNet. Numbers next to the hybrid branches indicate inheritance probabilities.

### Biogeography of Castanea and Hamamelis

#### Castanea

The biogeographic analyses with the DEC method based on the dated total evidence phylogenies resolved the ancestral area of *Castanea* as western North America (“C”) in the Paleocene, where the genus diversified and expanded the range into eastern Asia (“AC”) during the Eocene period (Fig. 7a - combined RAD-Hyb-Seq and morphology data; Supplementary Fig. S27a - primary Hyb-Seq-PPD data and morphology data, available on Dryad), then followed by further diversification in the two areas. During the Oligocene, the genus expanded the range into Europe and eastern North America (“ABCE”) and diversified again, leading to the origin of the common ancestor of the modern *Castanea* species in Eurasia and eastern North America (“ABE”) in the Miocene. The other lineages evolved prior to the Oligocene in eastern Asia and western North America are all represented by fossil species and their origins involved extinctions in Asia, Western North America, and Europe from the Paleocene through the Oligocene (Fig. 7a and Supplementary Fig. S27a, available on Dryad). A vicariance of the common ancestor of the modern *Castanea* species in the early Miocene resulted in the EA (“A”) and Eur-ENA (“BE”) lineages. The EA lineage later diversified further into at least seven species in the mid and late Miocene with one of them spread into western North America. At least three of these species including the one that expanded its range into western North America became extinct in the late Miocene (Fig. 7a). The lineage is now represented by the four EA species. The Eur-ENA lineage (“BE”) became isolated between the two areas in the early-mid Miocene. The ENA part speciated into two extant ENA species *C. pumila* and *C. dentata* in the mid-late Miocene, while the European (“E”) part spread back to Asia and speciated further, followed by extinction in Asia in the early-mid Miocene. One of them survived as *C. sativa* in Europe and the other is represented by a fossil species *C. atavia* in Europe (Fig. 7a and Supplementary Fig. S27a, available on Dryad).

**Fig. 7.**
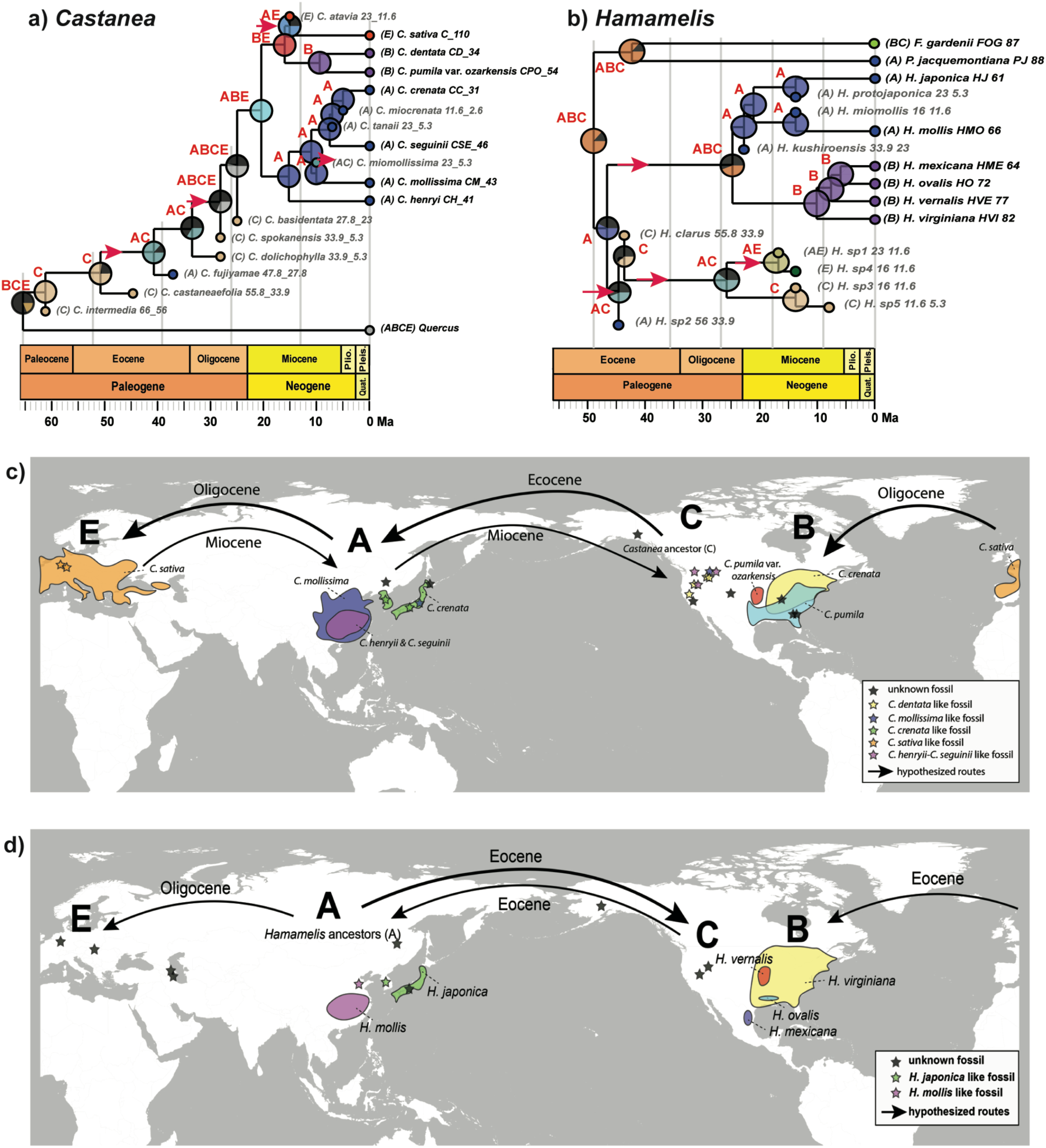
Results of biogeographic analyses of *Castanea* and *Hamamelis* using the DEC model based on dated phylogeny using total evidence with combined RAD-Hyb-Seq data and morphological data. a) DEC result of *Castanea*. b) DEC result of *Hamamelis*. Letters at each node indicate the most-likely ancestral distributions inferred from the analyses, whose probabilities are shown by non-black colors in the pie charts. The black color in the pie chart indicates the total proportion (likelihood) of all other alternative ancestral ranges in the results of DEC analyses. Inferred intercontinental dispersals are indicated by red arrows. Fossil taxa are labeled in gray color. c) Distribution and hypothesized migration routes of *Castanea* including living species and fossils. d) Distribution and hypothesized migration routes of *Hamamelis* including living species and fossils. A: eastern Asia; B: eastern North America and Mexico; C: western North America; E: Europe.

#### Hamamelis

The biogeographic analyses with the DEC method based on dated total evidence phylogeny of combined RAD-Hyb-Seq and morphology data resolved the ancestral area of *Hamamelis* as eastern Asia (“A”) in the mid-Eocene (47 Ma) (Fig. 7b - combined RAD-Hyb-Seq and morphology data; Supplementary Fig. S27b - primary Hyb-Seq-PPD data and morphology data, available on Dryad). The ancestor soon diverged into two lineages, one represented entirely by extinct fossil species, and the other containing both fossil and extant species. The lineage represented by fossil taxa spread into western North America (“C”) followed by subsequent diversification there, and dispersal back into Asia and Europe (“AC”). The other lineage spread to North America to obtain a widespread range in all three areas (“ABC”) before the Miocene and diverged into Asian and North American clades following the geographic isolation and extinction in western North America in the early Miocene (Fig. 7b and Supplementary Fig. S27b, available on Dryad). The Asian clade soon diversified into three lineages, with two surviving to present day. The North American clade diversified in the mid- and late-Miocene into four extant species of which one restricted to Mexico and the other three are restricted to ENA (“B”, Fig. 7b and Supplementary Fig. S27b, available on Dryad).

## DISCUSSION

### RAD-seq vs. Hyb-Seq Data

RAD-seq and targeted enrichment emerged to be two popular approaches in phylogenomic studies (e.g. Eaton and Ree 2013; Fernández-Mazuecos et al. 2018; Zhou et al. 2018; MacGuigan and Near 2019; Du et al. 2020 for RAD-seq data; Faircloth et al. 2013; McCormack et al. 2013; Leache et al. 2015; Léveillé-Bourret et al. 2018; Gaynor et al. 2020 for enrichment data), each with some notable advantages as discussed in the Introduction. We evaluated the two methods through side-by-side comparisons of phylogenomic results from data generated with RAD-seq and Hyb-Seq of Angiosperm 353 genes in the study of two EA-ENA disjunct genera, *Castanea* and *Hamamelis*. Our results showed, although there were fewer loci in the Angiosperm 353 gene data (e.g., 292 and 319 orthologs in well-trimmed “BWA100” datasets vs. 3,597 and 3,935 RAD-seq loci in the M20 matrices that contained the most RAD-seq loci of *Castanea* and *Hamamelis*, respectively) (Table 2 and Supplementary Table S4, available on Dryad), the total alignment length was much longer than that of the RAD-seq data owed to the longer average length of enrichment loci (Table 2). However, the proportion of the segregating sites in the Hyb-Seq data are similar to that of the RAD-seq data (Table 2). A similar pattern of differences in loci number, alignment length, and segregating sites between the RAD-seq and enrichment data was also reported in Harvey et al. (2016), but with lower values for these variables (the segregating sites: 4.07/site and 0.69% in the UCE matrices vs. 1.35/site and 1.41% in the RAD-seq matrices). However, the aforementioned differences in sequence data did not affect the topology recovered by phylogenetic methods in either genera (Figs. 3 and 4). Both of our RAD-seq and Hyb-Seq data well resolved the phylogenetic relationships within *Castanea* and *Hamamelis* with strong support (Figs. 3 and 4). This finding suggests that both approaches can generate a large amount of phylogenetic data for reliable inferences of species relationships within a genus. This is congruent with previous studies showing these two data sets performed similarly in phylogenetic inferences (e.g. Harvey et al. 2016; Manthey et al. 2016). However, the RAD-seq data may be unable to detect gene flow or introgression due to shortage of segregating sites per locus which decreases the chances of finding shared alleles on a very short phylogenetic branch (Harvey et al. 2016). We note that although both data supported the same species tree topology, the species tree may not be the correct species history, as found in our study of *Hamamelis* (see discussion below on introgression).

Although RAD-seq and Hyb-Seq data agreed in species relationships recovered from phylogenetic analyses, they appeared to slightly disagree in divergence times estimated using the same method. This result is unsurprising given the differences in alignment content and branch lengths in likelihood analyses. However, these differences had inconsistent effects across the two genera. The divergence times estimated from the Hyb-Seq data were younger (∼1-5 Ma) in *Castanea* but older (∼2-3 Ma) in *Hamamelis* at all nodes than those estimated from the RAD-seq data. Overall, the divergence time dating results are very similar between the RAD-seq and Hyb-Seq data (Fig. 5 and Supplementary Table S5, available on Dryad).

The phylogenetic congruence between data from RAD-seq and Hyb-Seq are encouraging and suggest that one may choose one or the other approach for a given phylogenomic study at the species level, according to their budget, labor and time available, conditions of DNA samples, and other goals. In general, the target enrichment approach can be effective with DNA samples at very low concentration or poor quality (Bi et al. 2012; Guschanski et al. 2013; McCormack et al. 2016), while the RAD-seq method requires DNA samples with relatively higher molecular weight. Although next-generation sequencing platforms have dramatically reduced the cost and time to obtain genome-wide phylogenetic markers (Glenn 2011; Wetterstrand 2015), funding and time may still be a limitation in large scale comparative studies (Harris et al. 2010). In such concerns, RAD-seq would be a better choice over Hyb-Seq.

It is noteworthy that our RAD-seq data captured a small number of the Angiosperm 353 genes included in the Hyb-Seq data (25 loci of RAD-seq matching to 21 loci of Hyb-Seq in *Castanea* and 13 loci of RAD-seq matching to 12 loci of Hyb-Seq in *Hamamelis*), indicating some level of overlap between the two datasets. This demonstrates that the RAD-seq data, as acquired using the experimental methods and analytical tools adopted in this study, likely represents a diverse array of genes and genomic regions, including the conserved low copy nuclear genes of angiosperms.

### Impacts of Paralogs in Hyb-Seq Data from HybPiper, Sequence Coding Method, and the Value of the PPD

Our results also showed that our new pipeline (PPD) identified many more putative paralogs than HybPiper. Although the “*consensus*” sequence data generated from HybPiper produced the phylogenetic tree with the same topology as the tree from the “*degenerated*” sequence data derived from PPD, the HybPiper data contained many more “false” phylogenetic informative sites (due to the presence of paralogous genes), resulting in longer branches affecting divergence time estimation (Fig. 5 and Supplementary Fig. S26 and Table S5, available on Dryad). The sequence data with better cleaning of paralogs and coded with the “*degenerated*” method are advantageous for phylogenomic studies, as they contain more accurate information for phylogenetic and divergence time estimations. Our analyses of the loci containing paralogous genes from the Hyb-Seq data often resulted in phylogenies different from those inferred from data of the orthologous genes (Supplementary Fig. S24, available on Dryad). The divergence times estimated from data including potential paralogous genes (i.e. the “*consensus*” data matrices from HypPiper) are likely older due to the additional variable sites introduced by gene paralogy (Supplementary Table S5, available on Dryad). Thus, our results clearly highlighted that paralogous gene content can negatively impact phylogenetic analyses. Our comparisons indicate that the PPD pipeline can effectively identify paralogs in the Hyb-Seq data and produce data with better quality for phylogenetic and divergence time dating analyses.

The advantage of “*degenerated*” (IUPAC consensus) coding of sequences from Hyb-Seq in phylogenetic studies has also been shown in previous studies. In Kates et al. (2018) the heterozygous sites were found to be disproportionately responsible for low resolution or topological uncertainty. In studies of Andermann et al. (2018b) and Lischer et al. (2014), although the IUPAC consensus matrix resulted in the same species tree as the “*consensus*” matrix, the divergence times estimated from the two data sets were different (underestimated divergence time in Andermann et al. 2018b and overestimated divergence time in Lischer et al. 2014). In our study, the “*consensus*” matrices resulted in long terminal branches (Supplementary Fig. S23, available on Dryad) and older divergence times at terminal nodes connecting species (dated to the mid-late Miocene in *Castanea* and the mid Miocene for the ENA species in *Hamamelis*) (Fig. 5). The longer branch lengths and older divergence times resulting from the “*consensus*” matrix (Fig. 5 and Supplementary Table S5, available on Dryad) are likely caused by mis-linkages of the most common SNPs within and among loci in the consensus matrix which resulted in inflated sequence variation among species.

In our PPD pipeline, we used the “*degenerated*” matrix using the IUPAC ambiguity codes instead of the allele phasing method because phasing many distant loci or phasing all heterozygous positions within loci is still not possible. Most allele phasing methods only selected the two most common alleles as the two phasing, which might not truly reflect their actual phases (Lischer et al. 2014; Kates et al. 2018). In addition, many commonly used phylogenetic software packages have options to treat ambiguity-coded positions as informative, including RAxML (Stamatakis 2014), IQ-Tree (Nguyen et al. 2015) for tree building, SVDQuartets (Chifman and Kubatko 2014) for species tree, and BEAST2 (Bouckaert et al. 2014) for divergence time estimation. Therefore, our new PPD pipeline produces alignments suitable for analysis across a wide range of modern phylogenetic tools.

### Influence of Gene Regions and Trimming Methods in Hyb-Seq Data

It is well-recognized that the rate and model of molecular evolution of genes and gene regions affect phylogenetic inferences. The sequence data from Hyb-Seq contain both exon and intron regions that evolve at strikingly different rates. How the exon and intron data would affect phylogenetic influences is worthy of exploration. Gene specific models of molecular evolution chosen from modeltest based on partitioned supercontig sequences may be less accurate, as they must account for the evolution of both the exon and intron regions in the same partition, which often evolve under different rates and models, and thus may not be accurate for either the exon or the intron regions within the same partition. The amount of phylogenetic information in the exon and intron regions also differs. How congruent are the analyses using the supercontig data with the approximate molecular models for the entire gene regions versus analyses using exon data alone with the more accurate models for exon regions remains to be explored. To gain insights into the question, we compared results from supercontig and exon data matrices constructed using the well-trimmed procedure and “*degenerated*” coding method.

Our results showed that in *Castanea*, the tree topologies inferred from the exon and supercontig data were identical, while in *Hamamelis*, the two datasets supported different relationships among *H. japonica*, *H. mollis*, and the ENA clade. The exon data supported a sister relationship between the two EA species *H. japonica* and *H. mollis*, with low support, consistent with the plastid data phylogeny where this relationship is strongly supported (Fig. 4 and Supplementary Fig. S24, available on Dryad), whereas the supercontig and RAD-seq data support a sister relationship between *H. japonica* and the ENA clade in all of the concatenated gene trees and species trees we inferred (Fig. 4). However, our PhyloNet result indicated that the relationships recovered by the exon data represented the true species history, while the relationship strongly supported by the supercontig data represents an ancient introgression (Fig. 6; see more details below). These results suggest that the information contained in exons can track a deeper true history, while the intron regions may track the more recent introgression history and be biased in species tree reconstruction, if the introgressed genomic contribution is high (35% in the case with *Hamamelis*; Fig. 6b). Our comparison between exon and supercontig data indicates the impact of rate of molecular evolution and suggests potential problems with the intron data from Hyb-seq and RAD-seq. We recommend vigorous exploration of datasets for more accurate reconstruction of phylogenetic histories of studied lineages.

Our comparisons among data sets trimmed differently by altering the matching length (k values in BWA implemented in our PPD pipeline) showed dramatic changes in the number of detected paralogs (48 vs. 116 in *Chastanea* and 27 vs. 106 in *Hamamelis* from k=19 to k=100 Table 2). Increasing the k value also resulted in the filtering of more mapping reads with low quality (matching length less than k). Therefore, we would like to suggest using our PPD pipeline with larger k value (100) in BWA and “*degenerated*” coding to obtain Hyb-Seq data in better quality for phylogenetic analyses.

### Biogeographic History of Castanea and Hamamelis and Insights into the EA-ENA Floristic Disjunction

The EA-ENA floristic disjunction represents a major phytogeographic pattern of in the Northern Hemisphere. Elements of the deciduous forests of the North Hemisphere exhibiting disjunct distributions in two or more areas (eastern Asia, eastern North America, western North America, and Europe) are often considered relics of the once widespread former mesophytic forests that continuously spanned the Northern Hemisphere during the late Paleogene and early Neogene (Wolfe 1980; Boufford and Spongberg 1983; Tiffney 1985a). The origin of the pattern is a major biogeographic puzzle of the earth biota which has attracted the attention of botanists for more than a century (Boufford and Spongberg 1983; Tiffney 1985a) and has been widely studied during the past 20 years (e.g., Wolfe 1980; Xiang et al. 1998a, 1998b, 2000; Wen 1999; Xiang and Soltis 2001; Donoghue and Smith 2004; Harris and Xiang 2009; Wen et al. 2010; Harris et al. 2013; Manos and Meireles 2015). Available evidence suggested the floristic disjunctions was the product of complex evolutionary processes including migration/dispersal, vicariance, extinction and speciation in different lineages at different times that were affected by geological and climatic changes in the past (Tiffney 1985a, 1985b; Wen 1999; Tiffney and Manchester 2001; Milne 2006; Manchester et al. 2009; Wen et al. 2010). However, the relative importance of these different processes in the formation of the phytogeographic pattern remains an open question and needs to be evaluated with evidence from individual disjunct taxa. Biogeographic studies of diverse disjunct genera integrating data from phylogeny, paleontology, and paleobotany are helpful to shed light on the question. Previous studies supported that the Bering land bridge (BLB) and the North Atlantic land bridge (NALB) were two important corridors for plant exchanges between North America and Eurasia in the spread of the disjunct taxa during the Cenozoic period, with the NALB being more important for the thermophilic taxa (in the Paleogene) while the BLB being more important for the temperate taxa (in the Neogene) (Wen et al. 2010). Furthermore, mega analysis of biogeographic histories of ∼100 lineages based on topologies of published phylogenies that mostly did not include fossils revealed bi-directional intercontinental dispersals, but “out of Asia” migration was more frequent (Donoghue and Smith 2004). These findings advanced our understanding of the biogeographic pattern but need to be updated with information from new studies employing better molecular data and more sophisticated analytical tools than what were available before. In our study of *Castanea* and *Hamamelis*, two of the many woody genera with living species isolated in EA and ENA (*Castanea* also in Europe) but many fossils in western North America (WNA) and Europe (Table 3), we set an example of such approach to provide new information for better understanding of the famous biogeographic pattern.

Our biogeographic analyses using the DEC method based on the dated total evidence tree suggests a western North American origin of *Castanea* in the Paleocene, with subsequent dispersals into Asia in the mid-Eocene and Europe-eastern North America in Oligocene (Fig. 7a). This result suggests a role of the BLB for the late Eocene dispersal into Asia and the role of the NALB for the late Oligocene dispersal into ENA and then into Europe. Paleontological evidence indicates that EA was physically connected to WNA by the BLB until ∼4.7 Ma (Tiffney and Manchester 2001; Wen et al. 2016; Graham 2018), while NALB connected ENA and Europe in the early Paleogene (up to early Eocene). However, previous authors hypothesized that dispersal and gene flow across the NALB may be possible after the early Eocene until the early Miocene via island hopping (Tiffney and Manchester 2001; Wen et al. 2016; Graham 2018). Our result also indicated that the most recent common ancestor of the extant species had a widespread range in EA, ENA, and Europe (ABE) in the early Miocene, supporting dispersal via NALB in the early Miocene. Our vigorous biogeographic analyses including fossils revealed a much more complicated history of the genus, involving Paleogene migration and extinctions prior to the origin of the present EA-ENA disjunct clade (Fig. 7a). Our study on *Castanea* also suggests that the present disjunct distribution of the genus in EA and ENA was a result of vicariance by geographic isolation and extinction and an “out of western North American” migration in the early history of the genus, in contrast to the major pattern of “out of Asia” migration detected in the previous mega-analysis (Donoghue and Smith 2004).

In *Hamamelis*, our study including fossils similarly revealed a more complicated biogeographic history of the genus than that of the extant taxa. An Asian distribution of the common ancestor of the genus in the early Eocene and a widespread ancestor of the extant lineage in Asia and North America in the late Oligocene supports “out of Asia” migration in the early history but a vicariance origin of the disjunct distribution of the modern taxa, respectively (Fig. 7b). In both genera, however, the divergence of the EA and ENA lineages occurred in the late Oligocene-Miocene time, agreeing with the major pattern found in previous studies (Donoghue and Smith 2004; Harris et al. 2013; reviewed in Wen et al. 2010). Our data support Paleogene dispersals via both the BLB and NALB and revealed many extinctions during the Eocene through Miocene in all four areas, especially more in EA and WNA in both genera (Fig. 7). Our data also indicate that *Castanea* and *Hamamelis* are relic taxa of the boreotropical flora evolved in the Paleocene and Eocene, respectively, and experienced a complicated biogeographical history in the past, whereas the modern species are descendants of Mesophytic Forests developed after the global climatic cooling in the early Oligocene (Tiffney 1985b).

### Evolutionary History and Ancient Introgression

In our study, both RAD-seq and Hyb-Seq data strongly supported the monophyly of each of *Castanea* and *Hamamelis*, and the monophyly of an EA species and the ENA species in each genus, respectively (Figs. 3 and 4). Both data also strongly supported the presence of ancient introgression between the European species *C. sativa* and the ancestor of EA clade after the divergence of *C. henryi* in *Castanea* and between *H. japonica* and the ancestor of the ENA clade of *Hamamelis* (Table 4, Figs. 6b and 7b). The ancestral ranges of *C. sativa* (“AE”) and EA clade (“A”) were overlapping in the Miocene and the introgression between the two lineages were possible. Morphologically, *C. sativa* does seem to show some intermediary between the EA and ENA clades (e.g. leaf size and pubescence on the abaxial surface of mature leaves). Our PhyloNet result revealed that the lineage leading to *Hamamelis japonica* contributed to 35% of the genome of the ENA clade (Fig. 6b). The age of *H. japonica* and its extinct ancestor fossil species (*H. protojaponica*) overlapped with the age of the ancestor of the ENA clade of *Hamamelis* in the mid-Miocene, lending evidence in agreement with the suggested introgression (Fig. 7b). Given the biogeographic history and ancestral range of the common ancestor of the two lineages (“ABC”), the divergence and subsequent introgression of these lineages likely occurred at the high latitudes of “ABC” along the BLB area where the lineages leading to *H. japonica* and the ENA clade were overlapping in range and hybridized before they became extinct in the BLB and western North America. A hybrid origin of *H. japonica* between the ancestor of *H. mollis* and the ancestor of the ENA lineage is also possible.

Ancient inter-clade or inter-species ingression have been also been uncovered in other lineages using the D-statistics method with enrichment data (Zarza et al. 2016; Alexander et al. 2017; Everson et al. 2019). However, the D-statistics method may be prone to false-positives due to model violations or outliers as could come from, e.g., undetected paralogy or even library construction artifacts (Lambert et al. 2019; Blair and Ané 2020). Our result similarly showed two introgression scenarios supported by D-statistics but not by PhyloNet (Table 4 and Fig. 6). Specifically, the D-statistic method suggested introgression between the eastern Asian *C. crenata* and the European *C. sativa*, and between *C. crenata* and the ENA clade, but these events were not revealed by PhyloNet (Fig. 6a). We suggest using D-statistics for an initial test, followed by analysis with PhyloNet for testing gene introgression for observed gene tree conflicts. PhyloNet infers phylogenetic networks with consideration of ILS (Yu et al. 2013) and is a more reliable approach than D-statistics alone.

In addition, our study of *Castanea* indicated that the plastid genome data extracted from the Hyb-Seq are valuable to reconstruct a more complete picture of the evolutionary history of a study group. By integrating evidence from the nuclear and plastid gene phylogenies we uncovered a chloroplast DNA capture event in *C. crenata* based on a strongly supported conflicting placement of the species between the plastid and nuclear gene trees (Fig. 3). Based on the plastid gene tree, the chloroplast DNA donor could be one of the extinct species of *Castanea*, such as *C. fujiyamae* that occurred in EA during 47.8 to 27.8 Ma (Fig. 6) or an outgroup taxon of the genus, such as *Castanopsis* which is closely related to *Castanea* and overlaps distribution with *C. crenata* in EA (Jisaburō et al. 1965).

Our phylogenetic data do not agree with the morphology-based classification scheme of three sections in *Castanea* (Sect. *Eucastanon*, Sect. *Balanocastanon*, and Sect. *Hypocastanon*) (Doye 1908; Johnson 1988). Our result indicated that Sect. *Eucastanon* that included *C. dentata*, *C. sativa*, *C. mollissima*, *C. seguinii*, and *C. crenata* is paraphyletic and the character of one nut per cupule in *C. pumila* (ENA) and *C. henryi* (EA) is homoplasious. The taxonomic status of the Allegheny chinkapin (*C. pumila*) and the Ozark chinkapin has been disputed (Johnson 1988; Nixon 1997). Johnson (1988) considered the Ozark chinkapin as a variety of *C. pumila*, while Nixon (1997) regarded it as a separate species *C. ozarkensis*. In our study, all individuals representing *C. pumila* including the Ozark chinkapins formed a monophyletic group sister to *C. dentata* with strong support. Therefore, our phylogenomic study do not support the recognition of *C. ozarkensis* as a distinct species. However, the hypothesis should be further tested with population level sampling of related taxa.

It is noteworthy that the species relationship revealed from PhyloNet is congruent with the chloroplast gene tree in grouping *H. japonica* and *H. mollis* as sisters (Fig. 4b). This relationship was revealed in the exon orthologous data (Supplementary Fig. S24, available on Dryad) and one of the species trees that was reconstructed by SVDquartets with the RAD-seq data (Supplementary Fig. S20, available on Dryad). A simulation study by Chou et al (2015) showed that when ILS is very high, ASTRAL performed better than SVDquartets in constructing species history, but when ILS is low SVD quartet performed better than ASTRAL. In a more recent simulation study, Long and Kubatko (2018) showed that when gene flow is present, ASTRAL is not consistent while SVDquartets performed well. Our results appears to agree with the finding of Long and Kubatko (2018) suggesting SVDquartets is more likely to recover the true species history with data from numerous gene loci (like the RAD-seq data) when there is high level of inter-species introgression as observed in *Hamamelis*. Nonetheless, in all species trees, the support for the sister taxon of the ENA clade was low, clearly indicating conflicts among gene trees. Our results further indicate that in addition to species tree methods, data type also affects phylogenetic inferences in the presence of introgression.

## SUPPLEMENTARY MATERIAL

Data available from the Dryad Digital Repository: https://doi.org/10.5061/dryad.2fqz612n3

Demultiplexed sequence data are available for download from the NCBI Sequence Read Archive (SRA) (BioProject PRJNA670453).

## ACKNOWLEDGEMENTS

We thank the Soltis lab at Florida Museum of Natural History, Fu lab at Zhejiang University, Gao lab at Kunming Institute of Botany, Chinese Academy of Sciences, JC Raulston Arboretum, Arnold Arboretum, University of Washington Botanic Gardens, Paul Jones at Sarah Duke Garden, and UNC herbarium for providing some leaf tissues, Jeffrey Thorne from North Carolina State University and Matt Johnson from Texas Tech University for discussion of the PPD pipeline. We also thank Steven Manchester and Kathleen Pigg for the discussion of *Castanea* and *Hamamelis* fossils.

## Figure Captions

Fig. S1-Fig. S7 Concatenated gene trees of *Castanea* from RAD-seq datasets. These figures show the phylogenetic trees derived from analyses of M20 to M80 datasets, respectively, using IQ-Tree. *Quercus castaneifolia* was used as an outgroup. All topologies are identical, showing two monophyletic groups, EA clade and ENA clade. Numbers on branches indicate values of UF bootstrap support.

Fig. S8 Concatenated gene tree of *Castanea* from analysis of a) RAD-seq M50 and b) primary Hyb-Seq-PPD data using RAxML. *Quercus castaneifolia* was used as an outgroup. Numbers on branches are values of bootstrap support.

Fig. S9 Concatenated gene tree of *Castanea* from analysis of a) RAD-seq M50 and b) primary Hyb-Seq-PPD data using MrBayes. *Quercus castaneifolia* was used as an outgroup. Numbers on branches are values of posterior probability.

Fig. S10 The species tree of *Castanea* from combined RAD-Hyb-Seq data derived from analyses using ASTRAL-III and SVDQuartets. The species tree result is congruent with the nuclear gene topology in Fig. 3. The numbers on the branches are support values from the ASTRAL-III and SVDQuartets, respectively. *Quercus castaneifolia* was used as an outgroup.

Fig. S11-Fig. S17 Concatenated gene trees of *Hamamelis* from analysis of RAD-seq data. These figures show the phylogenetic results from analyses of M20 to M80 datasets, respectively, using IQ-Tree. *Parrotiopsis jacquemontiana* and *Fothergilla major* were used as outgroups, but the trees were rerooted by the bybrid *Hamamelis* x *intermedia*. Numbers on branches are values of UF bootstrap support.

Fig. S18 Concatenated gene tree of *Hamamelis* from analysis of a) RAD-seq M50 and b) primary Hyb-Seq-PPD data using RAxML. *Parrotiopsis jacquemontiana* and *Fothergilla major* were used as outgroups. Numbers on branches are values of bootstrap support.

Fig. S19 Concatenated gene tree of *Hamamelis* from analysis of a) RAD-seq M50 and b) primary Hyb-Seq-PPD data using MrBayes. *Parrotiopsis jacquemontiana* and *Fothergilla major* were used as outgroups. Numbers on branches are values of posterior probability.

Fig. S20 The species tree of *Hamamelis* from analysis of RAD-seq M50 data using SVDQuartets. *Parrotiopsis jacquemontiana* and *Fothergilla major* were used as outgroups. The result shows that EA clade is a monophyly with relatively low support value 62.

Fig. S21 Species trees of *Hamamelis* from combined RAD-Hyb-Seq data derived from analyses using ASTRAL-III and SVDQuartets. Two species trees were identical except the ENA clade. The numbers on the branches are support values from the ASTRAL-III and SVDQuartets, respectively. The SVDQuartets result of ENA clade was shown in the insert. *Parrotiopsis jacquemontiana* and *Fothergilla gardenii* were used as outgroups.

Fig. S22 Divergence times resulting from well-trimmed “BWA100 degenerated” supercontig of orthologous genes from combined RAD-Hyb-Seq data of a) *Castanea* and b) *Hamamelis*. All results showed a similar divergence time results with that from degenerated superontig result from the primary Hyb-Seq data in Fig. 5.

Fig. S23 Comparisons of phylogenies derived from “*consensus*” and “*degenerated*” matrices of the Hyb-Seq data. The first two rows show the results from *Castanea* exon and supercontig regions and the last two rows show the results from *Hamamelis* exon and supercontig regions. The columns show the results from untrimmed “*consensus*”, well-trimmed “*consensus*”, and well-trimmed “*degenerated*”, respectively. The results indicated the tree topologies are identical in both “*consensus*” and “*degenerated*”, but “*consensus*” matrices have longer branch lengths than “*degenerated*” matrices. The ENA clades in both *Castanea* and *Hamamelis* are highlighted by blue color, EA clades are highlighted by read, and European clades are highlighted by yellow.

Fig. S24 Comparisons of phylogenies derived from *orthologs* vs. paralogs, exon vs. supercontig, matrices using different trimmed matrices and different mapping methods of the Hyb-Seq data. In orthologs and paralogs comparison, the result shows the paralogs in exon regions have different tree topologies with orthologs in both *Castanea* and *Hamamelis*. “well-trimmed” matrices have shorter branch length than “untrimmed” matrices (also shown in Table S5). “auto-trimmed” matrices have shortest branch lengths and different tree topologies in paralogs. The ENA clades in both *Castanea* and *Hamamelis* are highlighted by blue color, EA clades are highlighted by read, and European clades are highlighted by yellow.

Fig. S25 Divergence times resulting from “BWA100 degenerated” paralogous supercontig, “well-trimmed degenerated” exon, and “auto-trimmed degenerated” supercontig matrices in Hyb-Seq. All results showed a similar divergence time results with that from degenerated superontig result from Hyb-Seq in Fig. 5. But the “BWA100 degenerated” paralogous supercontig results show much older divergence times at shallow nodes (species level) (also shown in Table S5).

Fig. S26 Divergence times resulting from untrimmed Hyb-Seq “*consensus*” supercontig of orthologous genes of *Castanea* and *Hamamelis*. All results showed much older divergence time results with that from well-trimmed “*degenerated*” and “*consensus*” superontig results from Hyb-Seq data in Fig. 5.

Fig. S27 Results of biogeographic analyses of *Castanea* and *Hamamelis* using the DEC model based on Hyb-Seq data. a). Most likely ancestral ranges reconstructed for *Castanea* based on the total-evidence phylogeny with *Quercus castaneifolia* as outgroup. b). Most likely ancestral ranges reconstructed for *Hamamelis* based on the total-evidence phylogeny with *Parrotiopsis jacquemontiana* and *Fothergilla gardenii* as outgroups. A: eastern Asia; B: eastern North America and Mexico; C: western North America; E: Europe. The letters at each node indicate the most-likely ancestral distributions inferred from the analyses, whose probabilities are shown by non-black colors in the pie charts. The black color in the pie chart indicates the total proportion (likelihood) of all other alternative ancestral ranges in the results of DEC analyses. Intercontinental dispersal events inferred were indicated by red arrows. Fossil taxa are labeled in gray color.

