## Supplementary Figures for "A New Paralog Removal Pipeline Resolves Conflict between RAD-seq and Enrichment"

Fig. S1-Fig. S7 Concatenated gene trees of *Castanea* from RAD-seq datasets. These figures show the phylogenetic trees derived from analyses of M20 to M80 datasets, respectively, using IQ-Tree. *Quercus castaneifolia* was used as an outgroup. All topologies are identical, showing two monophyletic groups, EA clade and ENA clade. Numbers on branches indicate values of UF bootstrap support.

Fig. S1

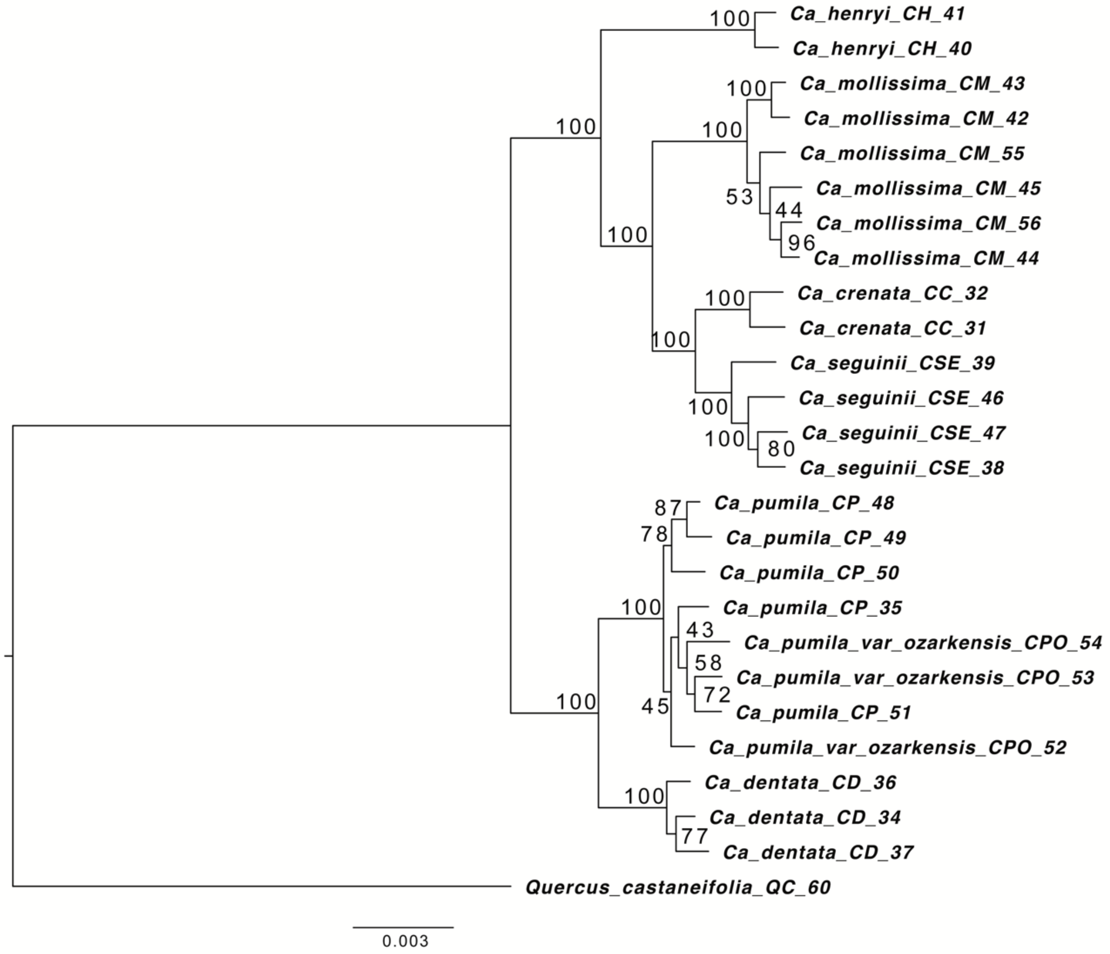

Fig. S2

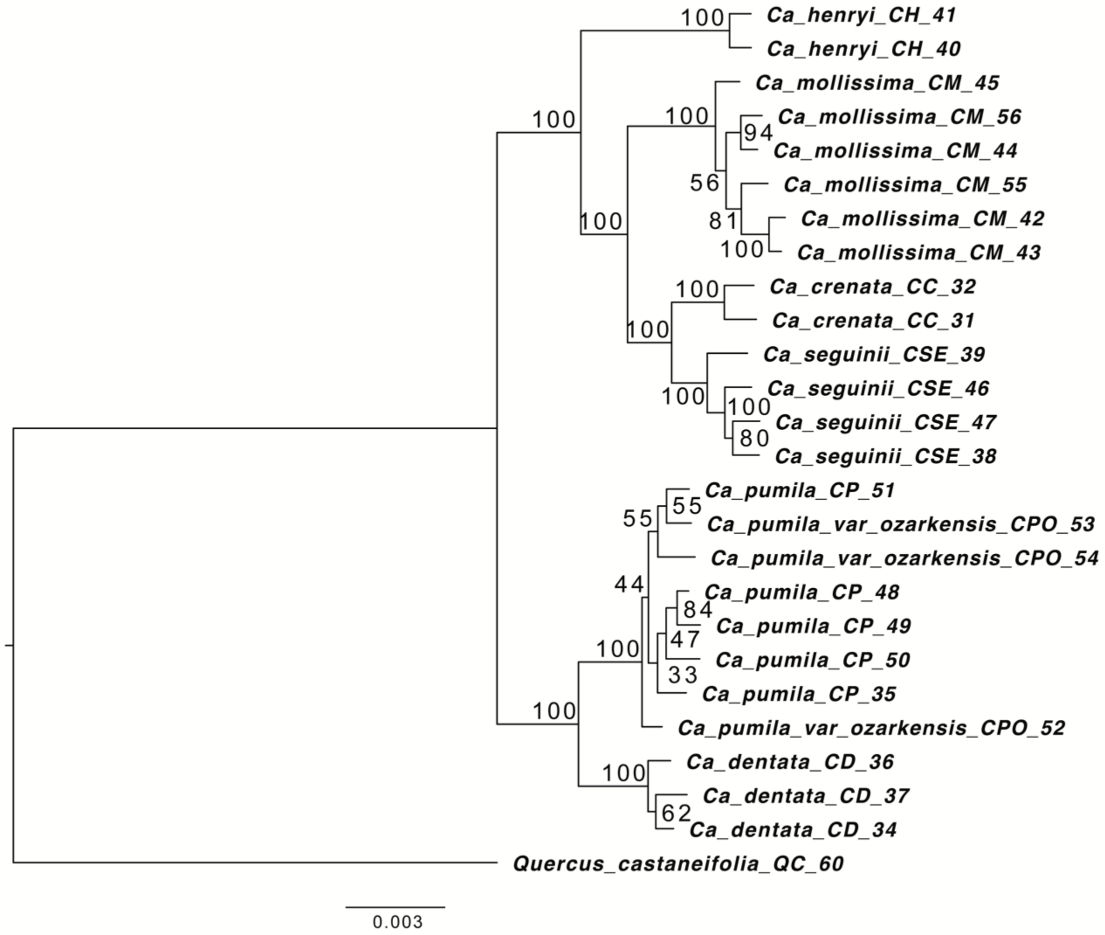

Ca_henryi_CH_41
Ca_henryi_CH_40
Ca_mollissima_CM_45
Ca_mollissima_CM_56
Ca_mollissima_CM_44
Ca_mollissima_CM_55
Ca_mollissima_CM_42
Ca_mollissima_CM_43
Ca_crenata_CC_32
Ca_crenata_CC_31
Ca_seguinii_CSE_39
Ca_seguinii_CSE_46
Ca_seguinii_CSE_47
Ca_seguinii_CSE_38
Ca_pumila_CP_51
Ca_pumila_var_ozarkensis_CPO_53
Ca_pumila_var_ozarkensis_CPO_54
Ca_pumila_CP_48
Ca_pumila_CP_49
Ca_pumila_CP_50
Ca_pumila_CP_35
Ca_pumila_var_ozarkensis_CPO_52
Ca_dentata_CD_36
Ca_dentata_CD_37
Ca_dentata_CD_34
Quercus_castaneifolia_QC_60
100
100
100
94
56
81
100
100
100
100
100
100
80
55
55
44
84
47
33
100
100
100
62
0.003

Fig. S3

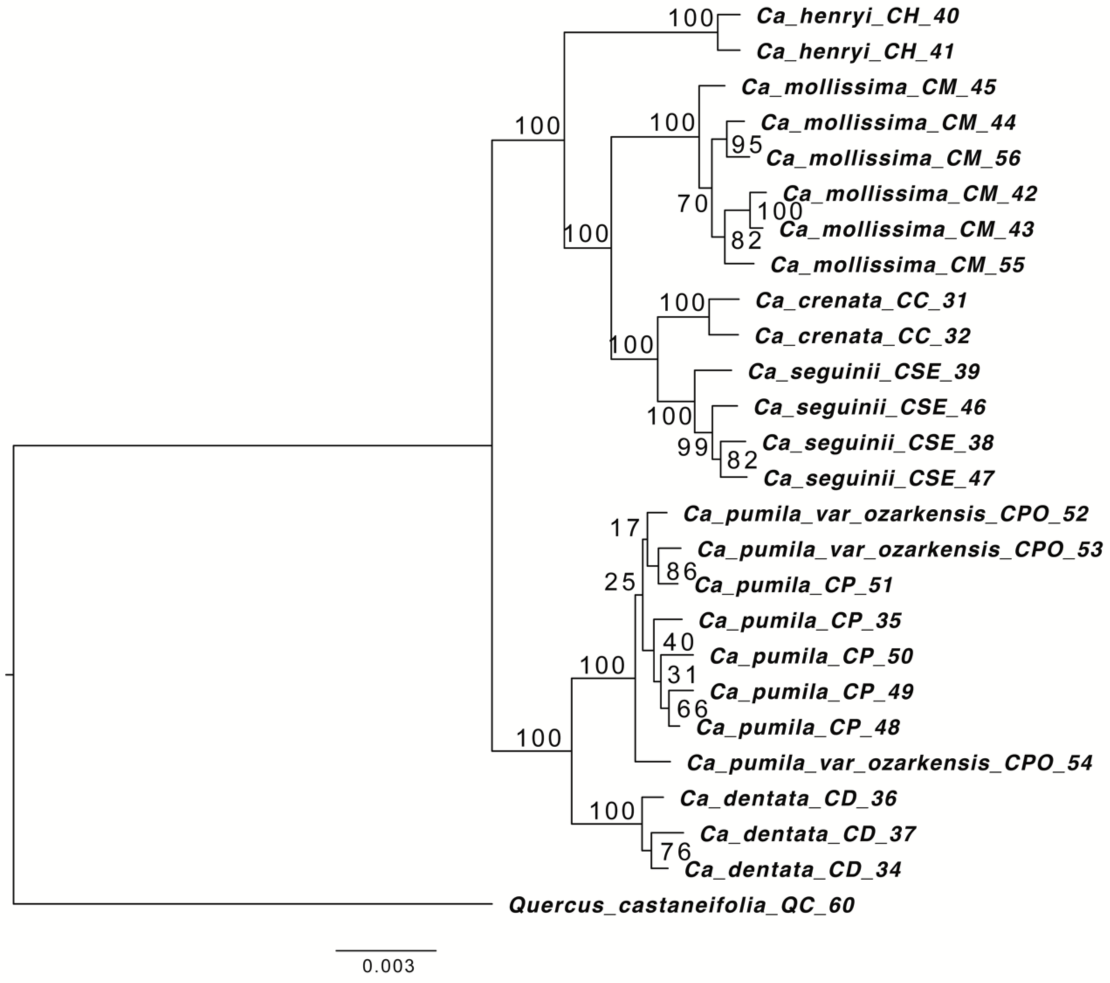
Ca_henryi_CH_40
Ca_henryi_CH_41
Ca_mollissima_CM_45
Ca_mollissima_CM_44
Ca_mollissima_CM_56
Ca_mollissima_CM_42
Ca_mollissima_CM_43
Ca_mollissima_CM_55
Ca_crenata_CC_31
Ca_crenata_CC_32
Ca_seguinii_CSE_39
Ca_seguinii_CSE_46
Ca_seguinii_CSE_38
Ca_seguinii_CSE_47
Ca_pumila_var_ozarkensis_CPO_52
Ca_pumila_var_ozarkensis_CPO_53
Ca_pumila_CP_51
Ca_pumila_CP_35
Ca_pumila_CP_50
Ca_pumila_CP_49
Ca_pumila_CP_48
Ca_pumila_var_ozarkensis_CPO_54
Ca_dentata_CD_36
Ca_dentata_CD_37
Ca_dentata_CD_34
Quercus_castaneifolia_QC_60
100
100
100
100
95
70
100
82
100
100
100
99
82
17
86
25
40
31
66
100
100
100
76
0.003

Fig. S4

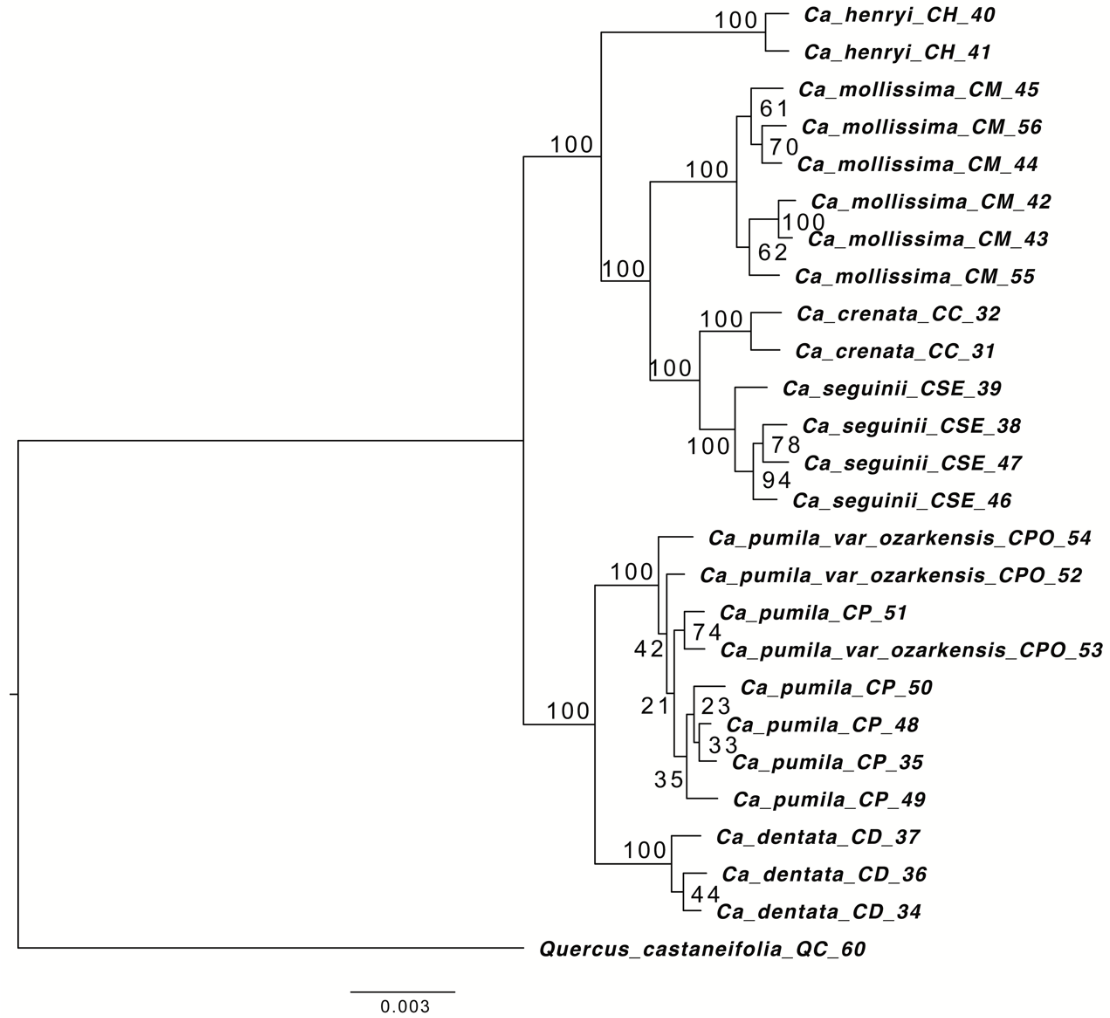

Ca_henryi_CH_40
Ca_henryi_CH_41
Ca_mollissima_CM_45
Ca_mollissima_CM_56
Ca_mollissima_CM_44
Ca_mollissima_CM_42
Ca_mollissima_CM_43
Ca_mollissima_CM_55
Ca_crenata_CC_32
Ca_crenata_CC_31
Ca_seguinii_CSE_39
Ca_seguinii_CSE_38
Ca_seguinii_CSE_47
Ca_seguinii_CSE_46
Ca_pumila_var_ozarkensis_CPO_54
Ca_pumila_var_ozarkensis_CPO_52
Ca_pumila_CP_51
Ca_pumila_var_ozarkensis_CPO_53
Ca_pumila_CP_50
Ca_pumila_CP_48
Ca_pumila_CP_35
Ca_pumila_CP_49
Ca_dentata_CD_37
Ca_dentata_CD_36
Ca_dentata_CD_34
Quercus_castaneifolia_QC_60
100
100
61
70
100
100
100
62
100
100
100
78
94
100
100
74
42
21
23
33
35
100
100
44
0.003

Fig. S5

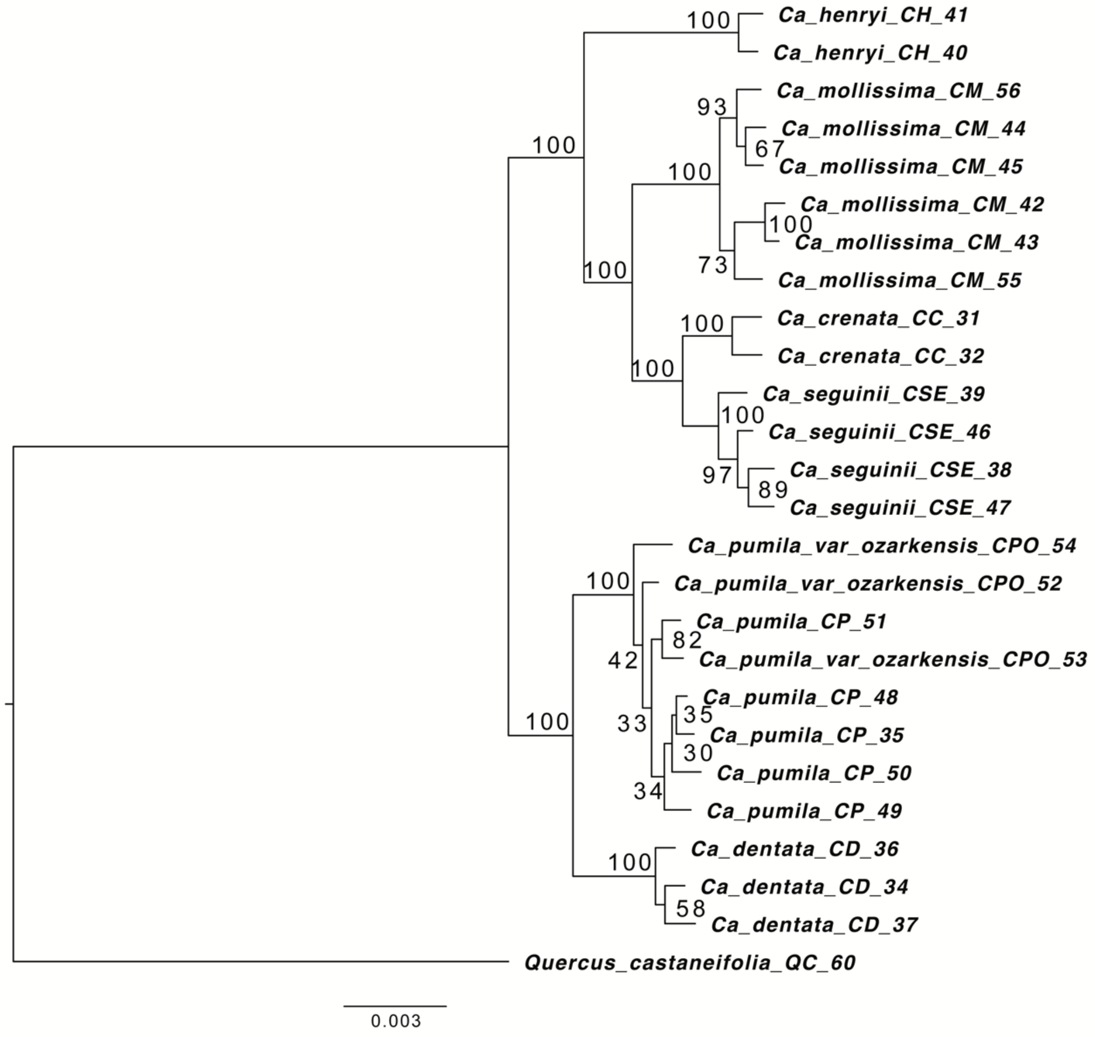
Ca_henryi_CH_41
Ca_henryi_CH_40
Ca_mollissima_CM_56
Ca_mollissima_CM_44
Ca_mollissima_CM_45
Ca_mollissima_CM_42
Ca_mollissima_CM_43
Ca_mollissima_CM_55
Ca_crenata_CC_31
Ca_crenata_CC_32
Ca_seguinii_CSE_39
Ca_seguinii_CSE_46
Ca_seguinii_CSE_38
Ca_seguinii_CSE_47
Ca_pumila_var_ozarkensis_CPO_54
Ca_pumila_var_ozarkensis_CPO_52
Ca_pumila_CP_51
Ca_pumila_var_ozarkensis_CPO_53
Ca_pumila_CP_48
Ca_pumila_CP_35
Ca_pumila_CP_50
Ca_pumila_CP_49
Ca_dentata_CD_36
Ca_dentata_CD_34
Ca_dentata_CD_37
Quercus_castaneifolia_QC_60
100
93
67
100
100
73
100
100
100
100
97
89
100
82
42
35
33
30
34
100
100
58
0.003

Fig. S6

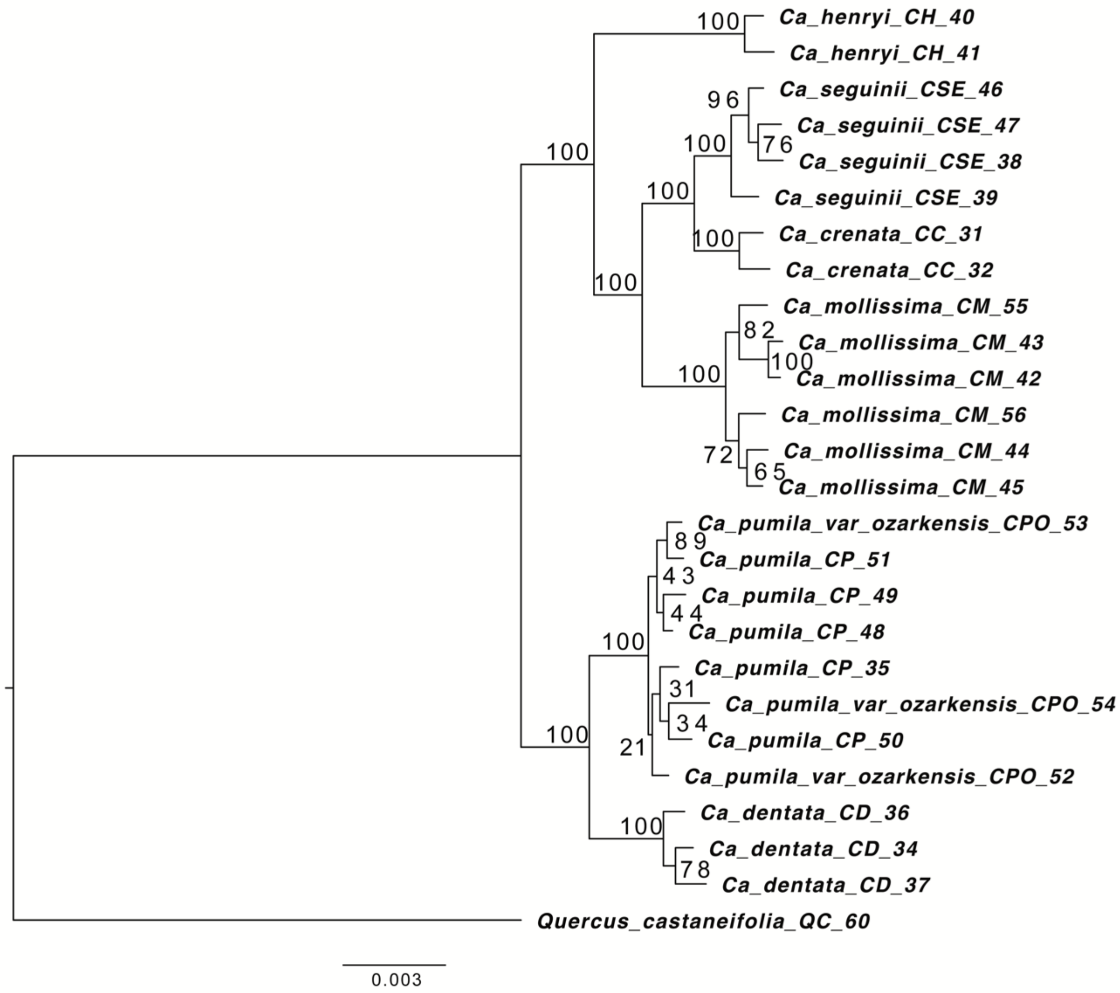
Ca_henryi_CH_40
Ca_henryi_CH_41
Ca_seguinii_CSE_46
Ca_seguinii_CSE_47
Ca_seguinii_CSE_38
Ca_seguinii_CSE_39
Ca_crenata_CC_31
Ca_crenata_CC_32
Ca_mollissima_CM_55
Ca_mollissima_CM_43
Ca_mollissima_CM_42
Ca_mollissima_CM_56
Ca_mollissima_CM_44
Ca_mollissima_CM_45
Ca_pumila_var_ozarkensis_CPO_53
Ca_pumila_CP_51
Ca_pumila_CP_49
Ca_pumila_CP_48
Ca_pumila_CP_35
Ca_pumila_var_ozarkensis_CPO_54
Ca_pumila_CP_50
Ca_pumila_var_ozarkensis_CPO_52
Ca_dentata_CD_36
Ca_dentata_CD_34
Ca_dentata_CD_37
Quercus_castaneifolia_QC_60
100
96
76
100
100
100
100
82
100
100
72
65
89
43
44
100
31
34
21
100
100
78
0.003

Fig. S7

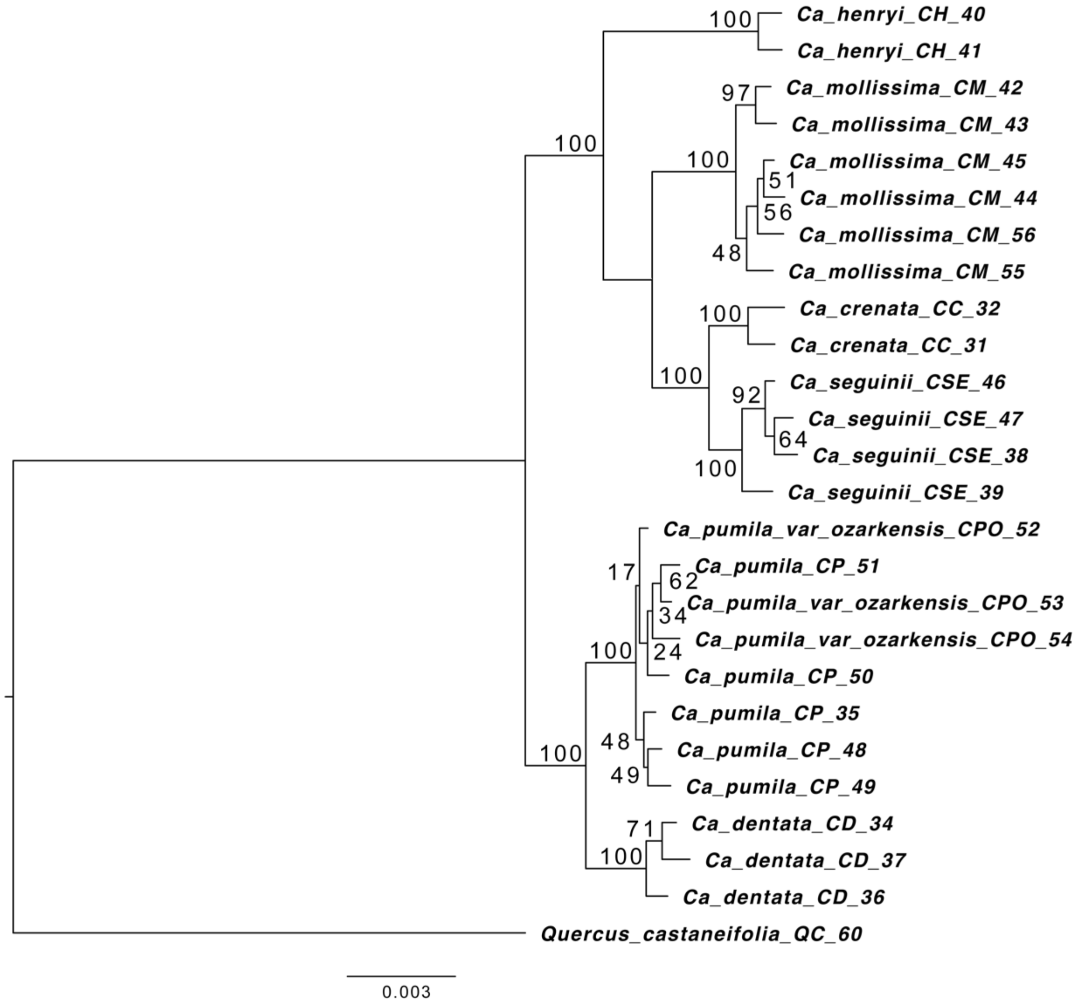
Ca_henryi_CH_40
Ca_henryi_CH_41
Ca_mollissima_CM_42
Ca_mollissima_CM_43
Ca_mollissima_CM_45
Ca_mollissima_CM_44
Ca_mollissima_CM_56
Ca_mollissima_CM_55
Ca_crenata_CC_32
Ca_crenata_CC_31
Ca_seguinii_CSE_46
Ca_seguinii_CSE_47
Ca_seguinii_CSE_38
Ca_seguinii_CSE_39
Ca_pumila_var_ozarkensis_CPO_52
Ca_pumila_CP_51
Ca_pumila_var_ozarkensis_CPO_53
Ca_pumila_var_ozarkensis_CPO_54
Ca_pumila_CP_50
Ca_pumila_CP_35
Ca_pumila_CP_48
Ca_pumila_CP_49
Ca_dentata_CD_34
Ca_dentata_CD_37
Ca_dentata_CD_36
Quercus_castaneifolia_QC_60
100
97
100
51
56
48
100
100
100
92
64
100
17
62
34
24
100
48
49
100
71
100
0.003

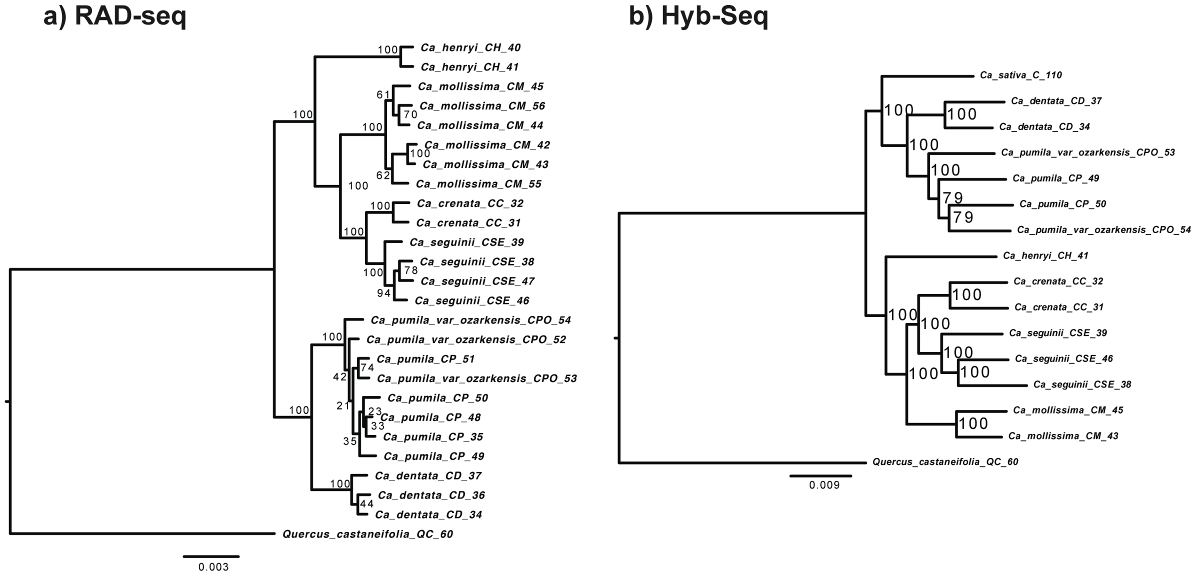


Fig. S8 Concatenated gene tree of *Castanea* from analysis of a) RAD-seq M50 and b) primary Hyb-Seq-PPD data using RAxML. *Quercus castaneifolia* was used as an outgroup. Numbers on branches are values of bootstrap support.

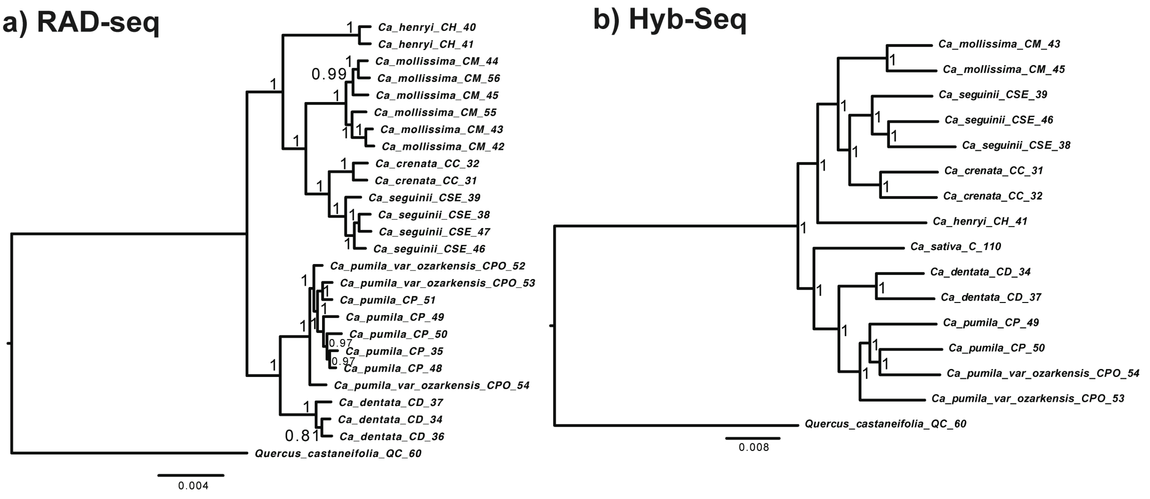


Fig. S9 Concatenated gene tree of *Castanea* from analysis of a) RAD-seq M50 and b) primary Hyb-Seq-PPD data using MrBayes. *Quercus castaneifolia* was used as an outgroup. Numbers on branches are values of posterior probability.

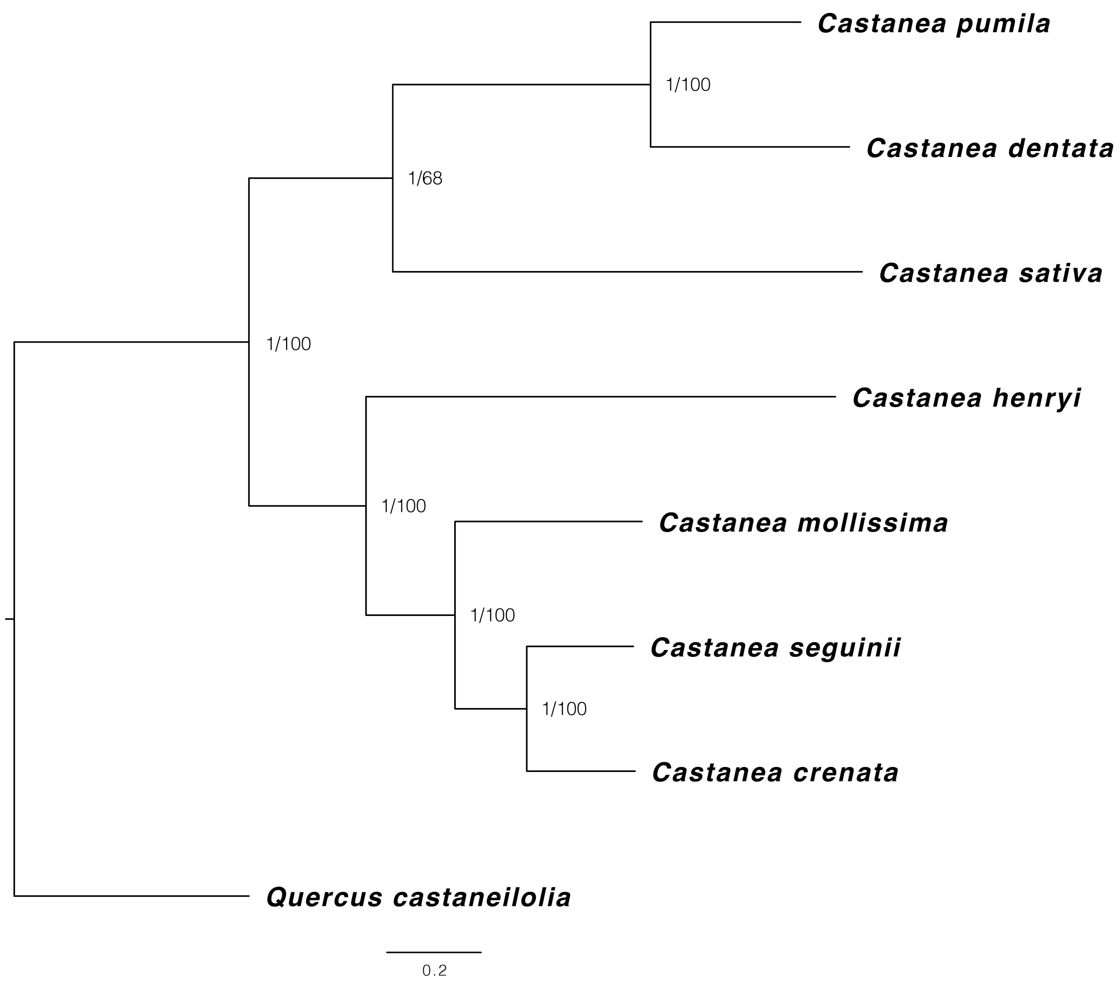


Fig. S10 The species tree of *Castanea* from combined RAD-Hyb-Seq data derived from analyses using ASTRAL-III and SVDQuartets. The species tree result is congruent with the nuclear gene topology in Fig. 3. The numbers on the branches are support values from the ASTRAL-III and SVDQuartets, respectively. *Quercus castaneifolia* was used as an outgroup.

Fig. S11-Fig. S17 Concatenated gene trees of *Hamamelis* from analysis of RAD-seq data. These figures show the phylogenetic results from analyses of M20 to M80 datasets, respectively, using IQ-Tree. *Parrotiopsis jacquemontiana* and *Fothergilla major* were used as outgroups, but the trees were rerooted by the bybrid *Hamamelis* x *intermedia*. Numbers on branches are values of UF bootstrap support.

Fig. S11

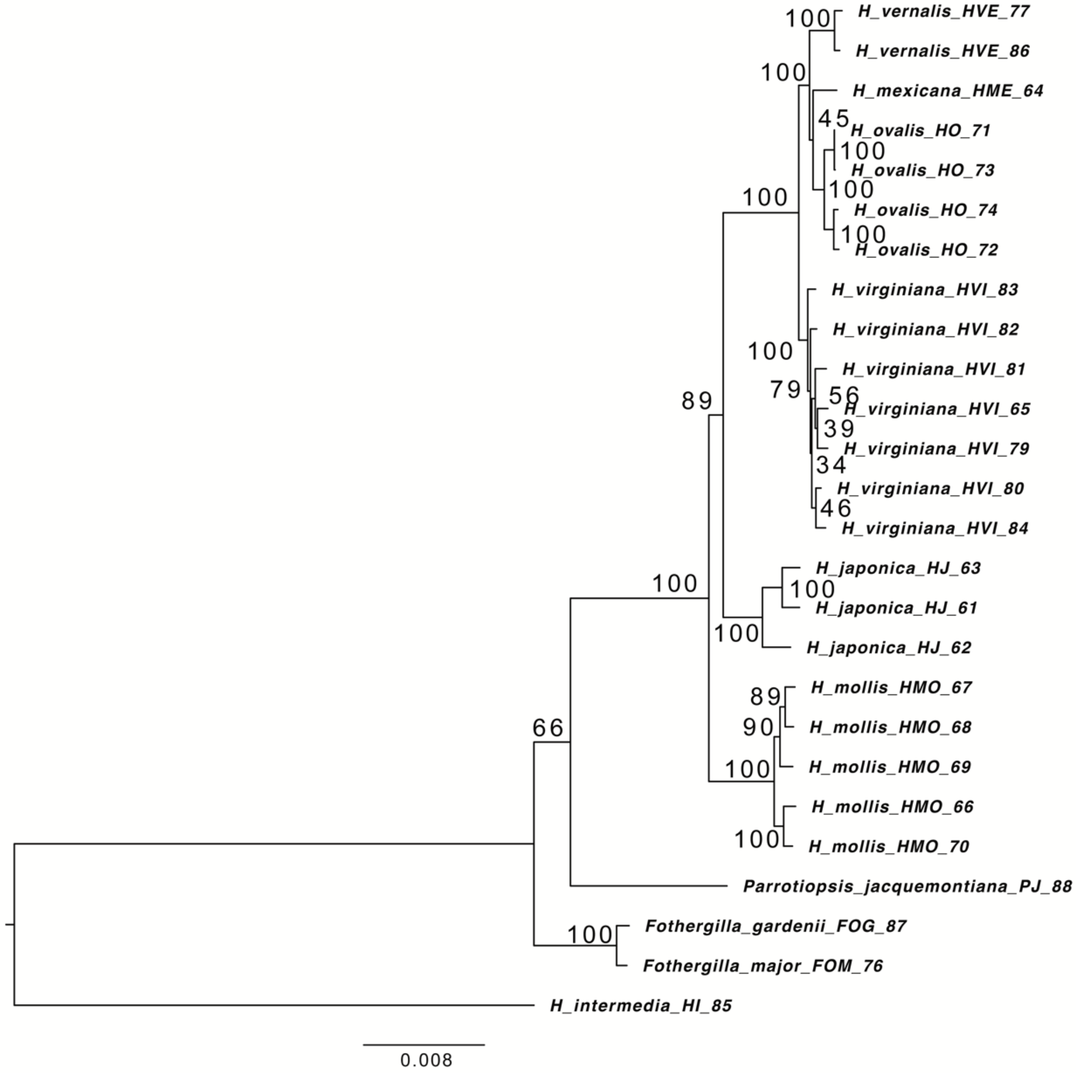

Fig. S12

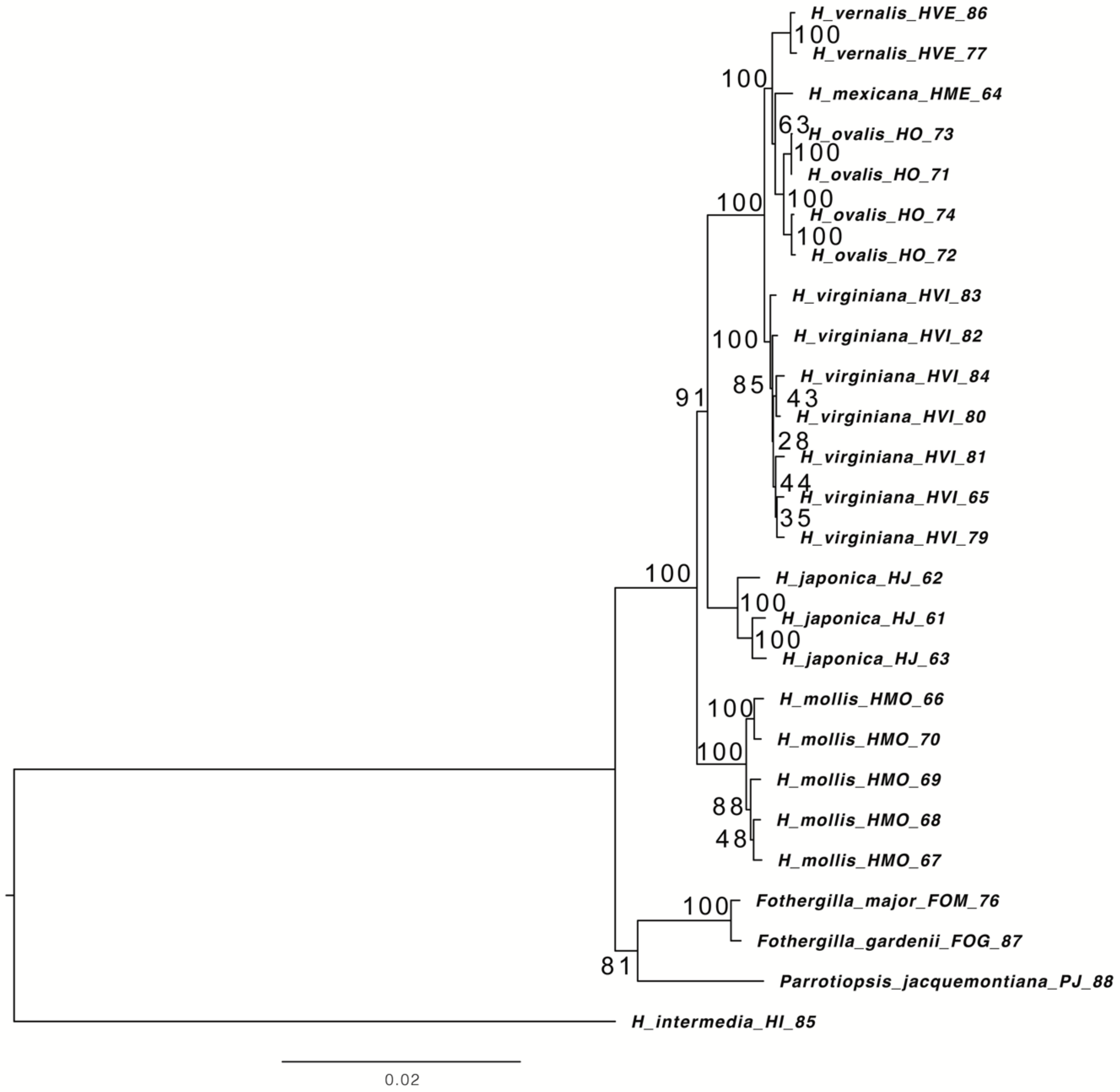
H_vernalis_HVE_86
100
H_vernalis_HVE_77
100
H_mexicana_HME_64
63
H_ovalis_HO_73
100
H_ovalis_HO_71
100
H_ovalis_HO_74
100
H_ovalis_HO_72
100
H_virginiana_HVI_83
100
H_virginiana_HVI_82
85
H_virginiana_HVI_84
43
H_virginiana_HVI_80
91
28
H_virginiana_HVI_81
44
H_virginiana_HVI_65
35
H_virginiana_HVI_79
100
H_japonica_HJ_62
100
H_japonica_HJ_61
100
H_japonica_HJ_63
100
H_mollis_HMO_66
H_mollis_HMO_70
100
H_mollis_HMO_69
88
H_mollis_HMO_68
48
H_mollis_HMO_67
100
Fothergilla_major_FOM_76
Fothergilla_gardenii_FOG_87
81
Parrotiopsis_jacquemontiana_PJ_88
H_intermedia_HI_85
0.02

Fig. S13

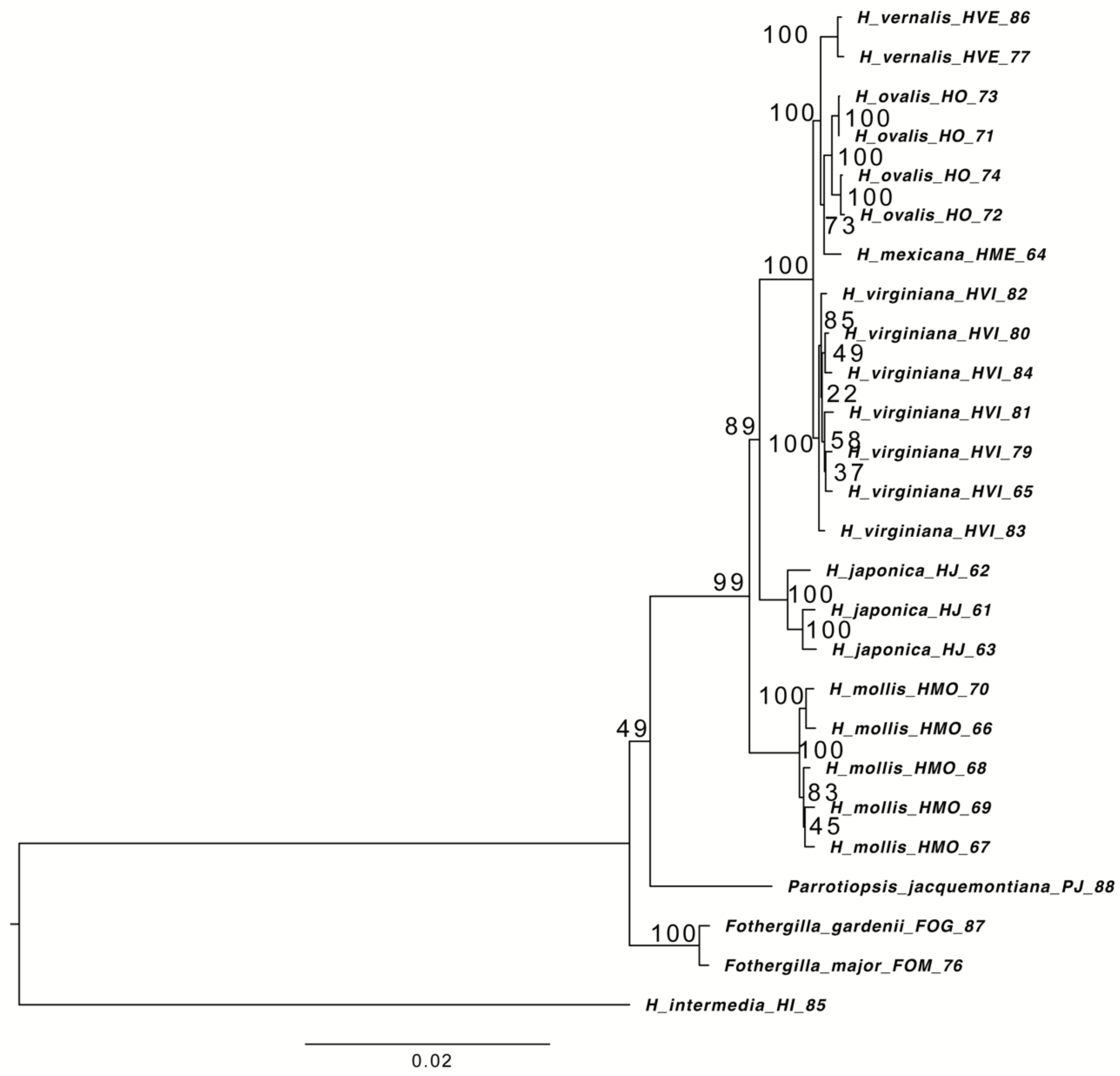

H_vernalis_HVE_86
H_vernalis_HVE_77
H_ovalis_HO_73
H_ovalis_HO_71
H_ovalis_HO_74
H_ovalis_HO_72
H_mexicana_HME_64
H_virginiana_HVI_82
H_virginiana_HVI_80
H_virginiana_HVI_84
H_virginiana_HVI_81
H_virginiana_HVI_79
H_virginiana_HVI_65
H_virginiana_HVI_83
H_japonica_HJ_62
H_japonica_HJ_61
H_japonica_HJ_63
H_mollis_HMO_70
H_mollis_HMO_66
H_mollis_HMO_68
H_mollis_HMO_69
H_mollis_HMO_67
Parrotiopsis_jacquemontiana_PJ_88
Fothergilla_gardenii_FOG_87
Fothergilla_major_FOM_76
H_intermedia_HI_85
0.02

Fig. S14

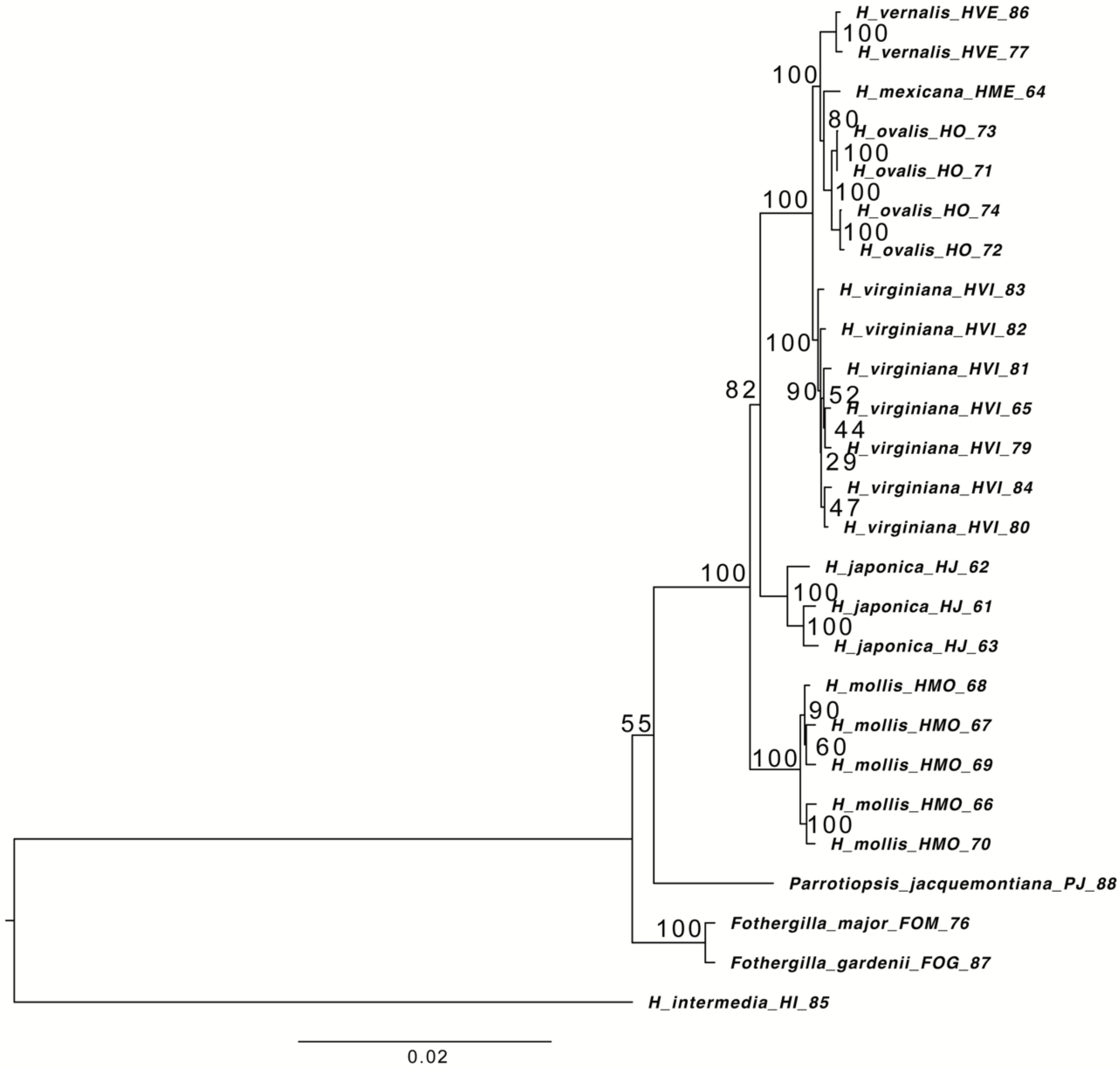
H_vernalis_HVE_86
H_vernalis_HVE_77
H_mexicana_HME_64
H_ovalis_HO_73
H_ovalis_HO_71
H_ovalis_HO_74
H_ovalis_HO_72
H_virginiana_HVI_83
H_virginiana_HVI_82
H_virginiana_HVI_81
H_virginiana_HVI_65
H_virginiana_HVI_79
H_virginiana_HVI_84
H_virginiana_HVI_80
H_japonica_HJ_62
H_japonica_HJ_61
H_japonica_HJ_63
H_mollis_HMO_68
H_mollis_HMO_67
H_mollis_HMO_69
H_mollis_HMO_66
H_mollis_HMO_70
Parrotiopsis_jacquemontiana_PJ_88
Fothergilla_major_FOM_76
Fothergilla_gardenii_FOG_87
H_intermedia_HI_85
100
100
80
100
100
100
100
100
82
90
52
44
29
47
100
100
100
90
60
100
100
55
100
0.02

Fig. S15

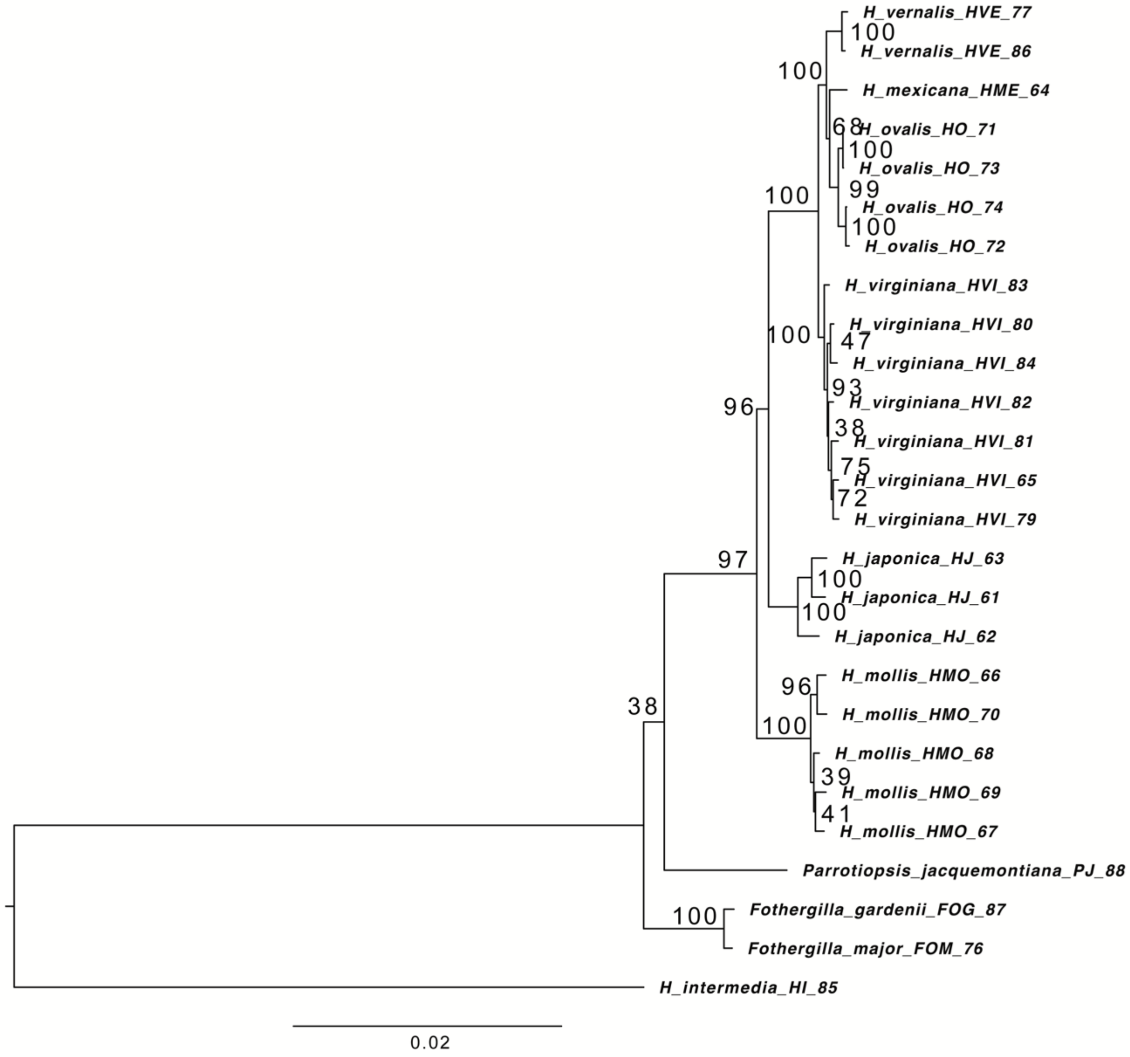
H_vernalis_HVE_77
100
H_vernalis_HVE_86
100
H_mexicana_HME_64
68
H_ovalis_HO_71
100
H_ovalis_HO_73
99
H_ovalis_HO_74
100
H_ovalis_HO_72
100
H_virginiana_HVI_83
100
H_virginiana_HVI_80
47
H_virginiana_HVI_84
93
H_virginiana_HVI_82
38
H_virginiana_HVI_81
75
H_virginiana_HVI_65
72
H_virginiana_HVI_79
96
97
H_japonica_HJ_63
100
H_japonica_HJ_61
100
H_japonica_HJ_62
96
H_mollis_HMO_66
H_mollis_HMO_70
100
H_mollis_HMO_68
39
H_mollis_HMO_69
41
H_mollis_HMO_67
38
Parrotiopsis_jacquemontiana_PJ_88
100
Fothergilla_gardenii_FOG_87
Fothergilla_major_FOM_76
H_intermedia_HI_85
0.02

Fig. S16

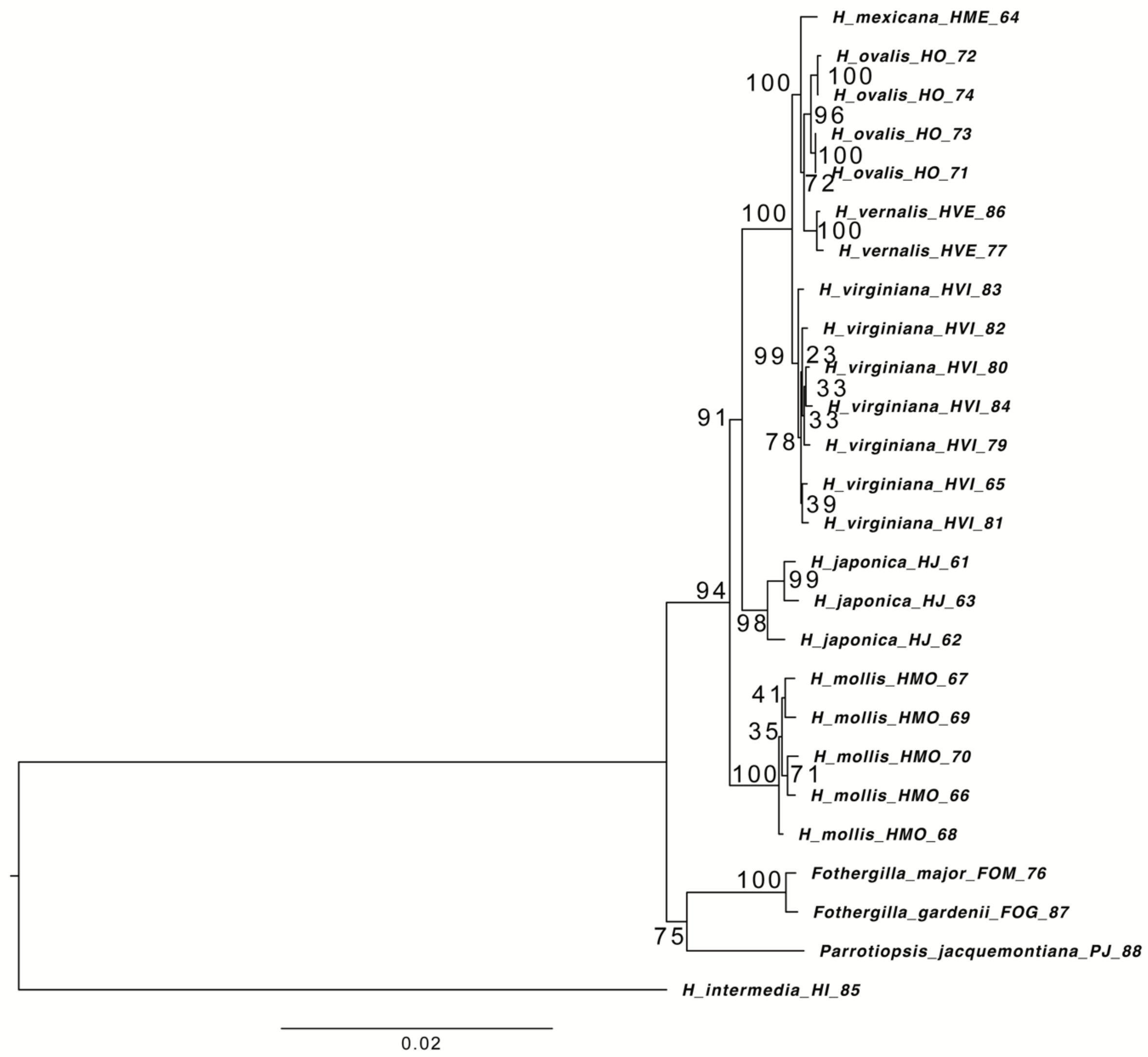

H_mexicana_HME_64
H_ovalis_HO_72
H_ovalis_HO_74
H_ovalis_HO_73
H_ovalis_HO_71
H_vernalis_HVE_86
H_vernalis_HVE_77
H_virginiana_HVI_83
H_virginiana_HVI_82
H_virginiana_HVI_80
H_virginiana_HVI_84
H_virginiana_HVI_79
H_virginiana_HVI_65
H_virginiana_HVI_81
H_japonica_HJ_61
H_japonica_HJ_63
H_japonica_HJ_62
H_mollis_HMO_67
H_mollis_HMO_69
H_mollis_HMO_70
H_mollis_HMO_66
H_mollis_HMO_68
Fothergilla_major_FOM_76
Fothergilla_gardenii_FOG_87
Parrotiopsis_jacquemontiana_PJ_88
H_intermedia_HI_85
100
96
100
72
100
100
100
99
23
33
33
78
39
91
99
98
94
41
35
71
100
100
75
0.02

Fig. S17

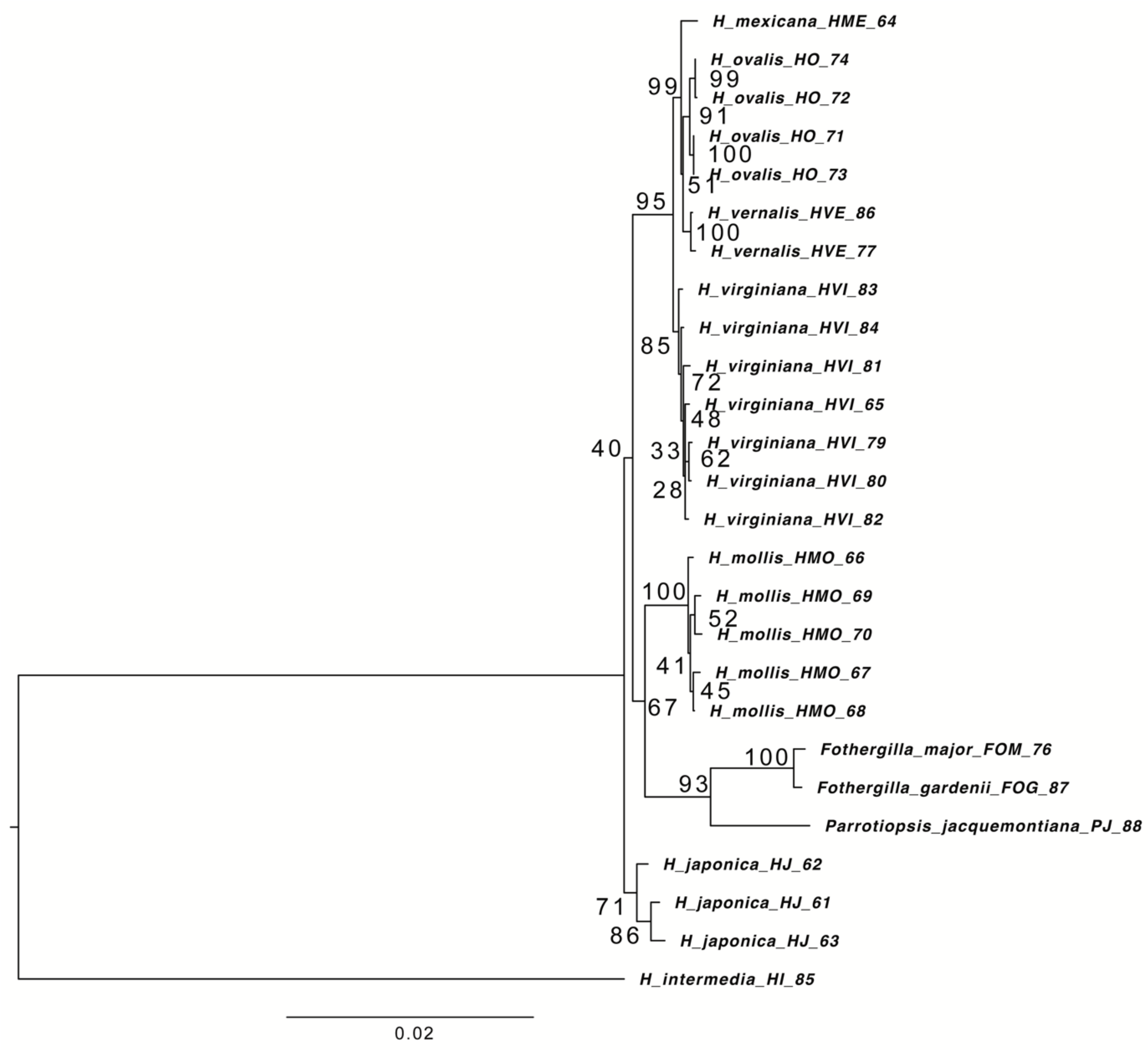
H_mexicana_HME_64
H_ovalis_HO_74
H_ovalis_HO_72
H_ovalis_HO_71
H_ovalis_HO_73
H_vernalis_HVE_86
H_vernalis_HVE_77
H_virginiana_HVI_83
H_virginiana_HVI_84
H_virginiana_HVI_81
H_virginiana_HVI_65
H_virginiana_HVI_79
H_virginiana_HVI_80
H_virginiana_HVI_82
H_mollis_HMO_66
H_mollis_HMO_69
H_mollis_HMO_70
H_mollis_HMO_67
H_mollis_HMO_68
Fothergilla_major_FOM_76
Fothergilla_gardenii_FOG_87
Parrotiopsis_jacquemontiana_PJ_88
H_japonica_HJ_62
H_japonica_HJ_61
H_japonica_HJ_63
H_intermedia_HI_85
99
99
91
100
51
95
100
85
72
48
40
33
62
28
100
52
41
45
67
100
93
71
86
0.02

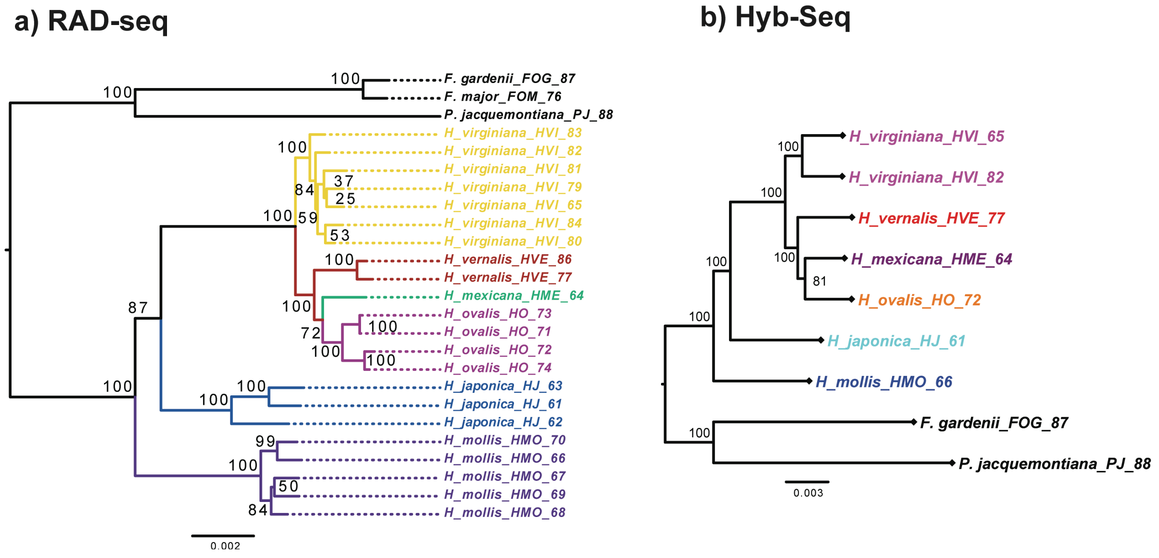


Fig. S18 Concatenated gene tree of *Hamamelis* from analysis of a) RAD-seq M50 and b) primary Hyb-Seq-PPD data using RAxML. *Parrotiopsis jacquemontiana* and *Fothergilla major* were used as outgroups. Numbers on branches are values of bootstrap support.

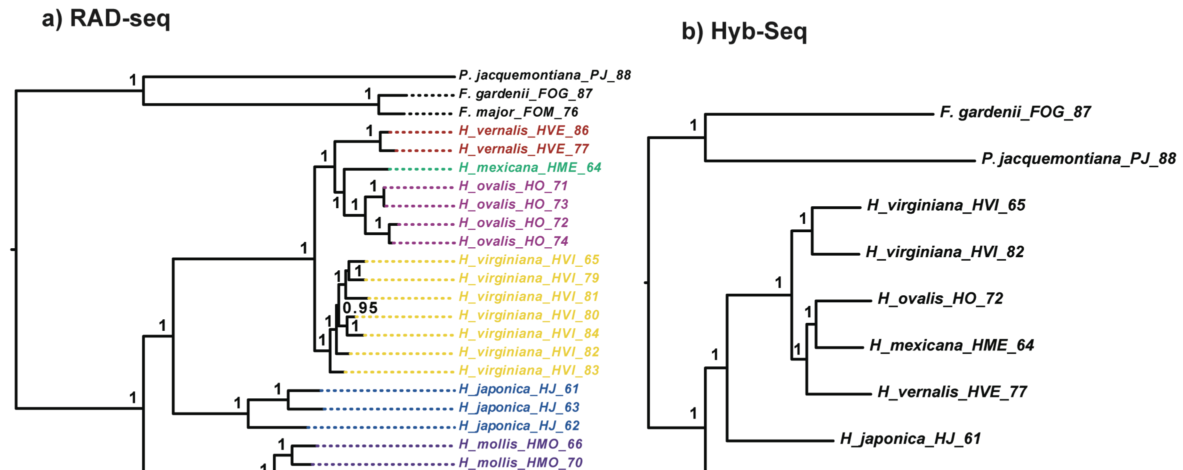


Fig. S19 Concatenated gene tree of *Hamamelis* from analysis of a) RAD-seq M50 and b) primary Hyb-Seq-PPD data using MrBayes. *Parrotiopsis jacquemontiana* and *Fothergilla major* were used as outgroups. Numbers on branches are values of posterior probability.

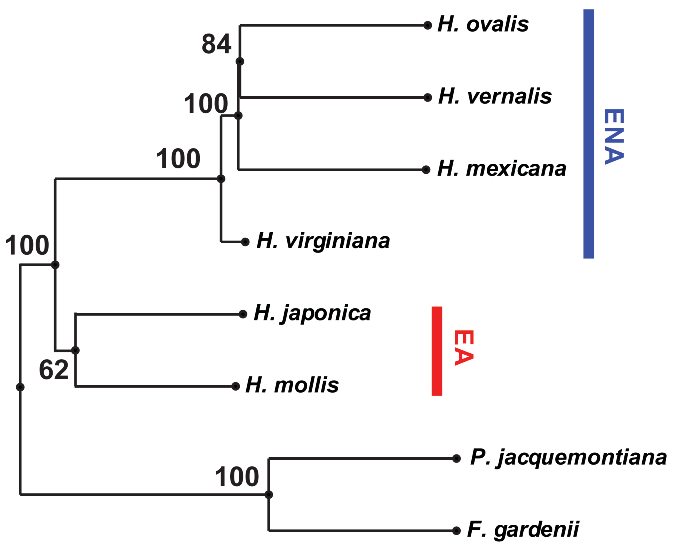


Fig. S20 The species tree of *Hamamelis* from analysis of RAD-seq M50 data using SVDQuartets. *Parrotiopsis jacquemontiana* and *Fothergilla major* were used as outgroups. The result shows that EA clade is a monophyly with relatively low support value 62.

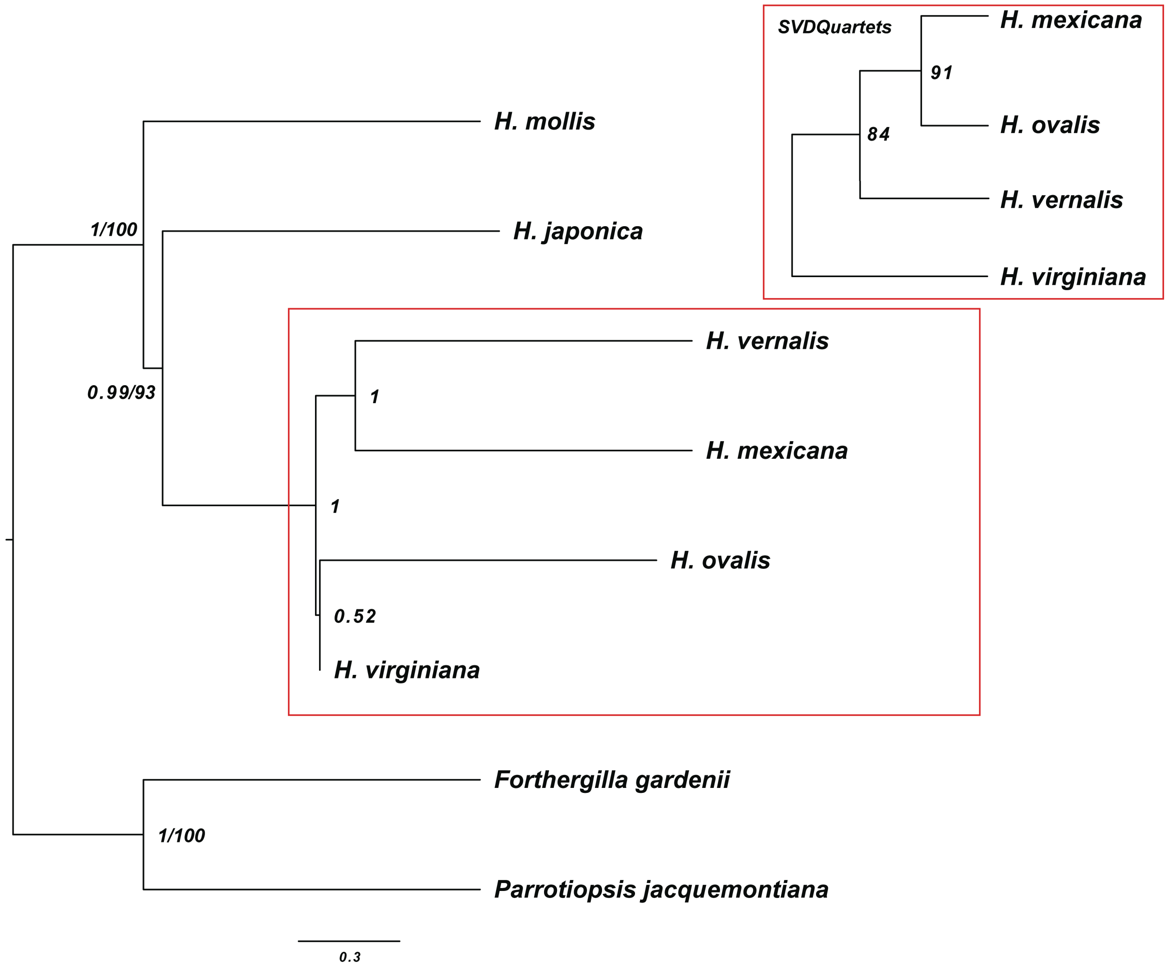


Fig. S21 Species trees of *Hamamelis* from combined RAD-Hyb-Seq data derived from analyses using ASTRAL-III and SVDQuartets. Two species trees were identical except the ENA clade. The numbers on the branches are support values from the ASTRAL-III and SVDQuartets, respectively. The SVDQuartets result of ENA clade was shown in the insert. *Parrotiopsis jacquemontiana* and *Fothergilla gardenii* were used as outgroups.

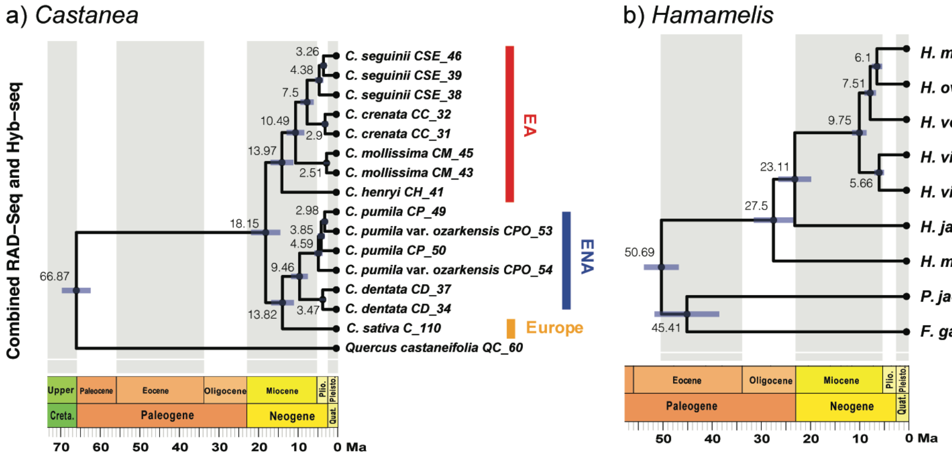


Fig. S22 Divergence times resulting from well-trimmed "BWA100 degenerated" supercontig of orthologous genes from combined RAD-Hyb-Seq data of a) *Castanea* and b) *Hamamelis*. All results showed a similar divergence time results with that from degenerated superontig result from the primary Hyb-Seq data in Fig. 5.

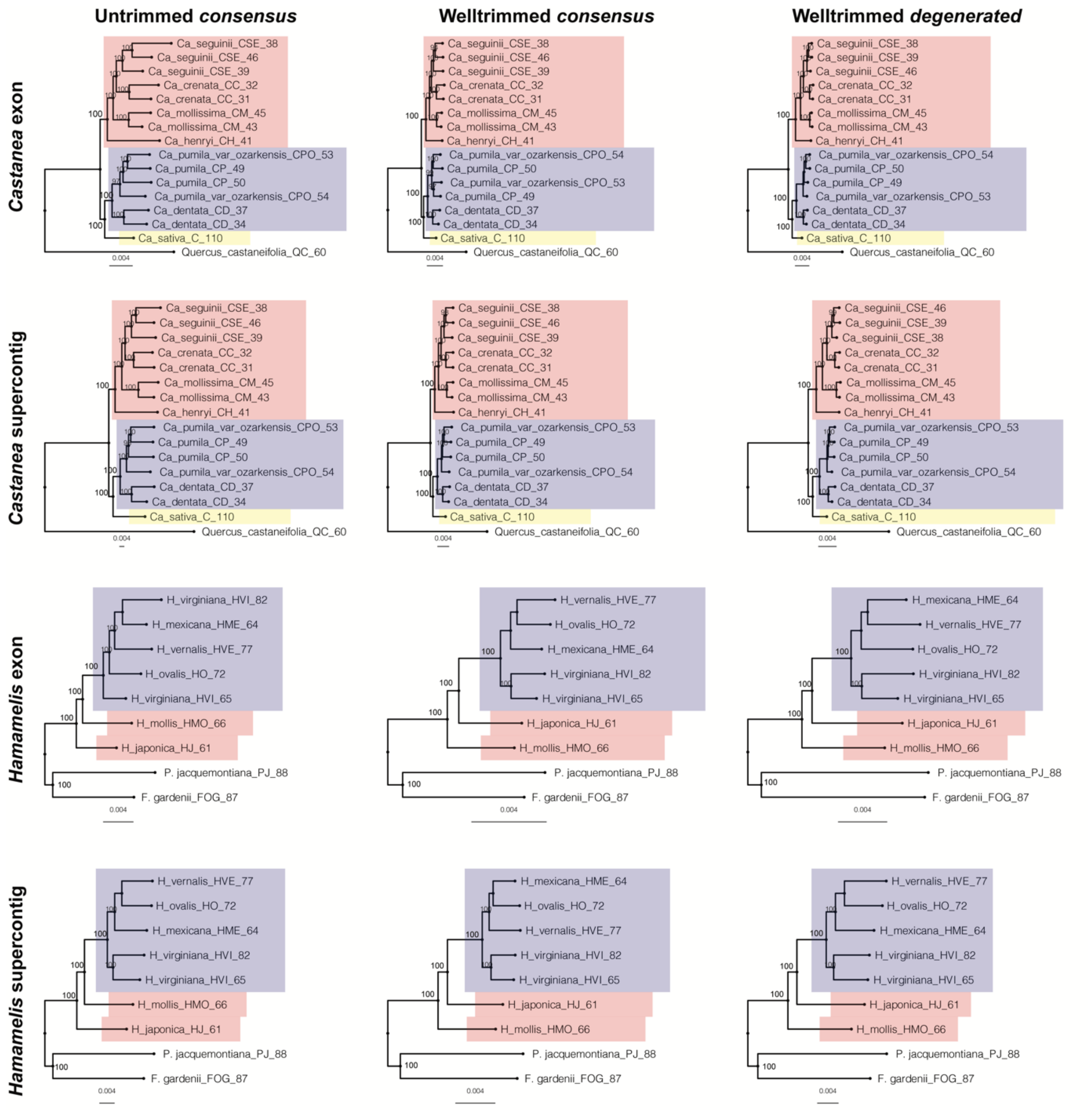


Fig. S23 Comparisons of phylogenies derived from "*consensus*" and "*degenerated*" matrices of the Hyb-Seq data. The first two rows show the results from *Castanea* exon and supercontig regions and the last two rows show the results from *Hamamelis* exon and supercontig regions. The columns show the results from untrimmed "*consensus*", well-trimmed "*consensus*", and well-trimmed "*degenerated*", respectively. The results indicated the tree topologies are identical in both "*consensus*" and "*degenerated*", but "*consensus*" matrices have longer branch lengths than "*degenerated*" matrices. The ENA clades in both *Castanea* and *Hamamelis* are highlighted by blue color, EA clades are highlighted by red, and European clades are highlighted by yellow.

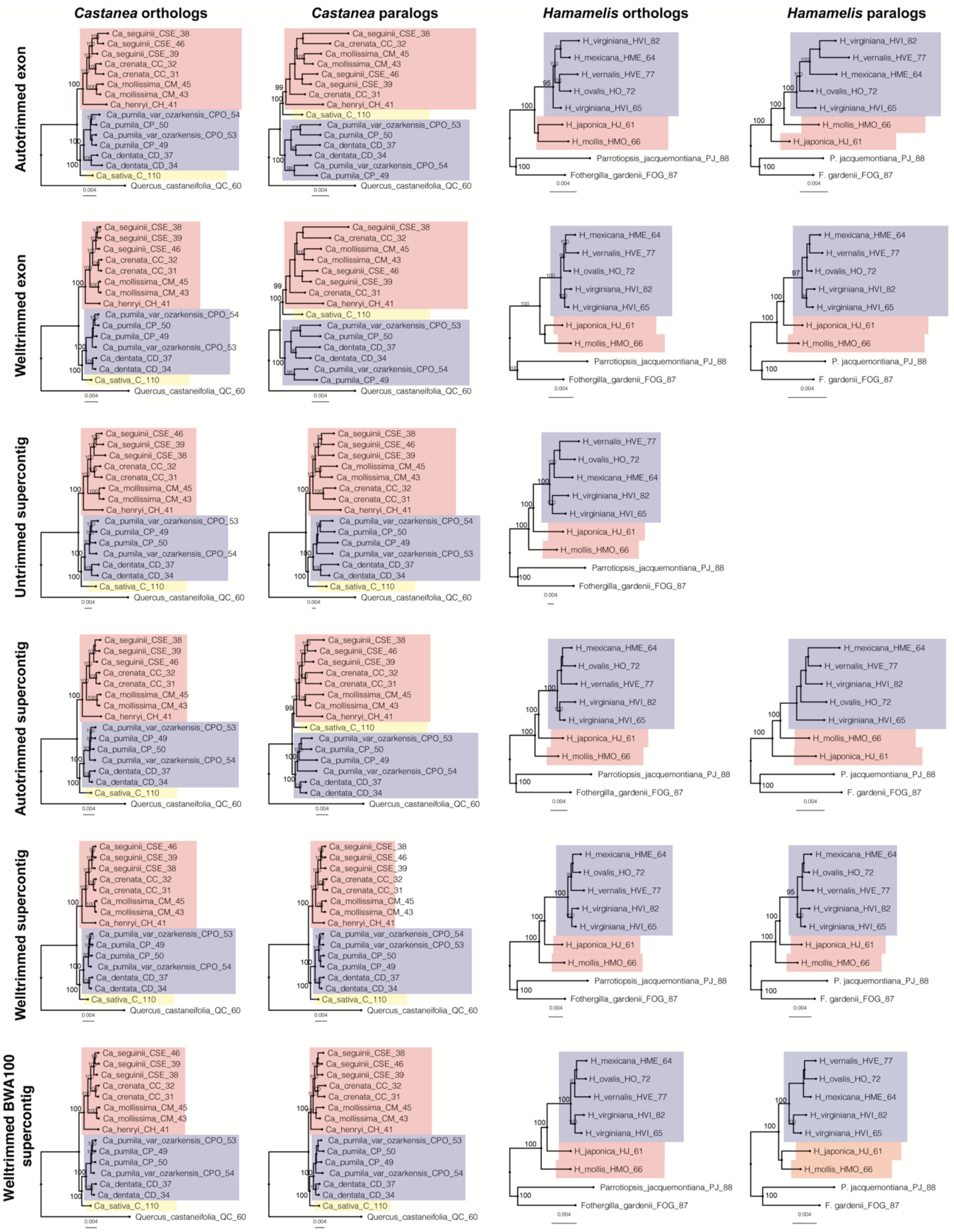

Fig. S24 Comparisons of phylogenies derived from *orthologs* vs. paralogs, exon vs. supercontig, matrices using different trimmed matrices and different mapping methods of the Hyb-Seq data. In orthologs and paralogs comparison, the result shows the paralogs in exon regions have different tree topologies with orthologs in both *Castanea* and *Hamamelis*. “well-trimmed” matrices have shorter branch length than “untrimmed” matrices (also shown in Table S5). “auto-trimmed” matrices have shortest branch lengths and different tree topologies in paralogs. The ENA clades in both *Castanea* and *Hamamelis* are highlighted by blue color, EA clades are highlighted by red, and European clades are highlighted by yellow.

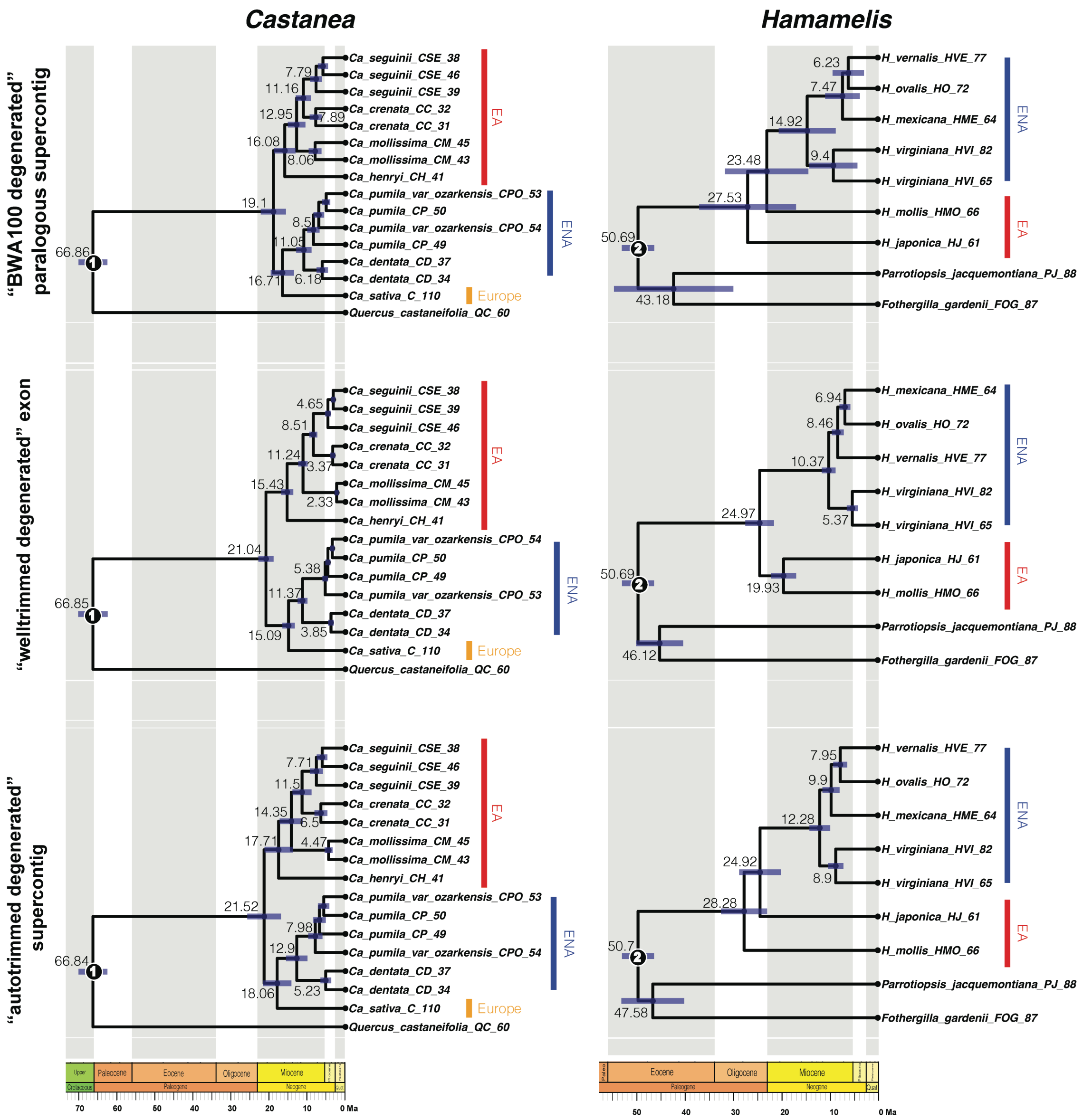


Fig. S25 Divergence times resulting from "BWA100 degenerated" paralogous supercontig, "well-trimmed degenerated" exon, and "auto-trimmed degenerated" supercontig matrices in Hyb-Seq. All results showed a similar divergence time results with that from degenerated superontig result from Hyb-Seq in Fig. 5. But the "BWA100 degenerated" paralogous supercontig results show much older divergence times at shallow nodes (species level) (also shown in Table S5).

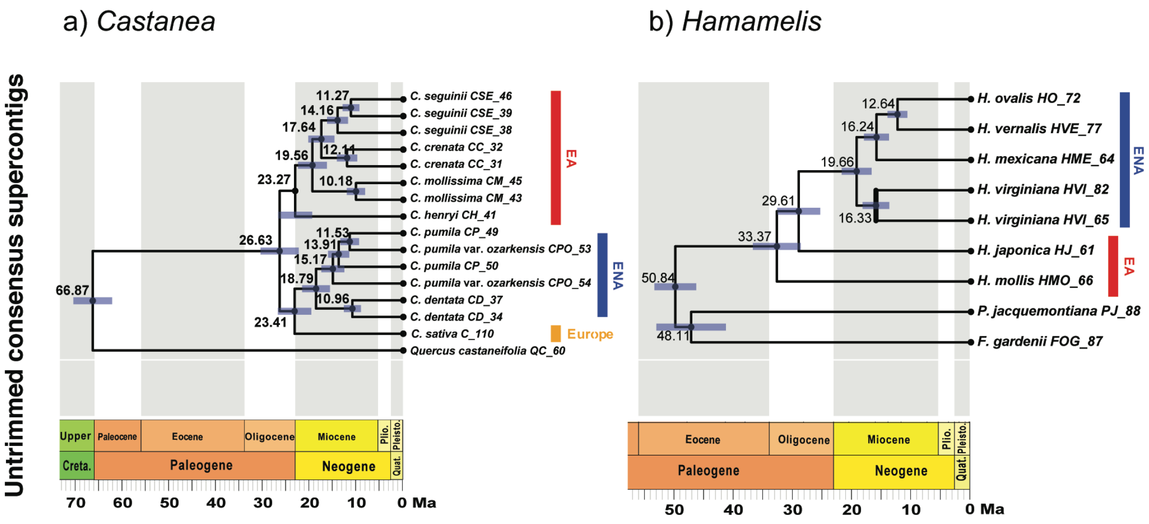


Fig. S26 Divergence times resulting from untrimmed Hyb-Seq "*consensus*" supercontig of orthologous genes of *Castanea* and *Hamamelis*. All results showed much older divergence time results with that from well-trimmed "*degenerated*" and "*consensus*" superontig results from Hyb-Seq data in Fig. 5.

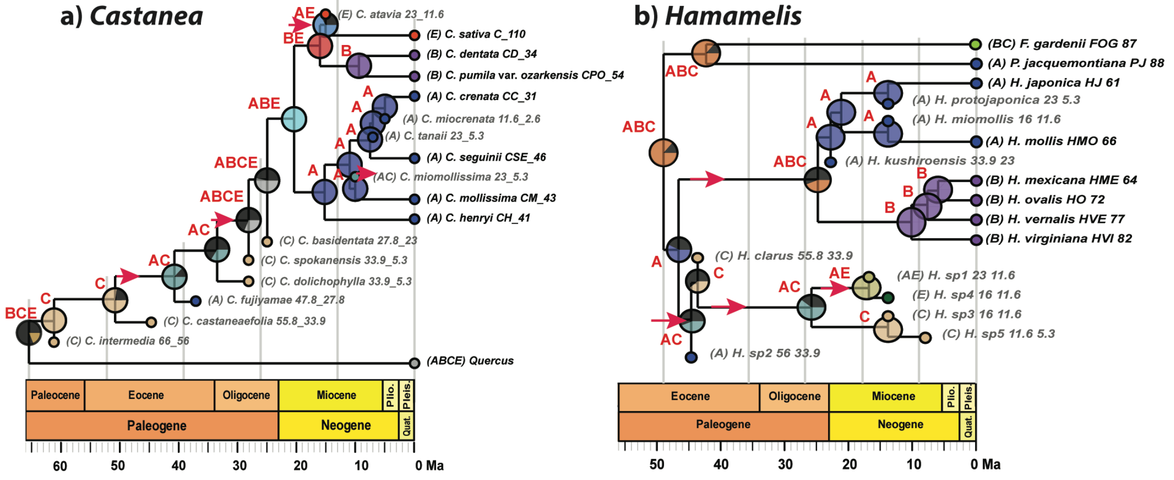


Fig. S27 Results of biogeographic analyses of *Castanea* and *Hamamelis* using the DEC model based on Hyb-Seq data. a). Most likely ancestral ranges reconstructed for *Castanea* based on the total-evidence phylogeny with *Quercus castaneifolia* as outgroup. b). Most likely ancestral ranges reconstructed for *Hamamelis* based on the total-evidence phylogeny with *Parrotiopsis jacquemontiana* and *Fothergilla gardenii* as outgroups. A: eastern Asia; B: eastern North America and Mexico; C: western North America; E: Europe. The letters at each node indicate the most-likely ancestral distributions inferred from the analyses, whose probabilities are shown by non-black colors in the pie charts. The black color in the pie chart indicates the total proportion (likelihood) of all other alternative ancestral ranges in the results of DEC analyses. Intercontinental dispersal events inferred were indicated by red arrows. Fossil taxa are labeled in gray color.
