## Supplementary Information for "A New Paralog Removal Pipeline Resolves Conflict between RAD-seq and Enrichment"

***SUPPLEMENTARY INTRODUCTION***

*Pipelines Developed for Enrichment Data*

In pipelines developed for analyses of target enrichment data, usually an arbitrary cut off of sequence identity value between the contigs of a putative Hyb-Seq locus and the reference target gene is used to determine if the locus contains paralogous sequences in an individual. Currently, HybPiper (Johnson et al. 2016), PHYLUCE (Faircloth 2016), and SECAPR (Andermann et al. 2018) are three popular pipelines for target enrichment data analyses. HybPiper uses BWA (Li and Durbin 2009) or BLASTx (Altschul et al. 1990) to classify the raw reads into individual gene locus, followed by SPAdes (Bankevich et al. 2012) to assemble the reads in a given individual into contigs. In HybPiper, if multiple contigs with a >10x coverage depth in an individual mapped to the same target gene with >85% sequence identity, this target gene is marked for presence of paralogs in the individual (Fig. 1b), which can be eliminated or need special care to determine its orthology to sequences of the same locus of other individuals in subsequent analyses. Both PhylUCE and SECAPR have similar steps in the data treatments, but these treatments use tool kits different from those in HybPiper. These pipelines apply SPAdes (Bankevich et al. 2012) and Abyss (Simpson et al. 2009), respectively, to assemble the raw reads first, and retain all contigs that can match to the target genes. But the method for paralog detection is only slightly different from that in HybPiper. Both PHYLUCE and SECAPR identify paralogous genes based on the observation of multiple contigs in an individual mapping to the same target genes; PHYLUCE also identifies paralogous genes by evidence of a contig matching two or more different target genes (Fig. 1c). Herrando-Moraira et al. (2018) showed that many more paralogous loci were found in the Hyb-Seq data of the tribe Cardueae (Compositae) using the PHYLUCE pipeline than using the HybPiper pipeline. In summary, all of the three aforementioned pipelines consider only sequence similarity to the target gene sequences in detecting paralogous genes.

In addition, most pipelines for enrichment data do not by default consider the sequence variants of contigs/loci generated through the pipelines and most assembly method SPAdes (Bankevich et al. 2012) and Abyss (Simpson et al. 2009) can only construct a single consensus sequence based on the most frequent base (reads) result. This approach loses all information from heterozygous sites for identification of potential paralogs, and may result in data containing phylogenetic noises from paralogous genes that can mislead the inference of species relationship. Although SECAPR and scripts from Kates et al. (2018) can perform allele phasing, all of the presently widely used pipelines for handling target enrichment data do not make use of the information from heterozygous sites to detect paralogous sequences.

***SUPPLEMENTARY METHODS***

***Pipeline Description***

The putative paralogs pipeline includes two major steps: first, a “*degenerated*” alignment matrix is generated from mapping reads back to contigs, which applies the IUPAC ambiguity codes to capture the heterozygous sites in generating the exon and supercontig matrices of the three sets of aforementioned genes; and second, the matrix is trimmed of highly heterozygous sites, misaligned regions, and particularly gappy columns.

The first part of PPD is to obtain the “*degenerated*” sequences for the exon region and supercontig of each locus. We used a bash script following Kates et al. (2018) (available on Github: <https://github.com/Bean061/putative_paralogs>). This involved using the “*consensus*” sequences from HybPiper as the references, and mapping the raw reads back to the references in BWA with the default minimum mapped seed length (-k) of 100 bp. After mapping, the mapped duplicate reads were discarded using picard (<https://broadinstitute.github.io/picard/>). The program GATK (McKenna et al. 2010; DePristo et al. 2011) was then used to identify the variable sites using the HaplotypeCaller, with “-ploidy 2” parameter for diploid species, and SelectVariants functions. Finally, we used the FastaAlternateReferenceMaker function in GATK to produce the “*degenerated*” (IUPAC) sequences for each gene.

The second part is used to trim alignment and detect the paralogous genes. It includes 8 steps: s1) Resort gene files: Use all “*degenerated*” sequence files from every individual as the input, and then sort the degenerated sequences orthologous to the 353 reference genes in each sample into individual locus files according to gene names. s2) Sequence filtering: Filter the sequences with more than 5% (default) heterozygosity according to the percentage information of heterozygous sites in every sequence. This setting can be changed by users with “-he” parameter. s3-s5) MSA generating: To obtain the better alignment result, add the reference sequence of each locus for alignment using MAFFT (--adjustdirection –maxiterate 1000 --globalpair) (Katoh and Standley 2013). The reference sequences were removed before trimming of the aligned sequences. s6) MSA trimming: Remove the gappy sites (i.e., sites missing in 50% or more individuals) using TrimAl (“-gt 0.51”). s7) MSA further trimming: Detect and trim the hypervariable sites using a sliding window method as shown in the red box. The polymorphic sites meeting the requirement in each window were marked and then removed from all individuals by TrimAl. The maximum sites number in a sliding window can be modified by “-m” parameter and sliding window length can be modified by “-w” parameter in PPD. The default values for “m” and “w” are 5 and 20, respectively, which represent if there are more than 4 polymorphic sites (not counting sites with heterozygous bases) in a 20 bp sliding window, all the polymorphic sites will be marked and removed by TrimAl. These criteria should be adjusted according to your observation of the non-trimmed taxa MSA. s8) Paralog identification: Consider a locus as paralogs if heterozygous site(s) shared by 50% (default) or more individuals at a locus. It can be changed according to your taxa using the “-hs” parameter. A hypothetical MSA of a locus/gene (on the left side) shows sequence with high heterozygous sites (top one), a polymorphic site that is heterozygous in >50% samples/individuals of a diploid organism (labeled as polymorphic site 2 in figure 2), and a sequence containing a region with apparent alignment ambiguity due to error in contig assembly (shown as hypervariable sites compared to the rest in figure 2). Identical heterozygous site(s) shared by over 50% individuals (Polymorphic site 2 in figure 2) in the MSA is used as the indication of presence of paralogs in the locus and is the criterion for calling putative paralogs in the PPD.

***Different Matrices for Hyb-Seq Data***

***The “consensus” vs. “degenerated” matrices***.—To directly evaluate the impact of character coding, we used the pipeline of HybPiper (Johnson et al. 2016) to generate “*consensus*” sequences of supercontig and exons for each locus, followed by well-trimmed procedure in PPD, and used the whole steps of PPD to generate the “*degenerated*” matrices (see Table 2).

***Matrices with different trimming methods***.—To demonstrate the importance of these customized trimming in PPD (“well-trimmed” method) steps that remove missing data at the end of locus, hypervariable sites in regions containing error sequences in some samples, and degenerated sequences indicative of paralogy, we generated data without any trimming (“untrimmed” method) or automatically trimmed once by TrimAl with the “automated1” parameter (“auto-trimmed” method) for comparison (see Table 2). For well-trimmed “*consensus*” matrices, we used the sequences directly from HybPiper and used the PPD to trim the them.

***Orthologs vs. paralogs***.—To compare the phylogenetic results from orthologs and paralogs, we generated Orthologs and paralogs based on PPD pipeline. We also compared the orthologous and paralogous matrices from different trimming method and different DNA regions (supercontig vs. exon), see Table 2.

***Supercontig vs. exon***.—To compare the phylogenetic results from supercontig and exon, we used the supercontig and exon matrix identified by HybPiper, followed by PPD pipeline and different trimming methods (Table 2).

***BWA parameters***.—To assess how mapping accuracy in BWA step may affect the subsequent analyses, we compared the default setting of minimum mapped seed length (19 bp) to our setting in PPD (100 bp). Our length is close to the length of the raw reads (~150 bp) after quality control of removing barcode and low quality sequences. The “*degenerated*” sequences of the supercontigs with higher mapping accuracy (“BWA100” matrices in Table 2) were generated for orthologous genes, paralogous genes, and all-genes, respectively, for comparisons with those generated using BWA k=19 bp (“BWA19” matrices in Table 2).

***Chloroplast genes from off-targeted enrichment data***.—We used two downloaded chloroplast genomes, *Castanea mollissima* (NC_014674, Jansen et al. 2011) and *Hamamelis mollis* (NC_037881, Dong et al. 2018) as the references to obtain the off-targeted chloroplast data. We used BWA (mem -k 100) and samtools (view -f 3) to get mapped reads to these two references into a bam format, followed by using Geneious v.2020.1.1 to generate a consensus sequence covered by more than 3x depth reads. Because all sequences were fragmented, we used the “--adjustdirection --addfragments” function in MAFFT to obtain the alignment for chloroplast data with the chloroplast genome as the reference. Finally, the alignment was trimmed by TrimAl with -gt parameter as 0.7 (filtering sites with gaps shared in over 30% individuals).

***Combining genes from RAD-seq and Hyb-Seq data***.—We used the *consensus* sequence of orthologous “*degenerated* BWA100” supercontig matrix as the reference to identify loci shared between RAD-seq and Hyb-seq data using ipyrad with the “denovo-reference” function. This allows us to remove sequences redundant to those in Hyb-Seq from the RAD-seq data. This filtering process used the same settings of parameters as described above for the M50 matrix. The filtered RAD-seq data were then combined with the primary Hyb-Seq-PPD data for the accessions with both data available.

***Fossils of Castanea and Hamamelis Used for Fossilized Birth Death Analysis***

In the divergence time analyses, the stem age of each genus was constrained by fossils. The earliest fossil of *Castanea* dates to the Paleocene/Eocene (55-66 Ma; Nixon and Crepet 1989) (Table 3) and was used as the minimum age of the stem node of the genus, while the earliest fossil of Fagaceae dates to the Campanian and Santonian in the Late Cretaceous (72-86 Ma; Herendeen et al. 1995; Sims et al. 1998) (Table 3) and was used as the maximum age of the stem node of the genus. For *Hamamelis* and its outgroups, *Fothergilla malloryi* Radtke, Pigg et Wehr sp. Nov. is the earliest fossil of *Fothergilla* and dates back to the early Eocene (49-50 Ma) (Table 3). Fossils of *Hamamelis* are younger than this fossil (Table 3). We therefore used the *Fothergilla* fossil to set minimum age of the crown node containing both genera. In Hamamelidaceae, the inflorescence and seed fossils of *Hamawilsonia Boglei* Bendict, Pigg & DeVore gen. et sp. nov. is the earliest and was found in the late Paleocene (56-59 Ma) of the United States (Table 3). The inflorescence of the fossil species shows similarity to *Hamamelis* while its seed fossils resemble *Sinowilsonia* (Benedict et al. 2008). *Sinowilsonia* is a genus in a separate clade sister to the clade containing *Hamamelis* and *Fothergilla* in the phylogeny of Hamamelidaceae (Li and Bogle 2001; Benedict et al. 2008). *Hamawilsonia*, therefore, likely represents an extinct common ancestor of *Hamamelis* and *Sinowilsonia*, given its morphology, which is older than the common ancestor of *Hamamelis* and *Fothergilla*. Therefore, we used the age of this fossil as the maximum bound of the crown node containing *Hamamelis* and *Fothergilla* (or the stem node of *Hamamelis*). In other words, the stem age of *Hamamelis* was set as 50-56 Ma in the divergence time analysis of *Hamamelis*. The lognormal distribution was used as the fossil calibration prior for both genera.

***Fossils of Castanea and Hamamelis used for Fossilized Birth Death analysis***

Fossil species *Castanea miomollissima* Hu and Chaney, similar to the modern species *C. mollisima*, was reported from the Miocene beds of Montana, western North America, and Japan in the Miocene (Becker 1969; Ishida 1970). Two other fossils, *Castanea tanaii* Huzioka (Huzioka 1972) and *Castanea miocrenata* Tanai *et* Onoe (Ozaki 1991), with affinities to *C. crentata*, were reported from the Miocene Korea and the late Miocene to Pliocene beds of Japan, respectively. Furthermore, *Castanea atavia* Unger (Van der Burgh 1983; Mai 2001), a fossil species allied to *C. sativa*, was found in the upper and mid Miocene beds of Germany. The remaining fossils represent taxa on the stems of the modern *Castanea* clade as no morphological characters are available to assign them any modern lineages. These fossils include the Oligocene *Castanea basidentata* R.N. Lakhanpal (Lakhanpal 1958) and *Castanea dolichophylla* Cockerell (= *Castanea orientalis* Chaney) (Hoffman 1932; MacGinitie 1953; Becker 1969), the Oligocene and Miocene *Castanea spokanensis* (Knowlton) Chaney and Axelrod (Becker 1972, 1973), the Eocene *Castanea castaneaefolia* (Unger) Knowlton (Hollick 1936), and the Paleocene *Castanea intermedia* Lesquereux (Brown 1962), all from western North America, and the mid-Eocene and Oligocene *Castanea fujiyamae* Tanai from Jilin, China and Japan (Tanai 1970, 1995; Manchester et al. 2005). In *Hamamelis*, three reliable fossils were used in the study: the Eocene *Hamamelis clarus* Hollick (Hollick1936) from Alaska, US, the Oligocene *Hamamelis kushiroensis* Tanai (Tanai 1970) from Japan and the Miocene *Hamamelis protojaponica* Tanai *et* N. suzuki from Korea (Huzioka 1972). The latter two fossils resemble the modern species *H. japonica*, according to the authors (Tanai 1970; Huzioka 1972).

***SUPPLEMENTARY RESULTS***

***Results on Branch Lengths***

For the supercontig data, the “automated1” argument in TrimAl filtered many more sites than the “well-trimmed” method applied in our PPD pipeline (e.g., average length per locus in data containing all genes: ~1082 bp vs. ~2426 bp in *Castanea* and ~759 bp vs. 2884 bp in *Hamamelis*; Table 3). For exon data, the matrices from “auto-trimmed” and “well-trimmed” were similar in sequence length per locus (~600 bp in both *Castanea* and *Hamamelis*; Table 3). In “untrimmed” matrices, the branch lengths of trees from the paralogs were much longer than the branch lengths of the tree from orthologs, but both were longer than the branches in the tree from the RAD-seq data (Supplementary Fig. S24, available on Dryad). Additionally, the branch lengths of the trees from “untrimmed” matrices were much longer than the branch lengths of trees from “well-trimmed” matrices in both “*consensus*” and “*degenerated*” sequences (Supplementary Figs. S23 and S24, available on Dryad). In the comparison between k values in BWA mapping, the results showed no differences in tree topologies from data derived from the default setting of k=19 pb (all “degenerated BWA19” orthologs matrices in Table 3) or from the more stringent setting of k=100 bp (the “*degenerated* BWA100” in Table 3; Fig. S24). The numbers of putative paralogs identified were, however, much lower with k = 100 bp (48 in *Castanea* and 27 in *Hamamelis*), compared to the numbers identified with k = 19 bp (116 paralogs in *Castanea* and 106 paralogs in *Hamamelis*) (see Table 3 under “*degenerated* BWA19” vs. “*degenerated* BWA100”, with supercontig and well-trimmed). In all analyses with our PPD (under “*degenerated*” in Table 3) pipelines, we found more paralogs than the analyses with HybPiper, which detected only 11 paralogs in *Castanea* and 2 paralogs in *Hamamelis*.

***SUPPLEMENTARY REFERENCE***

Altschul S.F., Gish W., Miller W., Myers E.W., Lipman D.J. 1990. Basic local alignment search tool. J. Mol. Biol. 215:403–410.

Andermann T., Cano Á., Zizka A., Bacon C., Antonelli A. 2018. SECAPR—a bioinformatics pipeline for the rapid and user-friendly processing of targeted enriched Illumina sequences, from raw reads to alignments. PeerJ. 6:e5175.

Bankevich A., Nurk S., Antipov D., Gurevich A.A., Dvorkin M., Kulikov A.S., Lesin V.M., Nikolenko S.I., Pham S., Prjibelski A.D., Pyshkin A.V., Sirotkin A.V., Vyahhi N., Tesler G., Alekseyev M.A., Pevzner P.A. 2012. SPAdes: A New Genome Assembly Algorithm and Its Applications to Single-Cell Sequencing. J. Comput. Biol. 19:455–477.

Becker H.F. 1969. Fossil plants of the Tertiary Beaverhead Basins in southwestern Montana. Palaeontographica, Abt. B. 127:1–142.

Becker H.F. 1972. The Metzel Ranch flora of the upper Ruby River basin, southwestern Montana. Palaeontographica Abt. B. 141:1–61.

Becker H.F. 1973. The York Ranch flora of the upper Ruby River basin, southwestern Montana. Palaeontographica Abt. B. 143:18–93.

Benedict J.C., Pigg K.B., DeVore M.L. 2008. *Hamawilsonia boglei* gen. et sp. nov.(Hamamelidaceae) from the late Paleocene Almont flora of central North Dakota. Int. J. Plant Sci. 169:687–700.

Brown R.W. 1962. Paleocene flora of the Rocky Mountains and great Plains. Washington: U.S. Govt. Print. Off.p. 1-119.

DePristo M.A., Banks E., Poplin R., Garimella K.V., Maguire J.R., Hartl C., Philippakis A.A., del Angel G., Rivas M.A., Hanna M., McKenna A., Fennell T.J., Kernytsky A.M., Sivachenko A.Y., Cibulskis K., Gabriel S.B., Altshuler D., Daly M.J. 2011. A framework for variation discovery and genotyping using next-generation DNA sequencing data. Nat. Genet. 43:491–498.

Dong W., Xu C., Wu P., Cheng T., Yu J., Zhou S., Hong D.-Y. 2018. Resolving the systematic positions of enigmatic taxa: Manipulating the chloroplast genome data of Saxifragales. Mol. Phylogenet. Evol. 126:321–330.

Faircloth B.C. 2016. PHYLUCE is a software package for the analysis of conserved genomic loci. Bioinformatics. 32:786–788.

Herendeen P.S., Crane P.R., Drinnan A.N. 1995. Fagaceous flowers, fruits, and cupules from the Campanian (Late Cretaceous) of central Georgia, USA. Int. J. Plant Sci. 156:93–116.

Herrando-Moraira S., Calleja J.A., Carnicero P., Fujikawa K., Galbany-Casals M., Garcia-Jacas N., Im H.-T., Kim S.-C., Liu J.-Q., López-Alvarado J., López-Pujol J., Mandel J.R., Massó S., Mehregan I., Montes-Moreno N., Pyak E., Roquet C., Sáez L., Sennikov A., Susanna A., Vilatersana R. 2018. Exploring data processing strategies in NGS target enrichment to disentangle radiations in the tribe Cardueae (Compositae). Mol. Phylogenet. Evol. 128:69–87.

Hoffman A.D. 1932. The Douglas Canyon flora of east central Washington. J. Geol. 40:735–738.

Hollick C.A. 1936. The tertiary floras of Alaska. Washington: U.S. Govt. Print. Off. p. 1–171.

Huzioka K. 1972. The tertiary floras of Korea. J. Ming. Coll. Akita Univ. 5(1): 1–83.

Ishida S. 1970. The Noroshi flora of Note Peninsula, Central Japan. Memoirs of the Faculty of Science, Kyoto University. Series of Geology and Mineralogy. 37(1):1–112.

Jansen R.K., Saski C., Lee S.-B., Hansen A.K., Daniell H. 2011. Complete Plastid Genome Sequences of Three Rosids (*Castanea*, *Prunus*, *Theobroma*): Evidence for At Least Two Independent Transfers of rpl22 to the Nucleus. Mol. Biol. Evol. 28:835–847.

Johnson M.G., Gardner E.M., Liu Y., Medina R., Goffinet B., Shaw A.J., Zerega N.J.C., Wickett N.J. 2016. HybPiper: Extracting Coding Sequence and Introns for Phylogenetics from High-Throughput Sequencing Reads Using Target Enrichment. Appl. Plant Sci. 4:1600016.

Katoh K., Standley D.M. 2013. MAFFT multiple sequence alignment software version 7: improvements in performance and usability. Mol. Biol. Evol. 30:772–80.

Lakhanpal R.N. 1958. The Rujada Flora of West Central Oregon. University of California Publications in Geological Sciences. 35:1–65.

Li H., Durbin R. 2009. Fast and accurate short read alignment with Burrows-Wheeler transform. Bioinformatics. 25:1754–1760.

Li J., Bogle A.L. 2001. A New Suprageneric Classification System of the Hamamelidoideae Based on Morphology and Sequences of Nuclear and Chloroplast DNA. Harv. Pap. Bot. 5(2):499–515.

MacGinitie H.D. 1953. Fossil plants of the Florissant Beds, Colorado. Washington, D. C.: Carnegie Institution of Washington. p. 1–198.

Mai D.H. 2001. Die mittelmiozänen und obermiozänen Floren aus der Meuroer und Raunoer Folge in der Lausitz. Teil III: Fundstellen und Palaeobiologie. Palaeontogr. Abt. B. 258:1–85.

Manchester S.R., Chen Z., Geng B., Tao J. 2005. Middle Eocene flora of Huadian, Jilin Province, Northeastern China. Acta Palaeobot. 45(1):3–26.

McKenna A., Hanna M., Banks E., Sivachenko A., Cibulskis K., Kernytsky A., Garimella K., Altshuler D., Gabriel S., Daly M., DePristo M.A. 2010. The Genome Analysis Toolkit: A MapReduce framework for analyzing next-generation DNA sequencing data. Genome Res. 20:1297–1303.

Nixon K.C., Crepet W.L. 1989. *Trigonobalanus* (Fagaceae): taxonomic status and phylogenetic relationships. Am. J. Bot. 76:828–841.

Ozaki K. 1991. Late Miocene and Pliocene floras in central Honshu, Japan. Yokohama, Japan: Kanagawa Prefectural Museum. p. 1–244.

Simpson J.T., Wong K., Jackman S.D., Schein J.E., Jones S.J.M., Birol I. 2009. ABySS: A parallel assembler for short read sequence data. Genome Res. 19:1117–1123.

Sims H.J., Herendeen P.S., Crane P.R. 1998. New genus of fossil Fagaceae from the Santonian (Late Cretaceous) of central Georgia, USA. Int. J. Plant Sci. 159:391–404.

Tanai T. 1970. The Oligocene floras from the Kushiro coal field, Hokkaido, Japan. J. Fac. Sci., Hokkaido Univ., Ser. 4, Geology and mineralogy. 14:383–514.

Tanai T. 1970. The Oligocene floras from the Kushiro coal field, Hokkaido, Japan. Journal of the Faculty of Science, Hokkaido University. Series 4, Geology and mineralogy. 14:383–514.

Tanai T. 1995. Fagacean Leaves from the Paleogene of Hokkaido, Japan. Bull. Natl. Mus. Nat. Sci. Ser. C. 21:71–102.

Van der Burgh J. 1983. Allochthonous seed and fruit floras from the Pliocene of the Lower Rhine Basin. Rev. Palaeobot. Palynol. 40:33–90.

***SUPPLEMENTARY* *TABLES*:**

**TableS1. Morphological Characters and character states included in *Castanea* and *Hamamelis* and their fossil records.**

| **Species** | **Leaf shape** | **Leaf length (cm)** | **Leaf width (cm)** | **Position of Leaf Maximum width (from base):** | **Number of secondary vein pairs** | **Degree between secondary vein and main vein** | **Distance between two secondary veins** |
| --- | --- | --- | --- | --- | --- | --- | --- |
| ***C. mollissima*** | Elliptic-oblong to oblong-lanceolate (0) | 18 (0) | 6.3 (0) | 0.5 (0) | 15 (1) | 65 (0) | 1 cm (0) |
| ***C. miomollissima*** | Lanceolate (0) | 8--12 (1) | 2--4 (2) | NA | NA | NA | NA |
| ***C. dentata*** | Obovate to oblanceolate (1) | 17.9 (0) | 6 (0) | 0.5 (0) | 21 (0) | 50 (1) | 0.7 (0) |
| ***C. sativa*** | Oblong-lanceolate (0) | 16.1 (0) | 4.3 (1) | 0.5 (0) | 15 (1) | 45 (1) | 1 cm (0) |
| ***C. henryi*** | Oblong-ovate, oblong-lanceolate, or lanceolate (0) | 13.5 (1) | 5.5 (0) | 0.5 (0) | 13 (1) | 40+ (1) | 1 cm (0) |
| ***C. seguinii*** | Oblong-obovate to elliptic-oblong (0) | 11.4 (1) | 3.5 (2) | 0.66 (1) | 18 (0) | 60+ (0) | 0.8 (0) |
| ***C. crenata*** | Oblong-lanceolate (0) | 12.4 (1) | 4--5 (1) | 0.5 (0) | 15 (1) | 55 (1) | 1 cm (0) |
| ***C. pumila*** | Obovate or oblanceolate (1) | 9.7 (2) | 3--4 (2) | 0.5 (0) | 14 (1) | 55 (1) | 0.4 (1) |
| ***C. fujiyamae*** | NA | NA | NA | 0.5 (0) | 13 (1) | NA | NA |
| ***C. basidentata*** | Lanceolate (0) | 9--20 [12] (1) | 2--4 [3.5] (2) | 0.5 (0) | 18-20 (0) | 70 (0) | NA |
| ***C. dolichophylla (C. orientalis)*** | Lanceolate (0) | 12--15 (1) | 2--2.5 (2) | 0.33 (2) | NA | NA | 1 cm (0) |
| ***C. miocrenata*** | Oblong (0) | NA | wider than *C. crenata* (0) | 0.5 (0) | NA | NA | NA |
| ***C. spokanensis*** | Oval (0) | NA | NA | 0.5 (0) | NA | NA | NA |
| ***C. tanaii*** | Linear lanceolate (0) | NA | 1.5--2.5 (2) | 0.5 (0) | NA | 50 (1) | 0.5 cm (1) |
| ***C. intermedia*** | NA | NA | NA | NA | NA | NA | NA |
| ***C. castaneaefolia*** | NA | NA | NA | NA | NA | NA | NA |
| ***C. atavia*** | NA | NA | NA | NA | NA | NA | NA |

Notes: NA means no available data.

Leaf shape: elliptic-oblong to oblong-lanceolate and ovate is 0, obovate to oblanceolate is 1.

Leaf length: average length longer than 15 cm (0); between 10 to 15 cm (1); shorter than 10 cm (2)

Leaf width: average width longer than 5 cm (0); between 4 to 5 cm (1); shorter than 4 cm (2)

Position of Leaf Maximum width (from base): equal to 0.5 (0), less than 0.5 (1), greater than 0.5 (2)

Number of secondary vein pairs: more than 15 pairs is 0; less or equal to 15 is 1

Degree between secondary vein and main vein: greater than 60° (0); less than 60° (1)

Distance between two secondary veins: greater than 0.5 cm is 0; less than 0.5 cm is 1.

| **Species** | **Leaf shape** | **Leaf length (cm)** | **Leaf width (cm)** | **Leaf base shape** | **Degree between secondary vein and main vein** | **Apex shape** |
| --- | --- | --- | --- | --- | --- | --- |
| ***H. clarus*** | ovate-lanceolate (0) | 8 (1) | 5.25 (1) | Base rounded (0) | <45 (0) | Cuneate, acute (0) |
| ***H. kushiroensis*** | NA | NA | NA | NA | NA | NA |
| ***H. protojaponica*** | NA | NA | NA | NA | NA | NA |
| ***H. mollis*** | Ovate (0) | 10 (0) | 8.1 (0) | Base rounded (0) | <30 (2) | Subcute to cuspidate (1) |
| ***H. japonica*** | Rhombic (1) | 7.7 (1) | 6.3 (1) | Cuneate or round (1) | 40 (0) | Acute (0) |
| ***H. virginiana*** | Ovate-Rhombic (0,1) | 10.6 (0) | 7.5 (0) | Cuneate or round (1) | 33 (1) | Subacute to cuspidate (1) |
| ***H. ovalis*** | Ovate (0) | 18 (0) | 12 (0) | Cuneate or round (1) | 37 (1) | Subacute to cuspidate (1) |
| ***H. vernalis*** | Rhombic to oblanceolate (2) | 8.9 (1) | 5.9 (1) | Strongly cuneate (2) | <30 (2) | Subacute to round (2) |
| ***H. mexicana*** | Ovate (0) | 6.5 (2) | 4.5 (2) | Cuneate or round (1) | 40 (0) | Subacute to cuspidate (1) |

Notes: NA means no available data

Leaf shape: ovate is 0, Rhombic is 1, Rhombic to oblanceolate is 2.

Leaf length: average length is over 10 cm (0); between 7-10 cm (1); less than 7 cm (2)

Leaf width: average width is over 7 cm (0); between 5-7 cm (1); less than 5 cm (2)

Leaf base shape: Cordate or round (0), Cuneate or round (1), Strongly cuneate (2)

Degree between secondary vein and main vein: greater than 40° is 0; between 30°-40° is 1; less than 30° is 2.

Apex shape: cuneate, acute is 0; subacute to cuspidate is 1; subacute to round is 2.

**Table S2 Dispersal rate matrix between geographic areas for DEC model.**

| **periods** | **Regions** | ***Castanea*** | | | | **Regions** | ***Hamamelis*** | | | |
| --- | --- | --- | --- | --- | --- | --- | --- | --- | --- | --- |
|  |  | **A** | **B** | **C** | **E** |  | **A** | **B** | **C** | **E** |
| 0--4.7 | A | 1 | 1 | 0.001 | 1 | **A** | 1 | 0.001 | 0.001 | 1 |
|  | B | 1 | 1 | 0.001 | 0.001 | **B** | 0.001 | 1 | 0.001 | 0.001 |
|  | C | 0.001 | 0.001 | 1 | 1 | **C** | 0.001 | 0.001 | 1 | 0.001 |
|  | E | 1 | 0.001 | 1 | 1 | **E** | 1 | 0.001 | 0.001 | 1 |
| 4.7--10 | A | 1 | 1 | 1 | 1 | **A** | 1 | 0.001 | 0.001 | 1 |
|  | B | 1 | 1 | 0.001 | 0.001 | **B** | 0.001 | 1 | 0.001 | 0.001 |
|  | C | 1 | 0.001 | 1 | 1 | **C** | 0.001 | 0.001 | 1 | 0.001 |
|  | E | 1 | 0.001 | 1 | 1 | **E** | 1 | 0.001 | 0.001 | 1 |
| 10--15 | A | 1 | 1 | 1 | 1 | **A** | 1 | 0.001 | 1 | 1 |
|  | B | 1 | 1 | 0.001 | 0.001 | **B** | 0.001 | 1 | 0.001 | 0.001 |
|  | C | 1 | 0.001 | 1 | 1 | **C** | 1 | 0.001 | 1 | 0.001 |
|  | E | 1 | 0.001 | 1 | 1 | **E** | 1 | 0.001 | 0.001 | 1 |
| 15--20 | A | 1 | 1 | 1 | 1 | **A** | 1 | 0.2 | 1 | 1 |
|  | B | 1 | 1 | 0.001 | 0.2 | **B** | 0.2 | 1 | 0.001 | 0.2 |
|  | C | 1 | 0.001 | 1 | 1 | **C** | 1 | 0.001 | 1 | 0.001 |
|  | E | 1 | 0.2 | 1 | 1 | **E** | 1 | 0.2 | 0.001 | 1 |
| 20--30 | A | 1 | 1 | 1 | 1 | **A** | 1 | 0.5 | 1 | 1 |
|  | B | 1 | 1 | 0.001 | 0.5 | **B** | 0.5 | 1 | 0.001 | 0.5 |
|  | C | 1 | 0.001 | 1 | 1 | **C** | 1 | 0.001 | 1 | 0.001 |
|  | E | 1 | 0.5 | 1 | 1 | **E** | 1 | 0.5 | 0.001 | 1 |
| 30--38 | A | 1 | 1 | 1 | 0.001 | **A** | 1 | 0.001 | 1 | 0.001 |
|  | B | 1 | 1 | 0.001 | 0.5 | **B** | 0.001 | 1 | 0.001 | 0.5 |
|  | C | 1 | 0.001 | 1 | 1 | **C** | 1 | 0.001 | 1 | 0.001 |
|  | E | 0.001 | 0.5 | 1 | 1 | **E** | 0.001 | 0.5 | 0.001 | 1 |
| 38--45 | A | 1 | 1 | 1 | 0.001 | **A** | 1 | 0.001 | 1 | 0.001 |
|  | B | 1 | 1 | 0.001 | 0.75 | **B** | 0.001 | 1 | 0.001 | 0.75 |
|  | C | 1 | 0.001 | 1 | 1 | **C** | 1 | 0.001 | 1 | 0.001 |
|  | E | 0.001 | 0.75 | 1 | 1 | **E** | 0.001 | 0.75 | 0.001 | 1 |
| 45--60 | A | 1 | 1 | 1 | 0.001 | **A** | 1 | 0.001 | 1 | 0.001 |
|  | B | 1 | 1 | 1 | 1 | **B** | 0.001 | 1 | 1 | 1 |
|  | C | 1 | 1 | 1 | 1 | **C** | 1 | 1 | 1 | 0.001 |
|  | E | 0.001 | 1 | 1 | 1 | **E** | 0.001 | 1 | 0.001 | 1 |
| 60--65.2 | A | 1 | 1 | 0.75 | 0.001 |  |  |  |  |  |
|  | B | 1 | 1 | 1 | 0.5 |  |  |  |  |  |
|  | C | 0.75 | 1 | 1 | 1 |  |  |  |  |  |
|  | E | 0.001 | 0.5 | 1 | 1 |  |  |  |  |  |

Notes: A: eastern Asia; B: eastern North America including Mexico; C: western North America; E: Europe. Probabilities are arbitrarily determined based on geological evidence from Tiffney & Manchester (2001) and Graham (2018), regarding the availabilities of the Bering land bridge connecting A and C, North Atlantic bridge connecting B and E, the barriers between A and E from Turgai Strait, North American epicontinental seaway and North American Rocky Mountains for B and C. Long distance dispersal was indicated by 0.001.

**Table S3. Number of reads detected in each sample from RAD-seq sequencing.**

| ***Castanea* (30)** | | | | |  | ***Hamamelis* (28)** | | | | |  |
| --- | --- | --- | --- | --- | --- | --- | --- | --- | --- | --- | --- |
| **Samples** | **Raw Reads #** | **Clean Reads #** | **Samples** | **Raw Reads #** | **Clean Reads #** | **Samples** | **Raw Reads #** | **Clean Reads #** | **Samples** | **Raw Reads #** | **Clean Reads #** |
| **CC_31** | 502625 | 502535 | **CSE_46** | 1788192 | 1787916 | **HJ_61** | 272330 | 272272 | **FOM_76** | 314301 | 314217 |
| **CC_32** | 1177959 | 1177785 | **CSE_47** | 80886 | 80875 | **HJ_62** | 3016285 | 3015568 | **HVE_77** | 394091 | 394026 |
| **CC_33** | 251423 | 251358 | **CP_48** | 84668 | 84654 | **HJ_63** | 114627 | 114603 | **HME_78** | 561577 | 561435 |
| **CD_34** | 1486011 | 1485760 | **CP_49** | 1832424 | 1832147 | **HME_64** | 187402 | 187360 | **HVI_79** | 1900954 | 1900655 |
| **CP_35** | 71229 | 71219 | **CP_50** | 1024235 | 1023973 | **HVI_65** | 1235353 | 1235085 | **HVI_80** | 502672 | 502557 |
| **CD_36** | 23077 | 23069 | **CP_51** | 251265 | 251226 | **HMO_66** | 200574 | 200536 | **HVI_81** | 723155 | 723048 |
| **CD_37** | 778018 | 777834 | **CPO_52** | 569283 | 569155 | **HMO_67** | 413779 | 413696 | **HVI_82** | 283754 | 283691 |
| **CSE_38** | 144326 | 144298 | **CPO_53** | 22532 | 22523 | **HMO_68** | 601369 | 601209 | **HVI_83** | 1586403 | 1586051 |
| **CSE_39** | 102443 | 102419 | **CPO_54** | 410325 | 410264 | **HMO_69** | 597945 | 597806 | **HVI_84** | 568 | 568 |
| **CH_40** | 208297 | 208253 | **CM_55** | 479731 | 479646 | **HMO_70** | 1010389 | 1010181 | **HI_85** | 184966 | 184935 |
| **CH_41** | 50859 | 50854 | **CM_56** | 445036 | 444966 | **HO_71** | 22305 | 22303 | **HVE_86** | 1193903 | 1193643 |
| **CM_42** | 924523 | 924297 | **CS_57** | 462925 | 462806 | **HO_72** | 497472 | 497368 | **FOG_87** | 602597 | 602445 |
| **CM_43** | 1480119 | 1479890 | **CS_58** | 411935 | 411838 | **HO_73** | 148458 | 148419 | **PJ_88** | 438669 | 438577 |
| **CM_44** | 2033411 | 2032928 | **FAG_59** | 1800428 | 1800138 | **HO_74** | 1439530 | 1439303 |  |  |  |
| **CM_45** | 871312 | 871134 | **QC_60** | 16395 | 16391 | **HVE_75** | 1120947 | 1120674 |  |  |  |

Note: Green color highlights the numbers of filtered reads in the range of 10000 - 50000. Yellow color highlights the number of filtered reads less than 10000. Average of raw reads in Castanea and Hamamelis are 659529.73 and 698799.11, respectively; while the average of clean reads in Castanea and Hamamelis are 659405.03 and 698651.11, respectively.

**Table S4 Number of loci and missing data in the seven matrices of RAD-seq data in *Castanea* and *Hamamelis*.**

| **Genus** | **Matrix** | **# of taxa** | **# of sequences containing < 30% Ns** | **# of loci** | **# of SNP** | **# of unlinked SNP** | **# of Ns in unlinked SNP matrix** |
| --- | --- | --- | --- | --- | --- | --- | --- |
| ***Castanea*** | M20 | 30 | 5 | 3597 | 19058 | 3338 | 54213 (54.14%) |
|  | M30 | 30 | 9 | 2544 | 14920 | 2458 | 33892 (45.96%) |
|  | M40 | 30 | 18 | 1818 | 11380 | 1773 | 20103 (37.79%) |
|  | M50 | 30 | 22 | 1290 | 8500 | 1269 | 11489 (30.18%) |
|  | M60 | 30 | 23 | 977 | 6726 | 963 | 7141 (24.72%) |
|  | M70 | 30 | 23 | 765 | 5341 | 755 | 4862 (21.47%) |
|  | M80 | 30 | 24 | 415 | 2885 | 407 | 2133 (13.47%) |
| ***Hamamelis*** | M20 | 28 | 3 | 3935 | 21656 | 3538 | 53410 (53.91%) |
|  | M30 | 28 | 9 | 2712 | 16673 | 2534 | 31188 (43.96%) |
|  | M40 | 28 | 14 | 2086 | 13412 | 1971 | 20450 (37.06%) |
|  | M50 | 28 | 17 | 1595 | 10559 | 1519 | 13210 (31.06%) |
|  | M60 | 28 | 19 | 1123 | 7697 | 1081 | 7421 (24.52%) |
|  | M70 | 28 | 23 | 757 | 5232 | 735 | 3942 (19.15%) |
|  | M80 | 28 | 25 | 395 | 2808 | 384 | 1502 (13.97%) |

**Table S5. Divergence time comparisons among RAD-seq data and Hyb-Seq matrices.**

| ***Castanea*** | | | | | | | | |
| --- | --- | --- | --- | --- | --- | --- | --- | --- |
| **Comparisons** | | | **Node1 age (Ma)** | **Node2 age (Ma)** | **Node3 age (Ma)** | **Node4 age (Ma)** | **Node5 age (Ma)** | **Node6 age (Mya)** |
| **RAD-seq (control)** | | | 23[21.4,25.8] | NA | 11.8[10.7,13.3] | 18.3[16.8,20.4] | 13.1[11.9,14.6] | 8.2[7.4,9.3] |
| ***consensus*** | **untrimmed_orthologs** | **supercontig** | 26.7[22.9,31.2] | 23.5[19.9,27.2] | 18.9[16.1,22.1] | 23.4[19.9,27.2] | 19.6[16.7,22.9] | 17.7[15.1,20.8] |
|  | **well-trimmed_orthologs** | **supercontig** | 19.8[16.9,23.5] | 16.5[13.9,19.5] | 12.3[10.4,14.8] | 16.3[13.7,19.3] | 13.1[11.0,15.5] | 10.7[9.0,12.8] |
| ***degenerated* (BWA19)** | **well-trimmed_orthologs** | **exon** | 21.0[19.4,23.5] | 15.1[13.7,16.9] | 11.4[10.2,12.8] | 15.4[14.1,17.3] | 11.2[10.2,12.6] | 8.5[7.6,9.7] |
|  | **untrimmed_orthologs** | **supercontig** | 21.8[18.4,25.4] | 17.7[15.1,21.0] | 13.2[11.2,15.5] | 17.8[15.1,20.9] | 14.1[12.0,16.7] | 11.5[9.8,13.5] |
|  | **auto-trimmed_orthologs** | **supercontig** | 21.5[17.3,26.2] | 18.0[14.9,22.3] | 12.9[10.4,16.1] | 17.7[14.1,21.6] | 14.4[11.5,17.7] | 11.5[9.2,14.2] |
|  | **well-trimmed_orthologs** | **supercontig** | 17.5[14.4, 21.3] | 12.9[10.5,15.5] | 8.7[6.9,10.6] | 13.1[10.6,15.9] | 9.7[7.8,11.8] | 6.7[5.3,8.2] |
| ***degenerated* (BWA100)** | **well-trimmed_orthologs** | **supercontig** | 17.9[14.3,21.8] | 13.6[10.9,16.7] | 9.3[7.4,11.6] | 13.8[11.0,17.0] | 10.4[8.2,12.8] | 7.6[6.0,9.4] |
|  | **well-trimmed_paralogs** | **supercontig** | 19.1[16.1,22.8] | 16.7[13.8,20.0] | 11.1[9.4,13.2] | 16.1[13.5,19.2] | 13.0[10.8,15.5] | 11.2[9.4,13.4] |
| **Combined RAD-Hyb-Seq data** | | | 18.3[14.9,22.4] | 14.0[11.1,17.0] | 9.6[7.6,11.9] | 14.1[11.2,17.1] | 10.6[8.4,12.9] | 7.6[6.0,9.4] |

Notes: Node1 means crown node of *Castanea*; Node2 means the node connecting ENA clade and *C. sativa*; Node3 means the crown node of ENA clade; Node4 means crown node of EA clade; Node 5 means the node including all EA clade species except *C. henryi*; Node6 include *C. crenata* and *C. seguinii*.

| ***Hamamelis*** | | | | | | | |
| --- | --- | --- | --- | --- | --- | --- | --- |
| **Comparisons** | | | **Node1 age (Ma)** | **Node2 age (Ma)** | **Node3 age (Ma)** | **Node4 age (Ma)** | **Topology** |
| **RAD-seq (control)** | | | 25.8[22.4,29.8] | 21.9[19.0,25.3] | 7.1[6.1,8.4] | 5.4[4.5,6.3] | 1 |
| ***consensus*** | **untrimmed** | **supercontig** | 33.4[30.0,38.2] | 29.6[26.5,33.9] | 19.7[17.5,22.6] | 16.2[14.3,18.6] | 3 |
|  | **well-trimmed** | **supercontig** | 28.9[26.1,32.7] | 25.4[22.9,28.8] | 13.6[12.1,15.4] | 10.9[9.8,12.5] | 1 |
| ***degenerated* (BWA19)** | **well-trimmed_orthologs** | **exon** | 25.0[22.4,28.4] | 19.9[17.5,22.9] | 10.4[9.1,12.0] | 8.5[7.3,9.8] | 1 |
|  | **untrimmed_orthologs** | **supercontig** | NA | NA | NA | NA |  |
|  | **auto-trimmed_orthologs** | **supercontig** | 28.3[23.6,33.4] | 24.9[20.7,29.5] | 12.3[10.2,14.5] | 9.9[8.2,11.8] | 1 |
|  | **well-trimmed_orthologs** | **supercontig** | 26.8[22.4,31.3] | 22.4[18.8,26.0] | 9.2[7.7,10.8] | 7.2[5.9,8.4] | 1 |
| ***degenerated* (BWA100)** | **well-trimmed_orthologs** | **supercontig** | 27.7[24.0,31.6] | 23.3[20.2,26.7] | 10.4[9.0,12.1] | 8.1[7.0,9.4] | 1 |
|  | **well-trimmed_orthologs** | **supercontig** | **24.9[22.2,28.5]** | **21.8[19.1,24.9]** | **10.9[9.6,12.3]** | **8.5[7.4,9.7]** | **2** |
|  | **well-trimmed_paralogs** | **supercontig** | 27.5[18.7,39.1] | 23.5[15.7,33.3] | 15.0[9.6,21.6] | 7.5[4.4,11.7] | 2 |
| **Combined RAD-Hyb-Seq data** | | | 27.5[23.7,31.9] | 23.1[20.1,26.8] | 9.8[8.4,11.4] | 7.5[6.4,8.8] | 1 |

Notes: Node1 means crown node of *Hamamelis*; Node2 means the crown node including all species except *H. mollis* in topology 1 or except *H. japonica* in topology 3 or means the node the crown node of *H. mollis* and *H. japonica*; Node3 means the crown node of ENA clade; Node4 means crown node of all ENA clade except *H. virginiana*.
